# Blood metabolomic profiling reveals new targets in the management of psychological symptoms associated with severe alcohol use disorder

**DOI:** 10.1101/2024.02.27.582239

**Authors:** Sophie Leclercq, Hany Ahmed, Camille Amadieu, Géraldine Petit, Ville Koistinen, Quentin Leyrolle, Marie Poncin, Peter Stärkel, Eloise Kok, Pekka J. Karhunen, Philippe De Timary, Sophie Layé, Audrey M. Neyrinck, Olli Kärkkäinen, Kati Hanhineva, Nathalie M. Delzenne

## Abstract

Alcohol use disorder (AUD) is a global health problem with limited therapeutic options. The biochemical mechanisms that lead to this disorder are not yet fully understood, and in this respect, metabolomics represents a promising approach to decipher metabolic events related to AUD. The plasma metabolome contains a plethora of bioactive molecules that reflects the functional changes in host metabolism but also the impact of the gut microbiome and nutritional habits. In this study, we investigated the impact of severe AUD (sAUD), and of a three-week period of alcohol abstinence, on the blood metabolome (non-targeted LC-MS metabolomics analysis) in 96 sAUD patients hospitalized for alcohol withdrawal. We found that the plasma levels of different lipids ((lyso)phosphatidylcholines, long-chain fatty acids), short-chain fatty acids (i.e. 3-hydroxyvaleric acid) and bile acids were altered in sAUD patients. In addition, several microbial metabolites, including indole-3-propionic acid, p-cresol sulfate, hippuric acid, pyrocatechol sulfate, and metabolites belonging to xanthine class (paraxanthine, theobromine and theophylline) were sensitive to alcohol exposure and alcohol withdrawal. 3-Hydroxyvaleric acid, caffeine metabolites (theobromine, paraxanthine and theophylline) and microbial metabolites (hippuric acid and pyrocatechol sulfate) were correlated with anxiety, depression and alcohol craving. Metabolomics analysis in post-mortem samples of frontal cortex and cerebrospinal fluid of those consuming a high level of alcohol revealed that those metabolites can be found also in brain tissue. Our data allow to for the identification of neuroactive metabolites, from interactions between food components and microbiota, which may represent new targets in the management of neuropsychiatric diseases such as sAUD.

## Introduction

Alcohol use disorder (AUD) is a global health problem accounting for substantial difficulties for the individuals who consume, their related persons, and for the society. The biochemical mechanisms that lead to this disorder are not yet fully understood, and in this respect, metabolomics represents an interesting approach to decipher metabolic events related to AUD. Improving our understanding of the pathology could lead to discovering potential novel targets for therapies (1).

Alcohol consumption clearly leads to alterations of the circulating metabolome (1–4). For instance, changes in the levels of lipids (fatty acids, phosphatidylcholine, steroids) and amino acids (glutamine, tyrosine, alanine, serotonin, asparagine) are commonly observed and, interestingly, some changes in the blood metabolite profile precedes the emergence of alcohol use related diseases, such as lower levels of serotonin and asparagine (5). Metabolomics studies conducted in rodent models of alcohol exposure have mostly targeted the liver tissue or urine (6–8). In human, plasma or serum metabolomics studies have tested the effects of low/moderate/excessive alcohol intake (9) but are rarely performed in clinical populations of severe AUD (sAUD) patients, including analysis of central nervous system (CNS) tissues(10). Furthermore, the methodology could differ across studies, some using nuclear magnetic resonance (NMR) while others using more sensitive mass spectrometry (MS) coupled with liquid or gas chromatography (LC or GC), leading to the detection of different metabolites.

The human blood metabolome consists of 1) small molecules that directly represent the functional changes in host metabolism, 2) metabolites produced by intestinal micro-organisms and 3) metabolites originating from nutrition of other exogenous sources like drugs (1,11).

The aims of this study are multiple. First, we investigated the impact of sAUD on the blood metabolome by non-targeted LC-MS metabolomics analysis. Second, we investigated the impact of a short-term alcohol abstinence on the blood metabolome followed by assessing the correlations between the blood metabolome and psychological symptoms developed in sAUD patients. Last, we hypothesized that metabolites significantly correlated with depression, anxiety or alcohol craving could potentially have neuroactive properties, and therefore the presence of those neuroactive metabolites was confirmed in the central nervous system using post-mortem analysis of frontal cortex and cerebrospinal fluid of persons with a history of heavy alcohol use.

Our data bring new insights on xenobiotics- or microbial-derived neuroactive metabolites, which can represent an interesting strategy to prevent or treat psychiatric disorders such as sAUD.

## RESULTS

### Clinical characteristics of the study participants

Two cohorts of sAUD patients (ALCOHOLBIS and GUT2BRAIN) were used in this study. All patients were hospitalized for a 3-week detoxification program, and tested at two timepoints: T1 which represents the first day of alcohol withdrawal, and T2 which represents the last day of the detoxification program. Both groups of patients were similar in terms of age, gender, smoking and drinking habits and presented with high scores of depression, anxiety and alcohol craving at T1 (Table 1). These biological and psychological similarities allow us to combine both cohorts (and consequently increase sample size) and compare them to a group of heathy controls for metabolomics analysis”.

**Table 1.**
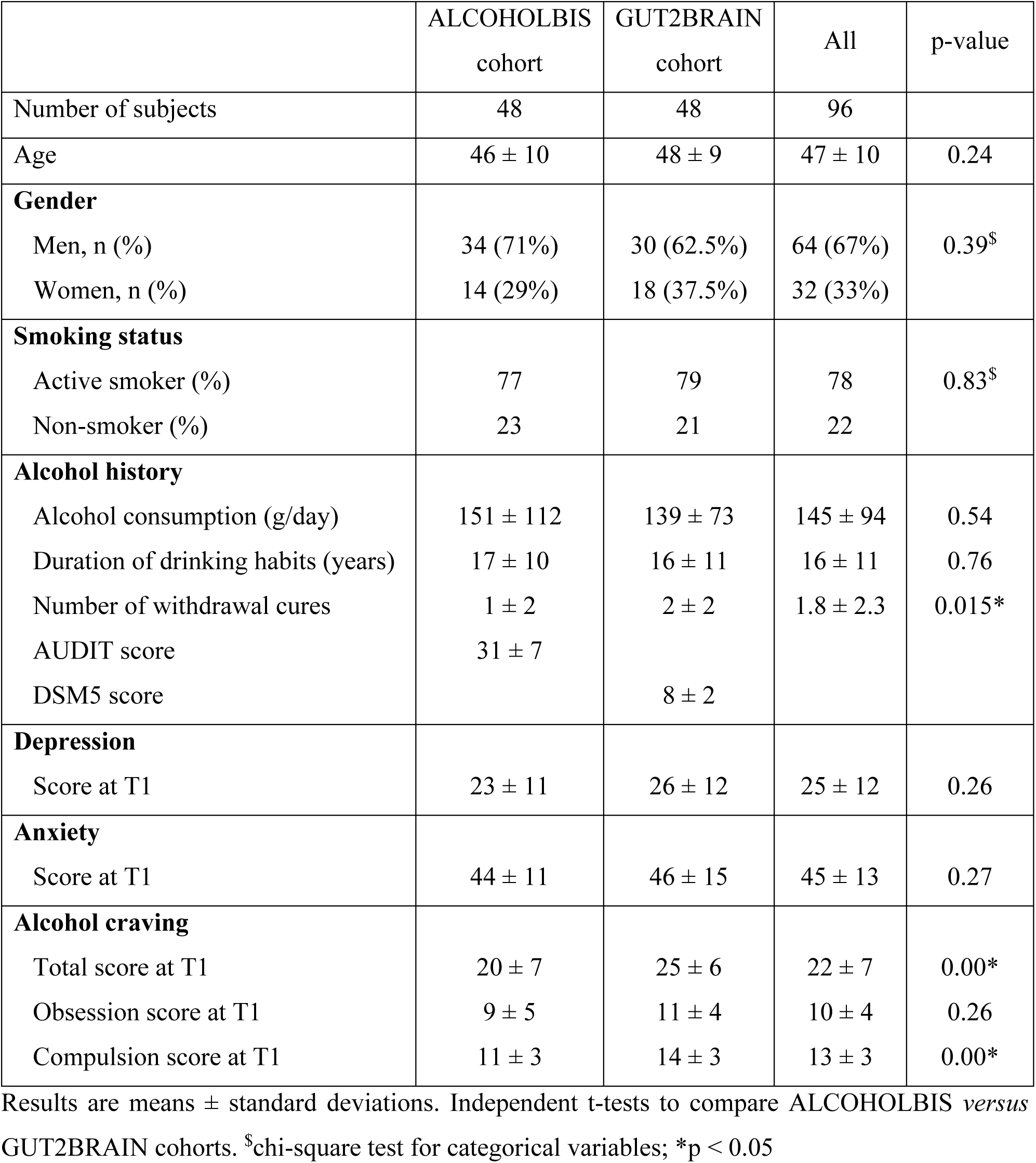
Clinical features of the study participants.

### Alterations in the plasma metabolome of sAUD patients

The metabolomics analysis allowed for sorting out a total of 11,651 features from the four analytical modes of the plasma samples. An unsupervised principal component analysis (PCA) model of the plasma metabolomic profiles between healthy controls and sAUD patients at the beginning of the withdrawal (T1) is shown in Fig. 1a. In addition, the scores plot and the performance of a supervised sparse partial least square discriminant analysis (sPLS-DA) model are shown in Figs. S1 and S2, respectively. Between healthy controls and sAUD T1, the annotated differential (Welch *t*-test *q*<0.05 and sPLS-DA variable importance in the projection (VIP) score > 2.0) metabolites included compounds from several metabolite classes as indicated in Fig. 1b and Table S3. Compared to healthy controls, the metabolic profiles of sAUD patients were characterized by an increase in long-chain fatty acids, such as 16:1 (palmitoleic acid), 18:1 (octadecenoic acid) and 22:4 (docosatetraenoic acid) fatty acids and phospholipids holding these fatty acids. In addition, several drugs (like diazepam, trazodone) and metabolites with steroid backbone such glycinated bile acids (glycohyodeoxycholic acid and glycochenodeoxycholic acid), steroid hormones and acylcarnitines were increased in the sAUD group. We also observed a significant increase in vitamin B6 metabolite, 4-pyridoxic acid, nicotine metabolite cotinine, a hydroxy fatty acid 3-hydroxyvaleric acid and stress hormone cortisol. However, lysophosphatidylcholines (LPCs) holding a saturated odd-chain (e.g. LPC 15:0 and LPC 17:0), polyunsaturated 18-carbon fatty acid or an ether bond (O-) containing lipid were consistently decreased in sAUD compared to controls (Fig. 1b). Further, we also showed a decrease in furan fatty acids 3-carboxy-4-methyl-5-pentyl-2-furanpropanoic acid (3-CMPFP) and 3-carboxy-4-methyl-5-propyl-2-furanpropionic acid (CMPF), in a carotenoid compound and in several metabolites belonging to the family of xanthine. In addition, several amino-acid derived bacterial metabolites such as pipecolic acid, 3-indole propionic acid, p-cresol sulfate and hippuric acid were significantly decreased in sAUD patients compared to controls. The top-ranked metabolites in Fig. 1b remained unknown regardless of manual curation.

**Figure 1.**
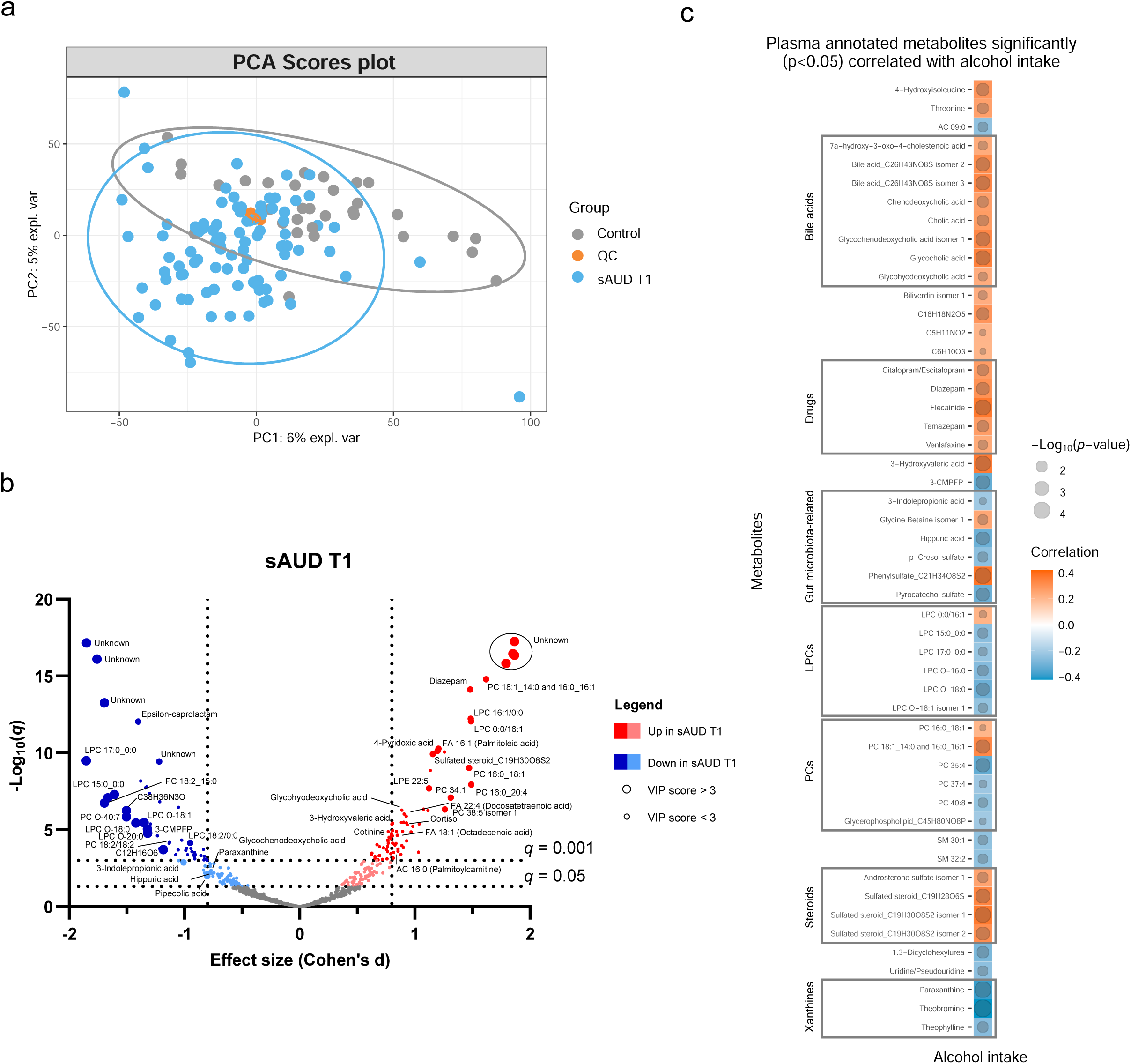
Chronic alcohol intake alters the plasma metabolome compared to healthy controls. **a** Principal component analysis score plot of the plasma metabolomic features between healthy controls and persons with sAUD at T1. QC samples are colored in orange. **b** Volcano plot depicting the effect size (Cohen’s D) and -Log10 transformed *q* values derived from Welch’s *t*- test analysis of the metabolomic features different between healthy controls and persons with sAUD at T1. Circle size represent the variable importance in projections (VIP) scores derived from the sPLS-DA model for the plasma metabolomic features between persons with sAUD at T1 and healthy controls. **c** Annotated metabolites having significant association (Spearman *p*<0.05) with alcohol intake (g/day) in persons with AUD at T1. Circle size refers to the level of significance, blue gradient color to the strength of negative while red to the strength of positive correlation coefficients. 3-CMPFP 3-carboxy-4-methyl-5-propyl-2-furanpropionic acid; AC acylcarnitine; FA fatty acid; LPC lysophosphatidylcholine; LPE lysophosphatidylethanolamine; PC phosphatidylcholine; PE phosphatidylethanolamine; SM sphingomyelin.

We then conducted a correlational analysis between blood metabolites and alcohol consumption reported by the patients. Alcohol intake was positively correlated with annotated bile acids, steroids and drugs while xanthines (paraxanthine, theobromine and theophylline), odd-chain or ether-bond LPCs and bacterial metabolite hippuric acid, p-cresol sulfate, pyrocatechol sulfate, and 3-indole propionic acid showed negative correlations (Fig. 1c).

### Alcohol withdrawal shapes the plasma metabolome

The score plot of the sPLS-DA in Fig. 2a shows a clear discrimination in plasma metabolomic profiles in the course of withdrawal in sAUD patients. The sPLS-DA model performance and the unsupervised PCA model scores plot are shown in supplementary Figs. S3 and S4, respectively. Annotated metabolites discriminating sAUD groups (paired *t*-test *q*<0.05, sPLS-DA VIP score > 2.0) before (T1) and after (T2) the 3-week withdrawal period included metabolites from a range of chemical classes (Fig. 2b, Table S3). Apart from the metabolites belonging to the class of bilirubins, the levels of identified metabolites that were significantly changed upon alcohol withdrawal (the major ones being indoles, acylcarnitines, glycerophospholipids and xanthines) came back towards to the levels observed in controls (Table S3). In the course of alcohol abstinence, we noted a significant decrease in 16-chain acylcarnitines, LPCs with 16- or 18-chain fatty acid tails excluding LPCs with ether bonds, retinol, tryptophyl-phenylalanine dipeptide and 3-hydroxyvaleric acid (Fig. 2b). On the contrary, LPCs with odd-chain fatty acid tails or ether bonds show a significant increase along with tryptophan derivatives 3-indoleacetic acid and 3-indolepropanoic acid and metabolites of the xanthine family such as theophylline, paraxanthine, theobromine and trigonelline during alcohol abstinence. The changes in metabolites belonging to the xanthine family during alcohol withdrawal could be explained by the changes in dietary intake of coffee, tea and chocolate (see Fig S5).

**Figure 2.**
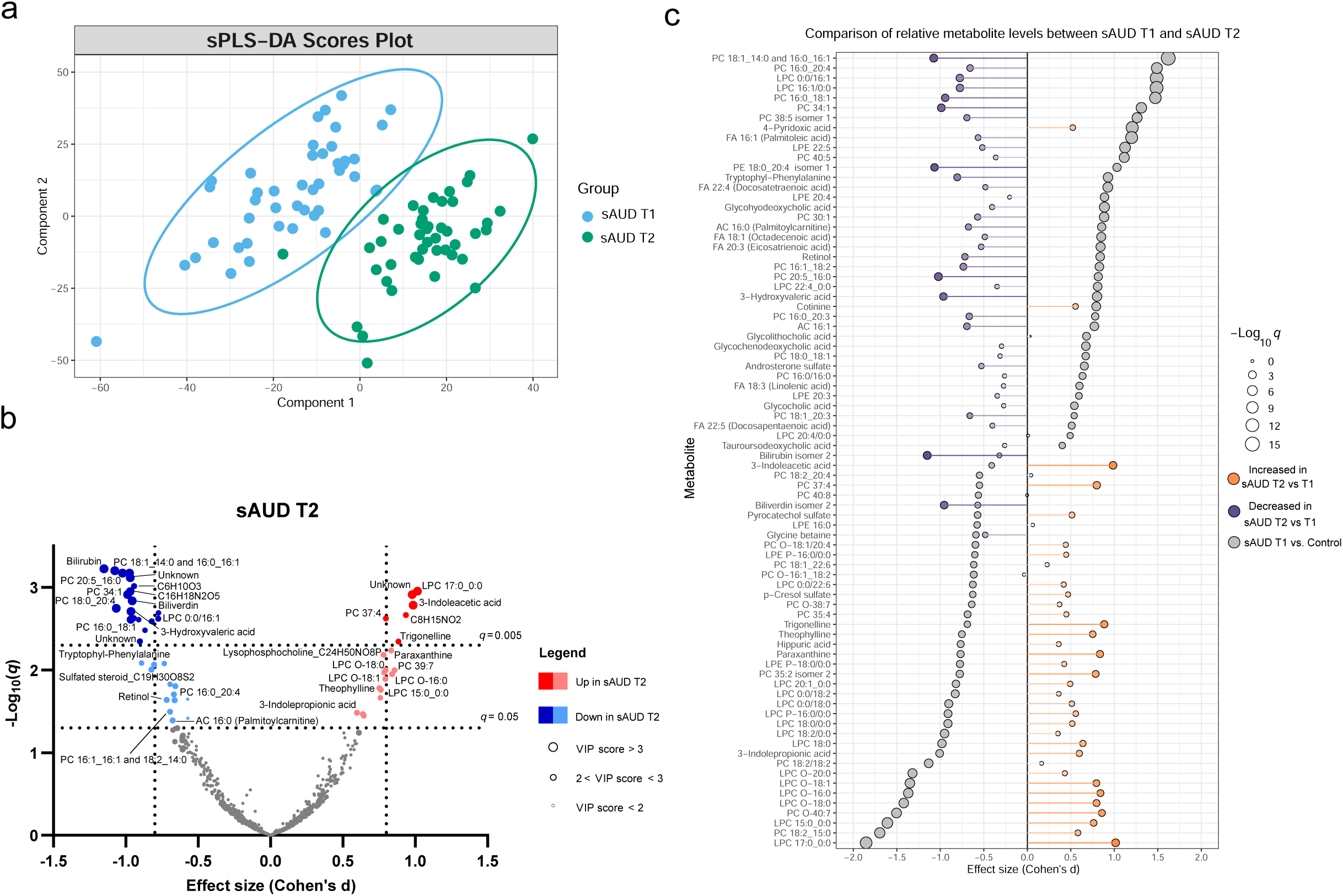
3-week alcohol withdrawal shapes the plasma metabolome. **a** Scores plot of the sparse partial least square discriminant analysis (sPLS-DA) of the plasma metabolomes at the start (T1) and end (T2) of the alcohol withdrawal in persons with sAUD. **b** Volcano plot depicting the effect size (Cohen’s D) and -Log10 transformed *q* values derived from paired *t*-test analysis of the metabolomic features different between in persons with sAUD at T1 and T2. Circle size represent the variable importance in projections (VIP) scores for the plasma metabolomic features in the sPLS-DA model of persons with sAUD at T1 and T2. **c** Lollipop plot of the effect size (Cohen’s D) and -Log10 transformed *q* values of the altered annotated metabolites between sAUD T1 and healthy controls as well as sAUD T1 and sAUD T2. Circle size refers to the level of significance, grey color to the comparison between controls and sAUD T1, orange color to relative increase while violet to relative decrease towards the end of alcohol withdrawal (T2). AC acylcarnitine; FA fatty acid; LPC lysophosphatidylcholine; LPE lysophosphatidylethanolamine; PC phosphatidylcholine.

Overall, Fig. 2c demonstrates that a number of identified metabolites altered in sAUD patients relative to control are affected by alcohol withdrawal. Apart from 4-pyridoxic acid, cotinine, and heme metabolites bilirubin and biliverdin, the shifts observed in the selected metabolites are generally in the opposite direction as compared to the baseline.

### Correlations between blood metabolites and psychological symptoms

Correlation analysis shows that, at T1, 96 annotated features were significantly (p<0.05) correlated with psychological scores of anxiety, depression and alcohol craving (with sub- scores of obsession and compulsion) (Fig 3). Annotated bile acids, drugs, lysophosphatidylethanolamines (LPE), fatty acids, LPCs apart from LPCs with an ether-bond or 15:0 or 17:0 fatty acid tail were consistently positively correlated with psychological symptoms, and more particularly with the compulsive component of alcohol craving. Members of the xanthine family, pyrocatechol sulfate, a pentose sugar (mannose/fructose), hippuric acid, 1-methyl-pyridone-carboxamide, acylcarnitines with maximum of 10 carbons, creatinine and kynurenine were negatively correlated with psychological symptoms. Within the acylcarnitine metabolite class, an interesting pattern was observed, as the shorter chain-length acylcarnitines were consistently negatively correlated with the psychological parameters, and the long-chain ones demonstrated positive correlation.

**Figure 3.**
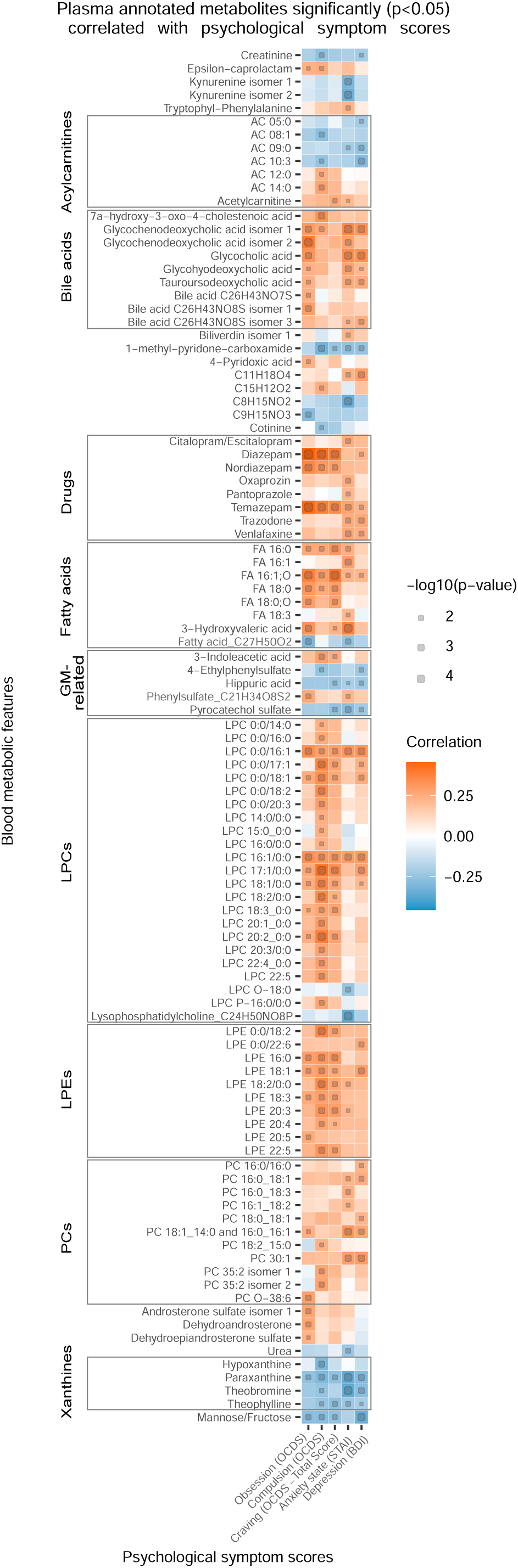
Plasma metabolites associated with psychological symptom scores. Heatmap of the annotated metabolites having significant (p<0.05) Spearman correlation with one or more psychological symptom score of obsession, compulsion, alcohol craving, anxiety state or depression. Circle size refers to the level of significance, blue gradient color to the strength of negative while red to the strength of positive correlation coefficients. AC acylcarnitine; FA fatty acid; GM gut microbiota; LPC lysophosphatidylcholine; LPE lysophosphatidylethanolamine; PC phosphatidylcholine.

### Heavy alcohol use-related alterations in the brain metabolome

Based on the annotated significantly altered plasma metabolites, we conducted a targeted search in a metabolomics dataset consisting of cerebrospinal fluid (CSF) and frontal cortex samples collected from deceased individuals with a history of heavy alcohol use and control individuals. 79 and 74 of the annotated plasma metabolites were identified from the CSF and frontal cortex, respectively (Fig. 4a, Table S3). We looked specifically at metabolites significantly correlated with at least one psychological symptom. 3-Hydroxyvaleric acid, cotinine, theobromine and paraxanthine were indeed present in the central nervous system (CNS) and significantly (Welch *t*-test p<0.05) different between heavy alcohol use and control groups in both frontal cortex (Fig 4b) and CSF (Fig 4c). Additional significantly altered metabolites found only in the frontal cortex were LPE 20:3, nordiazepam, PC 16:0_18:1 and urea (Fig. 4b). In the CSF, the independent differential metabolites were FA 16:1 (palmitoleic acid), hippuric acid, LPCs 16:0 and 18:1, LPEs 16:0, 18:1 and 20:4 and pyrocatechol sulfate (Fig. 4c).

**Figure 4.**
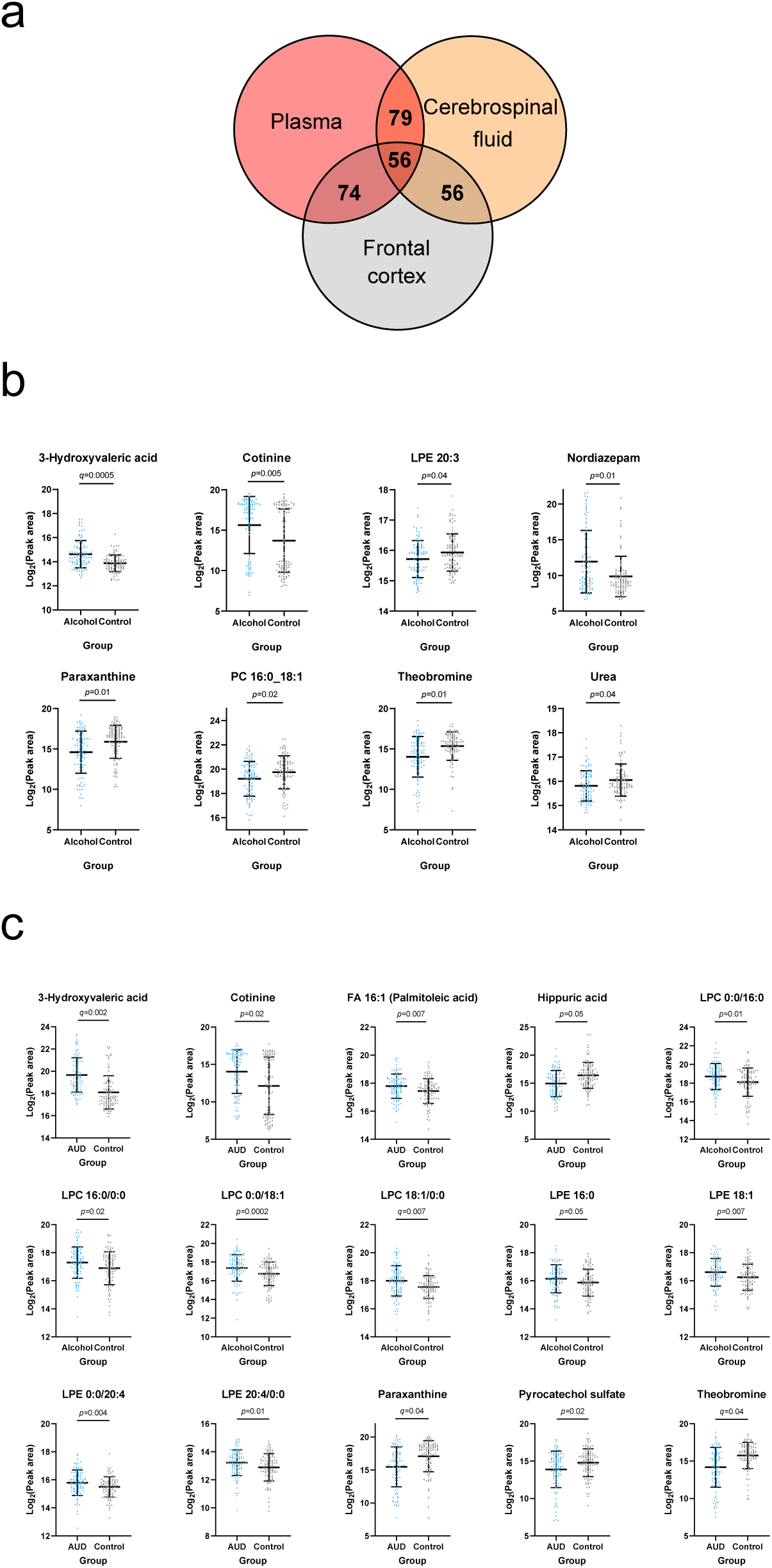
Overlapping metabolites within the plasma and brain metabolome. **a** Venn diagram of the annotated plasma metabolites that are also present in the frontal cortex and cerebrospinal fluid metabolome. **b** Significantly altered (*p*<0.05) frontal cortex metabolites with a corresponding plasma metabolite showing an association with one or more psychological symptom score. **c** Significantly altered (*p*<0.05) cerebrospinal fluid metabolite with a corresponding plasma metabolite showing an association with one or more psychological symptom score. Data expressed as mean ± SD with individual values shown. Statistical values derived from Welch’s *t-*test comparing metabolomic features between control and alcohol groups.

## DISCUSSION

The circulating metabolome reflects the crosstalk between nutrition, microbiome and host metabolism (12), with diet and microbiome being the strongest determinants of the human blood metabolome (11,13). In this study, we showed the impact of sAUD, and the impact of a short-term abstinence, on the blood metabolome. We analyzed the correlations between blood metabolites and psychological symptoms, as well as the presence of identified metabolites in the CNS of individuals considered as heavy alcohol drinkers.

### Impact of sAUD on the blood metabolome

In 2019, a review summarized the results obtained from 23 studies that have used a metabolomics approach for measuring changes in metabolite profiles in relation to alcohol use (1). Changes in lipids have been highlighted as the most consistent changes across studies.

Lipids are an integral part of cell membranes and signaling molecules in the body. PCs and LPCs have been suggested to form a new class of biomarkers for alcohol consumption (2). For instance, in our study palmitoleic acid (FA 16:1) was largely increased in sAUD patients compared to controls, and in other studies, this metabolite has likewise been significantly associated with alcohol consumption (14,15). Another clear observation in our study was the lower level of odd-chain lipids in sAUD patients. Since the lipids containing FA 15:0 and FA 17:0 have been suggested to be products from bacterial metabolism (16,17), the existence of gut dysbiosis in sAUD patients could explain the lower abundance of LPC 17:0 and LPC 15:0 (18).

We found that some bile acids, sulphated steroids and 3-hydroxyvaleric acid were positively associated with the amount of alcohol consumed. Sulphated steroids and hydroxyvalerate have previously been associated with alcohol intake (19,20). Metabolites belonging to the xanthine family (theobromine, theophylline, paraxanthine) and microbial metabolites (hippuric acid, indole-3-propionic acid, p-cresol sulfate, pyrocatechol sulfate) were negatively correlated with alcohol consumption. Altogether, these results suggest that these metabolites are sensitive to alcohol exposure. Interestingly, these metabolites were also correlated with the severity of the psychological symptoms suggesting that they could play a role in the symptomatology of alcohol use disorder.

### Effect of short-term alcohol abstinence on the blood metabolome

Since alcohol consumption is known to influence lipid metabolism, it was expected that a short-term alcohol abstinence could reverse or ameliorate lipidomic alterations. Indeed, we found that some phospholipids that were increased in sAUD patients at baseline, such as PC 16:0_18:1, PC18:1_14:0 and 16:0_16:1 as well as LPC 16:1, were downregulated during alcohol withdrawal to reach the levels of healthy controls after detox. On the other hand, LPC 15:0 and LPC 17:0 that were decreased in sAUD patients at baseline, increased during alcohol withdrawal, but did not reach the levels of controls at the end of detox.

The metabolite that contributed the most to the differences observed with alcohol detoxification was bilirubin. While bilirubin was not statistically higher in sAUD patients at baseline vs controls, we observed a significant reduction of this metabolite after a 3-week alcohol withdrawal. In a previous study, serum bilirubin was found to be associated with alcohol consumption, cigarette smoking and coffee consumption (21). Interestingly, the caffeine metabolites belonging to the xanthine family such as paraxanthine, theophylline and theobromine that were decreased at baseline in sAUD patients compared to controls, increased significantly during alcohol withdrawal to reach the levels of healthy controls. Changes in dietary intake of coffee, tea and chocolate during alcohol withdrawal could explain these results Also, the bacterial metabolites indole derivatives such as 3-indolepropionic acid and 3-indoleacetic acid increased during alcohol withdrawal to reach the levels of healthy controls. Intriguingly, 3-hydroxyvaleric acid significantly decreased during alcohol withdrawal and was found to be lower than healthy controls at the end of detoxification period.

Metabolites that remained significantly higher in sAUD patients at the end of detoxification compared to controls are stress hormone cortisol, palmitoleic acid (FA 16:1), some bile acids, some drugs (diazepam, trazodone), vitB6 metabolite (4-pyridoxic acid, which is likely due to the fact that patients received vitamin B supplement during their hospital stay) and cotinine (nicotine metabolite that reveals the higher proportion of smokers in sAUD patients compared to controls).

### Identification of blood metabolites with potential neuroactive properties

The metabolites belonging to the xanthine family (theobromine, paraxanthine and theophylline) are metabolites of caffeine produced upon cytochrome P450 dependent oxidation in the liver. They were all decreased in the blood of sAUD patients at baseline and were negatively correlated with alcohol intake, alcohol craving, depression and anxiety. The decrease in caffeine metabolites has previously been described in the urine of AUD patients, that is linked to the increasing severity of alcoholic liver disease (22).

Theobromine is the principle alkaloid found in cocoa beans and is responsible for the bitter taste of chocolate. It is known for its mood improving effect (23). Like caffeine, theobromine is an inhibitor of brain adenosine receptors and phosphodiesterase. A study in rats showed that the antagonist of A2a adenosine receptor produced a reduction of ethanol reinforcement (24), suggesting adenosine receptor as a potential target for the treatment of alcohol abuse. In a randomized, double-blind, placebo-controlled trial, the phosphodiesterase inhibitor pentoxifylline associated with escitalopram showed greater reduction of depression scores compared to patients receiving escitalopram alone (25). In another study, Apremilast which is also a phosphodiesterase inhibitor, suppressed excessive alcohol drinking in AUD patients (26). Paraxanthine has a psychostimulant effect and can modulate dopamine release in the striatum

(27). Interestingly, in 2017 a systematic review indicated that consumption of coffee, tea and cocoa could have protective effects against depression (28).

Lipids, and mostly LPCs (except ether LPC derivatives) and LPEs were significantly and positively correlated with the compulsive component of alcohol craving. LPCs are secreted by the liver and are actively transported via the blood-brain barrier (BBB) and have been associated with pro-inflammatory events (29). LPCs are also precursors of brain lysophosphatidic acid (LPA), which regulates glutamatergic transmission and cortical excitability within the CNS. Recently, LPA has been shown to induce hyperphagia following food restriction and this effect was dependent on hypothalamic agouti-related peptide (AgRP) neurons (30). AgRP neurons have also been implicated in circuitry controlling non-feeding behavior, including those associated with reward, anxiety and compulsive disorders, more particularly in anorexia nervosa (31). Therefore, we hypothesize that the positive correlation between peripheral LPC and compulsion for alcohol drinking found in sAUD patients who have just been deprived of alcohol could be mediated by the effect of LPA on AgRP neurons. Consistent with that, postmortem brain tissues from patients consuming a high intake of alcohol showed increased levels of many LPCs (10).

Circulating bile acids can reach the brain by crossing the BBB, either by simple diffusion or active transport. Some bile acids show neuroprotective effects (32) while others are rather neurotoxic (33). In Alzheimer disease patients, the levels of glycochenodeoxycholic acid was associated with worse cognition (34). In our study, both primary (glycochenodeoxycholic acid and glycocholic acid) and secondary (glycohyodeoxycholic acid, tauroursodeoxycholic acid) bile acids were positively correlated with depression and anxiety in sAUD patients.

3-Hydroxyvaleric acid, also called β-hydroxypentanoate, was significantly and positively correlated with anxiety and alcohol craving. This metabolite is formed from odd carbon fatty acids in the liver and can reach the brain. 3-hydroxyvaleric acid is a C5-ketone body and is a precursor of propionyl-CoA that refills intermediates of citric acid cycle and is useful for alternative energy fuel in the brain (35,36).

Other co-metabolites, i.e. produced by the gut microbiota and then processed by the liver, were negatively correlated with all psychological symptoms. Pyrocatechol sulfate is a phenolic compound derived from the gut microbiota, present in the CSF of mice, and implicated in synapse formation and fear extinction learning (37). In Parkinson disease patients, the plasma level of pyrocatechol sulfate is decreased compared to controls (38). In our study, blood pyrocatechol sulfate was significantly and negatively correlated with all psychological symptoms of sAUD patients (i.e. anxiety, depression and craving) suggesting a neuroprotective role of this metabolite. Interestingly, 4-ethylphenylsulfate, another gut-derived metabolite linked with neurodevelopment abnormalities, autism and anxiety behavior in mice (39–41) showed negative correlations with depression and the compulsive component of alcohol craving.

Hippuric acid, the glycine conjugate of benzoic acid has long been associated with the microbial degradation of specific dietary components, including polyphenolic compounds (like chlorogenic acid and catechin) found in fruits, vegetables, coffee and tea (42). Hippuric acid is indeed a host-microbe cometabolite (43). It is synthesized in the liver and in the renal cortex from the microbial metabolite benzoate. The plasma concentration of hippuric acid has been shown to be 17-fold higher in conventional mice compared with their germ-free counterparts suggesting a substantial contribution of the gut microbiota in its production (44). We showed that blood hippurate levels, that correlated negatively with anxiety, depression and craving, were decreased in sAUD patients, as shown in ethanol-treated mice (6) and humans characterized by high alcohol intake and those with major depression (45,46). Urinary hippurate excretion is also decreased in depression, schizophrenia and autism spectrum disorders patients (42). A recent Mendelian randomization study including > 13,000 individuals from five European cohorts characterized for depression suggested that low hippuric acid levels in the circulation is part of the causal pathway leading to depression (47), which was consistent with a significant decrease of the dietary sources of hippuric acid including fresh fruits and vegetables in depressed patients (47).

Another way to support the neuroactive effects of the blood metabolites that are correlated with one or several psychological factors is to demonstrate their presence in the brain. We therefore conducted a targeted search in a database of post-mortem frontal cortex and CSF metabolomics analysis (10) and found that 3-hydrovaleric acid, caffeine metabolites (theobromine, paraxanthine and theophylline) and microbial metabolites (hippuric acid and pyrocatechol sulfate) that were correlated with anxiety, depression and alcohol craving in our plasma cohort were also present in the brain and in CSF, and the direction of their changes in the plasma (increase or decrease) mimicked changes in the central nervous system.

### Advantages and limitations of the study

Most of the studies assessing the impact of alcohol consumption on the blood metabolome were cross-sectional, and only included male participants (45). Here, we reported longitudinal data to assess the impact of a short-term alcohol abstinence on the blood metabolome, both in male and female AUD patients. In the study of Zhu et al (9), the AUD patients recruited were alcohol abstinent, but for various periods of abstinence. To avoid the bias of abstinence duration, our patients were enrolled in a rigorous and standardized manner, within 24 hours after the last drink. Furthermore, alcohol consumption was carefully evaluated with the time line follow back method, which allows precise calculation of the amount (and type) of alcohol consumed (48).

This study also presents some limitations. First, the metabolomics analysis was conducted with LC-MS while some important molecules, like lipoproteins, could have been measured with NMR-based methods. Combining NMR and MS-based methods could have covered a wider spectrum of metabolites. However, the non-targeted metabolic profiling with two different chromatographic methods and ionization polarities covers a wide range of metabolites ideal for our discovery-based approach. Large studies are usually required in metabolomics to observe small and medium size changes. Here, we included only s96 AUD patients, but they were all well characterized and received standardized therapies (for instance, vitB supplementation) during alcohol withdrawal.

The selection of the control group is always challenging in alcohol research. Here, the healthy subjects were matched for sex, age and BMI but not for smoking status or nutritional intake. Alcohol addiction is a major cause of malnutrition in developed countries and tobacco smoking is more prevalent in alcohol users compared to healthy subjects. These two main confounding factors, although being an integral part of the alcoholic pathology, are known to influence the blood metabolome (49–51). Furthermore, another limitation is that the control group was tested only once, while the sAUD patients were tested twice (T1 and T2). This means that we do not take into consideration the intra-personal variability of the metabolomics data when interpreting the results of alcohol withdrawal effects.

## Conclusion

LC-MS metabolomics plasma analysis allowed for the identification of metabolites that were clearly linked to alcohol consumption, and reflected changes in metabolism, alterations of nutritional status, and gut microbial dysbiosis associated with alcohol intake. In particular the changes in lipid class involving odd-chain fatty acids and ether-bond lipids as well as compounds produced by gut microbiota seem to be the most prominent indicators of metabolic malfunction related to severe alcohol use disorder, and thus warrant further studies and targeted intervention. Also, the discovery of metabolites associated with behavioral and psychiatric traits related to sAUD were of importance, and could be considered potential new therapeutic targets in the management of sAUD, namely as adjuvants in the period of alcohol abstinence. The novelty of our work was to characterize the impact of sAUD on the blood metabolome, and the impact of a short-term alcohol abstinence in the same individuals, within a cohort that included both male and female patients. Intervention studies are needed in order to bring the proof of concept that nutritional approaches – namely the addition of specific lipids, or of nutrients modulating the gut microbiome - for example prebiotic dietary fibers - may be essential and so far underestimated components of alcohol withdrawal efficacy.

## SUBJECTS AND METHODS

### Study design and participants

A total of 96 sAUD patients hospitalized for a 3-week detoxification program in the alcohol withdrawal unit at Cliniques Universitaires Saint-Luc, Brussels, Belgium were recruited. These patients belong to two different cohorts, namely ALCOHOLBIS (patients recruited in 2015 and 2019) and GUT2BRAIN (patients recruited in 2018-2019) (Table 1). The severity of AUD was evaluated by a psychiatrist using the Diagnostic and Statistical Manual of Mental Disorders (DSM) criteria, fourth edition (DSM-IV) (ALCOHOLBIS cohort) or fifth edition (DSM-5) (GUT2BRAIN cohort). Patients evaluated with the DSM-IV received the diagnosis of “alcohol dependence”, while the patients evaluated with the DSM-5 received the diagnosis of “severe alcohol use disorder” (6 or more criteria). To simplify, we used the term “sAUD” (for severe alcohol use disorder) that includes both diagnosis (sAUD and alcohol dependence)”. Patients were eligible if they had been drinking until the day of admission to the detoxification unit or the day before, and if they also did not suffer from inflammatory bowel disease, other chronic inflammatory diseases (such as rheumatoid arthritis) or cancer, nor from metabolic disorders such as obesity (BMI > 30 kg/m2), diabetes and bariatric surgery, or severe cognitive impairment (MMSE < 24). We also excluded subjects who had taken antibiotics, probiotics, or prebiotics in the 2 months prior to enrolment and those who were taking non-steroidal anti-inflammatory drugs or glucocorticoids within 1 month of inclusion. Patients with known cirrhosis or significant liver fibrosis (≥F2) detected by Fibroscan (>7.6 kPa) on the day of admission were also excluded from the study. No other psychiatric diseases, and no other addiction (except tobacco) have been diagnosed in these patients.

sAUD patients were tested twice, on the day following their admission (T1) and on days 18–19 (T2) corresponding to the last days of the detoxification program. The patients of the GUT2BRAIN cohort were initially enrolled in a randomized, double-blind, placebo-controlled study assessing the impact of prebiotic fiber supplementation on the gut-liver-brain axis (52,53). For this reason, only biological and psychological data obtained at admission (T1), and before the beginning of the prebiotic/placebo treatment, were considered. The patients of the ALCOHOLBIS cohort did not take part in any other clinical study during the 3-week hospitalization stay. Therefore, for this cohort, data obtained at both times of alcohol withdrawal (T1 and T2) were considered. Thirty-two healthy controls (13 from the GUT2BRAIN cohort and 19 from the ALCOHOLBIS cohort) matched for age, gender and BMI with no AUD (Alcohol use disorders test [AUDIT] score <8 in males and <7 in females) were also recruited using flyers posted in Brussel’s public setting (Table S1). The inclusion/exclusion criteria were the same as for sAUD patients except for the alcohol related items. Healthy controls and sAUD patients were not matched for smoking status.

The study was approved by the “Comité d’éthique Hospitalo-facultaire Saint-Luc UCLouvain” (2017/04JUL/354 and 2014/31dec/614, identification number NCT03803709 at ClinicalTrials.gov). The study has been carried out in accordance with The Code of Ethics of the World Medical Association and followed the ethical guidelines set out in the Declaration of Helsinki. All participants provided written informed consent in compliance with the European law 2001/20/CE guidelines.

For investigating presence of the potentially neuroactive metabolites in the CNS, we used metabolomics data from frontal cortex (Broadman area 9) and CSF samples from the Tampere Sudden Death Study (TSDS) cohort, which have been described in detail elsewhere (10). TSDS was collected from forensic autopsies done in the area of the Pirkanmaa Hospital District during 2010-2015, a total of 700 subjects. Out of these we identified 97 heavy alcohol users based on autopsy reports and medical records (diagnosis of alcohol-related diseases: ICD-10 codes F10.X, G31.2, G62.1, G72.1, I42.6, K70.0-K70.4, K70.9, and K86.0, or signs of heavy alcohol use in the clinical or laboratory findings, e.g., increased levels of gamma-glutamyl transferase, mean corpuscular volume, carbohydrate-deficient transferrin). Lack of these findings was inclusion criteria for the control group (n = 100), most of whom had died due to cardiovascular diseases (Table S2). Samples were stored at -80 °C until use.

### Assessment of psychological symptoms

sAUD patients were tested for depression, anxiety and alcohol craving with self-reported questionnaires: the Beck Depression Inventory [BDI] (54), the State-Trait Anxiety Inventory (STAI form YA)(55), and the Obsessive-Compulsive Drinking Scale [OCDS](56) (see supplemental material for details)

### Biological sampling

To avoid variation due to fasting state and circadian rhythm, blood samples were collected in all participants in the morning between 8:00 and 8:30 am after an overnight fasting, at T1 and T2. Blood was drawn in tubes containing EDTA as an anticoagulant. The samples were centrifuged at 1000 g for 15 min at 4°C and the plasma was frozen at −80°C until analysis.

### Nontargeted metabolomics analysis

Plasma sample preparation and LC-MS measurement were performed as previously detailed in Klåvus et al (57) (see supplemental material for methodological details of the LC-MS analysis, peak picking and data processing).

### Statistical analysis

R software version 4.0.3. was used for statistical analyses. Multivariate analyses, namely PCA for dimension reduction and sPLS-DA for group discrimination, were conducted by ‘mixOmics’ R package v. 6.14.1 (58). For the sPLS-DA model we used a cross-validation (CV) procedure of 10-fold CV repeated 50 times. Univariate analyses were conducted by ‘notame’ R package v. 0.2.1(57). Significant features were shortlisted using Welch’s and paired t-tests. All *p*-values were corrected using the Benjamini-Hochberg false discovery rate (FDR) to calculate the *q*-value. For all tests, *p* and *q* values < 0.05 were considered statistically significant. Visualizations were created by the previously mentioned R packages and GraphPad Prism v. 8.4.2. Correlation analyses were performed at T1 using R software version 3.6.1. Spearman coefficient was calculated and p-value < 0.05 was considered statistically significant.

## ACKNOWLEDGMENTS AND DISCLOSURES

SL is a Research Associate of the Fonds de la Recherche Scientifique – FNRS. Collection of TSDS samples was supported by European Union 7th Framework Program (grant number 201668 for AtheroRemo Project, State Researc h Funding for Tampere University Hospital, Finnish Foundation for Cardiovascular Research). PDT received funding from Fondation Saint Luc. NMD is a recipient of grants from the Fonds de la Recherche Scientifique (FRS-FNRS) [PDR T.0068.19], and from the Fédération Wallonie-Bruxelles (Action de Recherche Concertée ARC18- 23/092).

OK and KH are founders of Afekta Technologies Ltd. The other authors report no financial interests or potential conflicts of interest.

## AUTHOR CONTRIBUTIONS

Conceptualized and designed the study: SL, CA, PS, PdT, NMD Performed participant visits: SL, CA, GP, MP

Collected and provided the TSDS post-mortem samples: EK, PJK

Performed data analysis, interpreted the data and drafted the manuscript: SL, HA, CA, QL, PdT, SLa, AMN, OK, KH, NMD

Performed the analysis of the metabolomics data: HA, VK, OK, KH Contributed to ethics application; SL, PS, PdT, NMD

All authors approved the final version of the manuscript.

