## supplementary figures for "Blood metabolomic profiling reveals new targets in the management of psychological symptoms associated with severe alcohol use disorder"

**Figure S1.** Scores plot of the sparse partial least square discriminant analysis (sPLS-DA) separates the plasma metabolome of healthy controls and persons with AUD at the start of the withdrawal (T1)

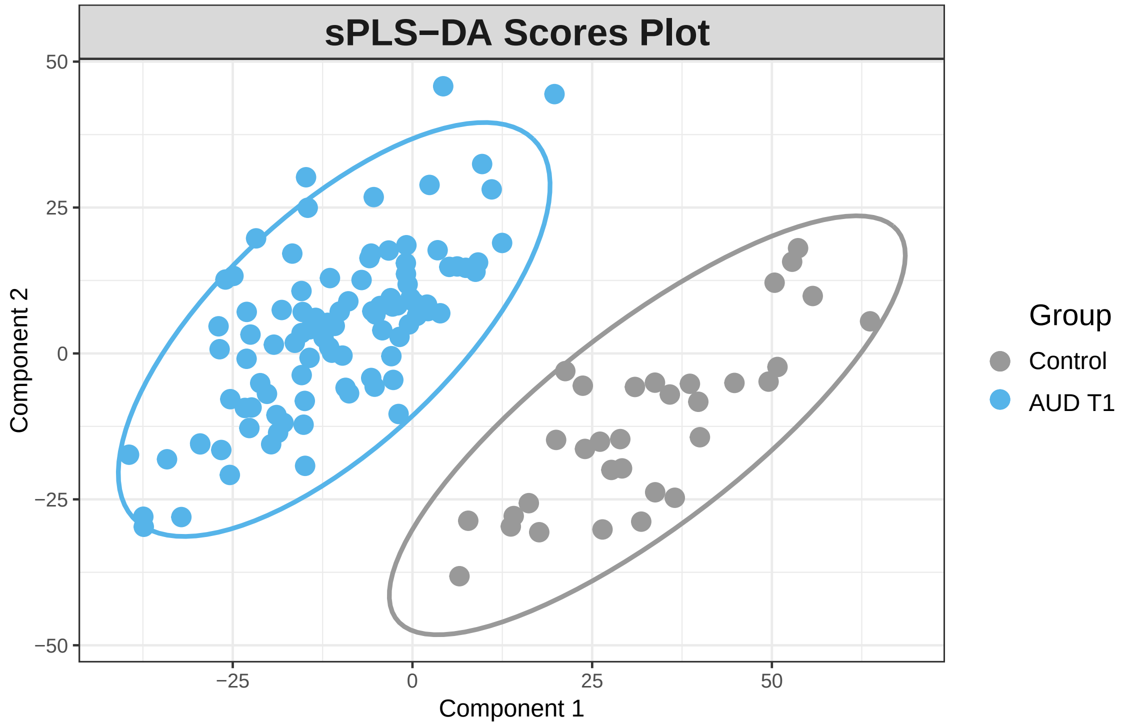

**Figure S2.** Performance of the sPLS-DA model separating plasma metabolome of healthy controls and persons with AUD at T1. Dotted line represents the optimal number of components included in the final model.

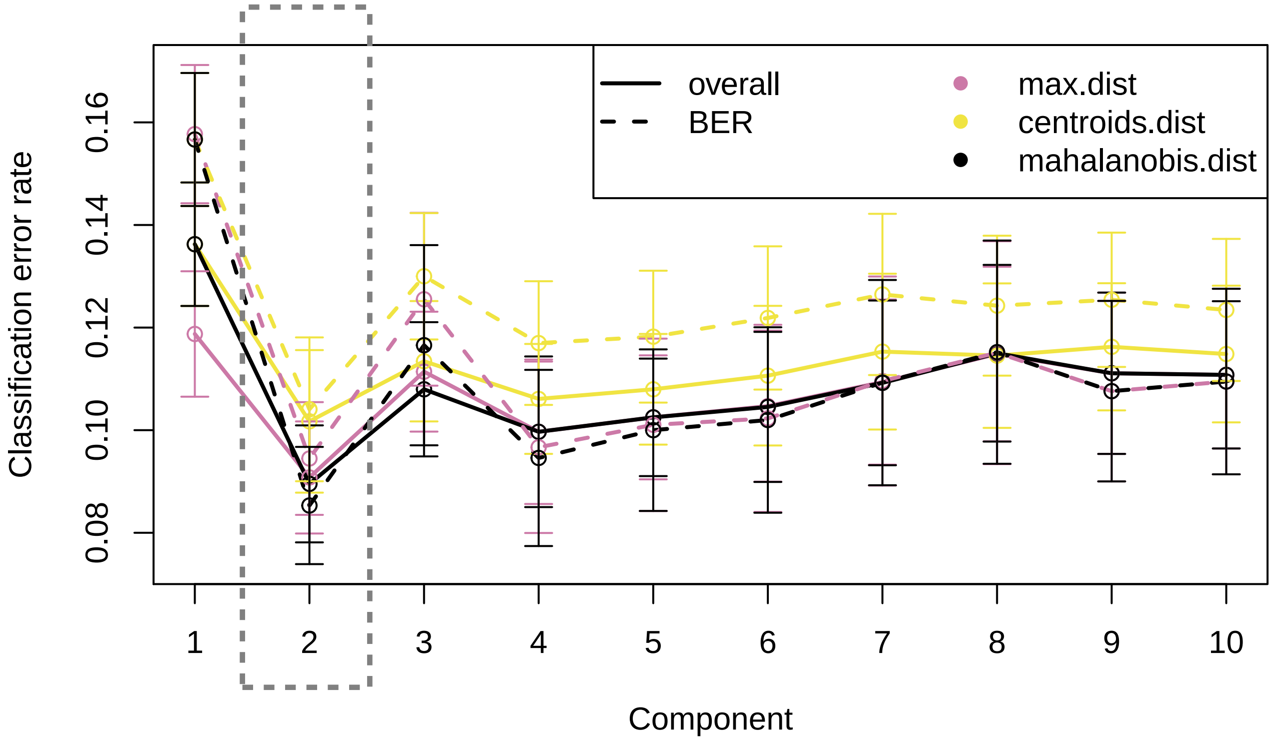

**Figure S3.** Performance of the sPLS-DA model discriminating plasma metabolome of persons with AUD between the beginning (T1) and end (T2) of alcohol withdrawal. Dotted line represents the optimal number of components included in the final model.

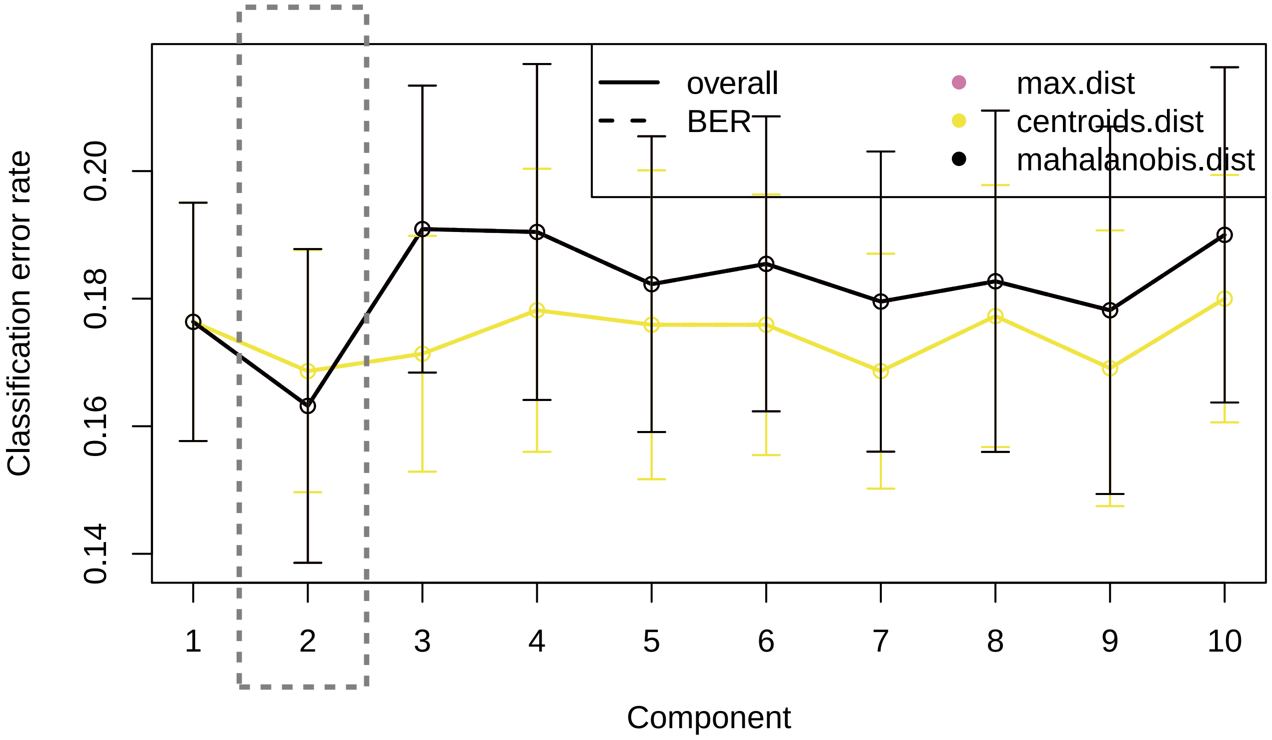

**Figure S4.** Principal component analysis score plot between the plasma metabolomic features in persons with AUD at the beginning (T1) and end (T2) of alcohol withdrawal.

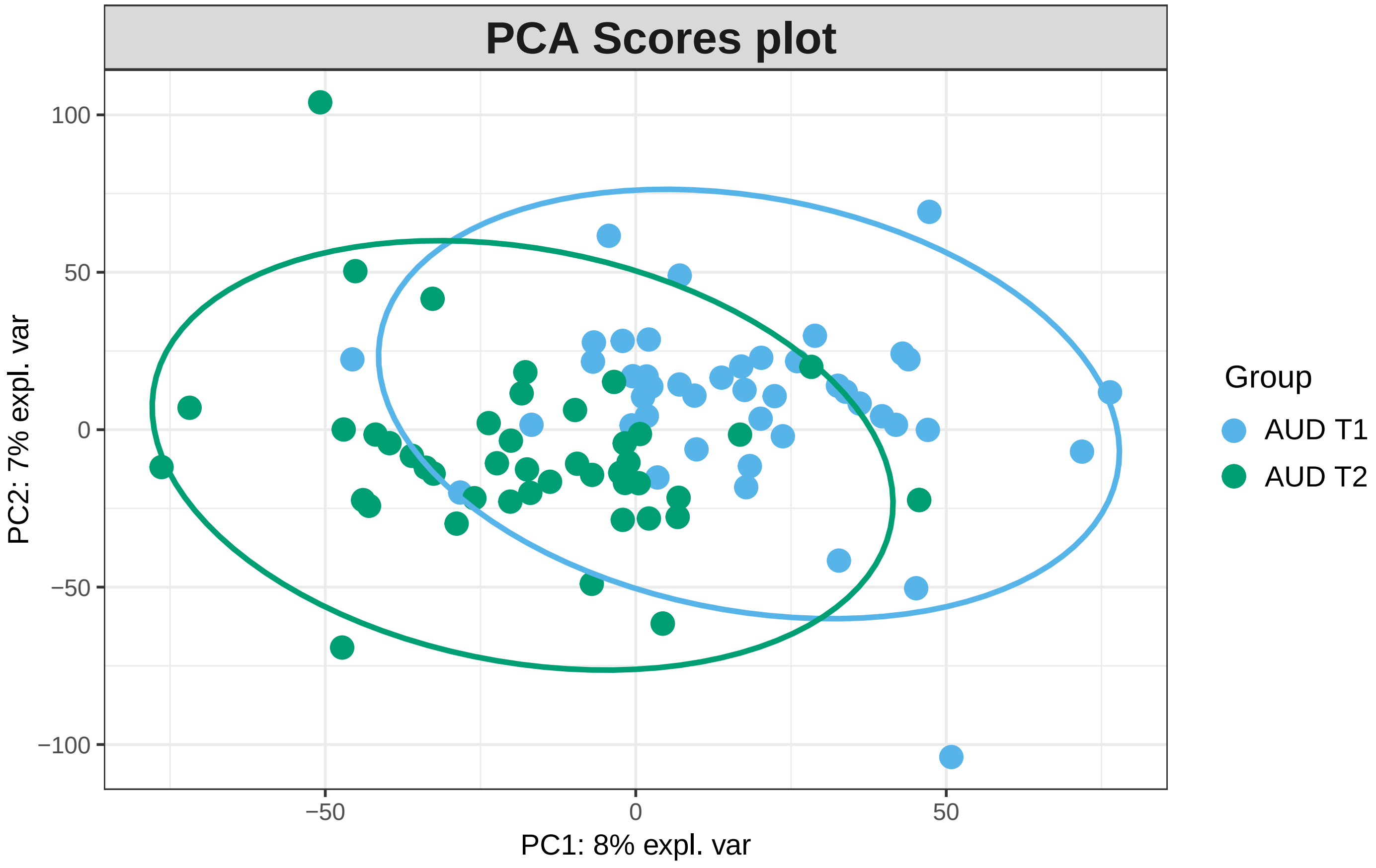

**Fig S5:**

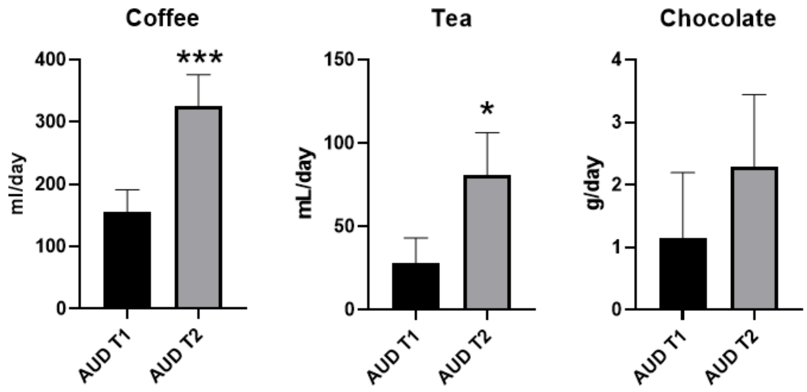


*FigS5: Changes in dietary intake of coffee, tea and chocolate during alcohol withdrawal. * p< 0.05; *** p < 0.001*
