## supplementary tables for "Blood metabolomic profiling reveals new targets in the management of psychological symptoms associated with severe alcohol use disorder"

### Supplemental tables

**Table S1.** biological features of healthy controls

| | ALCOHOLBIS cohort | GUT2BRAIN cohort | All | p value |
|---|---|---|---|---|
| Number of subjects | 19 | 13 | 32 | |
| Age | 43 ± 8 | 47 ± 12 | 45 ± 10 | 0.35 |
| Gender | | | | |
| Men, n (%) | 8 (42%) | 8 (61.5%) | 16 (50%) | 0.28$ |
| Women, n (%) | 11 (58%) | 5 (8.5%) | 16 (50%) | |
| Smoking status | | | | |
| Active smoker (%) | 1 (5%) | 3 (23%) | 4 (12.5%) | 0.14$ |
| Non-smoker (%) | 18 (95%) | 10 (77%) | 28 (87.5%) | |
| BMI | 23.6 ± 4.7 | 23.9 ± 3.2 | 23.7 ± 4.1 | 0.20 |
| AUDIT | | 3.3 ± 2.2 | | |

Results are means ± standard deviations. Independent t-tests to compare ALCOHOLBIS *versus* GUT2BRAIN cohorts; $chi-square test for categorical variables

**Table S2.** Background characteristics of the selected subjects from the TSDS cohort

| | Control (n = 100) | | Alcohol (n = 97) | | p value |
|---|---|---|---|---|---|
| PMI (days, mean ±SD) | 5.8 | ± 3.0 | 5.5 | ± 2.1 | 0.3081[a] |
| Age (years, mean ±SD) | 57 | ± 13 | 57 | ± 10 | 0.8320[a] |
| Sex (n females, %) | 12 | (12 %) | 22 | (22%) | 0.0738[b] |
| BMI (mean ±SD) | 30.9 | ± 8.0 | 28.0 | ± 7.2 | 0.0090[a] |
| Brain (grams, mean ±SD) | 1471 | ± 152 | 1448 | ± 148 | 0.2965[a] |
| Smoking (n, %)[c] | 32 | (55 %) | 37 | (67 %) | 0.1873[b] |
| CSF Alcohol (‰, mean ±SD)[d] | 0.13 | ± 0.37 | 0.94 | ± 0.67 | <0.0001[a] |
| Urine Alcohol (‰, mean ±SD)[e] | 0.28 | ± 0.54 | 1.12 | ± 0.56 | <0.0001[a] |

[a]Welch's t-test; [b] $\chi 2$ test; [c]smoking status known only from 113 subjects (58 controls, 55 heavy alcohol users); [d]CSF alcohol concentration only from 106 subjects (50 controls, 56 heavy alcohol users); [e]Urine alcohol concentration only from 38 subjects (18 controls, 20 heavy alcohol users); BMI, body mass index; CSF, cerebrospinal fluid; PMI, post-mortem interval
