## supplementary material for "Blood metabolomic profiling reveals new targets in the management of psychological symptoms associated with severe alcohol use disorder"

### 1. Assessement of psychological symptoms

The Beck Depression Inventory (BDI) is a 21-item self-report inventory designed to measure the severity of depressive symptoms, with a maximum score of 63. The validated French translation of the second version of the BDI (BDI-II) was used in this study (1).

The state report of the State-Trait Anxiety Inventory (STAI Form YA) is a valid and reliable 20-item self-report inventory for measuring the state of anxiety. The scores range from 20 to 80 where higher scores indicate greater anxiety. A valid French version was administered (2).

The Obsessive–Compulsive Drinking Scale (OCDS) is a questionnaire that assesses the cognitive aspects of alcohol craving during the preceding 7 days. This 14-question questionnaire provides a global craving score, as well as two subscores: an obsessive score (6 items) and a compulsive score (8 items). A valid French version was used in this study (3).

The amount of alcohol consumed the week before hospitalization was measured in grams per day using the time-line follow back approach (4).

### 2. LC-MS methodology

*LC-MS analysis*. Plasma sample preparation and LC-MS measurement followed the parameters previously detailed in Klåvus et al (Klåvus et al., 2020). Samples were randomized and thawed on ice before processing. 100 µl of plasma was added to 400 µl of LC-MS grade acetonitrile, mixed by pipetting four time, followed by centrifugation in 700 g for 5 minutes at 4 °C. A quality control sample was prepared by pooling 10 µl of each sample together. Extraction blanks having only cold acetonitrile and devoid of sample were prepared following the same procedure as sample extracts. LC-MS grade acetonitrile, methanol, water, formic acid and ammonium formate (Riedel-de Haën™, Honeywell, Seelze, Germany) were used to prepare mobile phase eluents in reverse phase (Zorbax Eclipse XDBC18, 2.1 × 100 mm, 1.8 μm, Agilent Technologies, Palo Alto, CA, USA) and hydrophilic interaction (Acquity UPLC® BEH Amide 1.7 μm, 2.1 × 100 mm, Waters Corporation, Milford, MA, USA) liquid chromatography separation. In reverse phase separation, the samples were analyzed by Vanquish Flex UHPLC system (Thermo Scientific, Bremen, Germany) coupled to high-resolution mass spectrometry (Q Exactive Focus, Thermo Scientific, Bremen, Germany) in both positive and negative polarity mass range from 120 to 1200, target AGC 1e6 and resolution 70,000 in full scan mode. Data dependent MS/MS data was acquired for both modes with target AGC 8e3 and resolution 17,500, precursor isolation window was 1.5 amu, normalized collision energies were set at 20, 30 and 40 eV and dynamic exclusion at 10.0 seconds. In hydrophobic interaction separation, the samples were analyzed by a 1290 LC system coupled to a 6540 UHD accurate mass Q-ToF spectrometer (Agilent Technologies, Waldbronn, Karlsruhe, Germany) using electrospray ionization (ESI, Jet Stream) in both positive and negative polarity with mass range from 50 to 1600 and scan rate of 1.67 Hz in full scan mode. Source settings were as in the protocol. Data dependent MS/MS data was acquired separately using 10, 20 and 40 eV collision energy in subsequent runs. Scan rate was set at 3.31 Hz, precursor isolation width of 1.3 amu and target counts/spectrum of 20,000, maximum of 4 precursor pre-cycle, precursor exclusion after 2 spectra and release after 15.0 seconds. Detectors were calibrated prior sequence and continuous mass axis calibration was performed throughout runs by monitoring reference ions from infusion solution for operating at high accuracy of < 2 ppm. Quality control samples were injected in the beginning of the analysis to equilibrate the system and after every 12 samples for quality assurance and drift correction in all modes. All data were acquired in centroid mode by either MassHunter Acquisition B.05.01 (Agilent Technologies) or in profile mode by Xcalibur 4.1 (Thermo Fisher Scientific) softwares.

Metabolomics analysis of TSDS frontal cortex and CSF samples using the same 1290 LC system coupled with a 6540 UHD accurate mass Q-ToF spectrometer has been previously accomplished by Kärkkäinen et al (Kärkkäinen et al., 2021).

*Peak picking and data processing*. Raw instrumental data (*raw and *.d files) were converted to ABF format using Reifycs Abf Converter (https://www.reifycs.com/AbfConverter). MS-DIAL (Version 4.70) was employed for automated peak picking and alignment with the parameters according to Klåvus et al., 2020 (Klåvus et al., 2020) separately for each analytical mode. For the 6540 Q-ToF mass data minimum peak height was set at 8,000 and for the Q Exactive Focus mass data minimum peak height was set at 850,000. Commonly, m/z values up to 1600 and all retention times were considered, for aligning the peaks across samples retention time tolerance was 0.2 min and MS1 tolerance 0.015 Da and the “gap filling by compulsion” was selected. Alignment results across all modes and sample types as peak areas were exported into Microsoft Excel sheets to be used for further data pre-processing.

Pre-processing including drift correction and quality assessment was done using the notame package v.0.2.1 R software version 4.0.3 separately for each mode. Features present in less than 80% of the samples within all groups and with detection rate in less than 70% of the QC samples were flagged. All features were subjected to drift correction where the features were log-transformed and a regularized cubic spline regression line was fitted for each feature against the quality control samples. After drift correction, QC samples were removed and missing values in the non-flagged features were imputed using random forest imputation. Finally, the preprocessed data from each analytical mode was merged into a single data matrix.

Molecular feature characteristics (exact mass, retention time and MS/MS spectra) were compared against in-house standard library, publicly available databases such as METLIN, HMDB and LIPIDMAPS and published literature. Annotation of metabolites and the level of identification was based on the recommendations given by the Chemical Analysis Working Group (CAWG) Metabolomics Standards Initiative (MSI) (Sumner et al., 2007): 1 = identified based on a reference standard, 2 = putatively annotated based on physicochemical properties or similarity with public spectral libraries, 3 = putatively annotated to a chemical class and 4 = unknown.”

### 3. EICs, retention times and reference spectra for level 1 identifications

1-Methylhistidine

RT std: 6.21min, RT experimental: 6.42min, RT Δ 0.21min

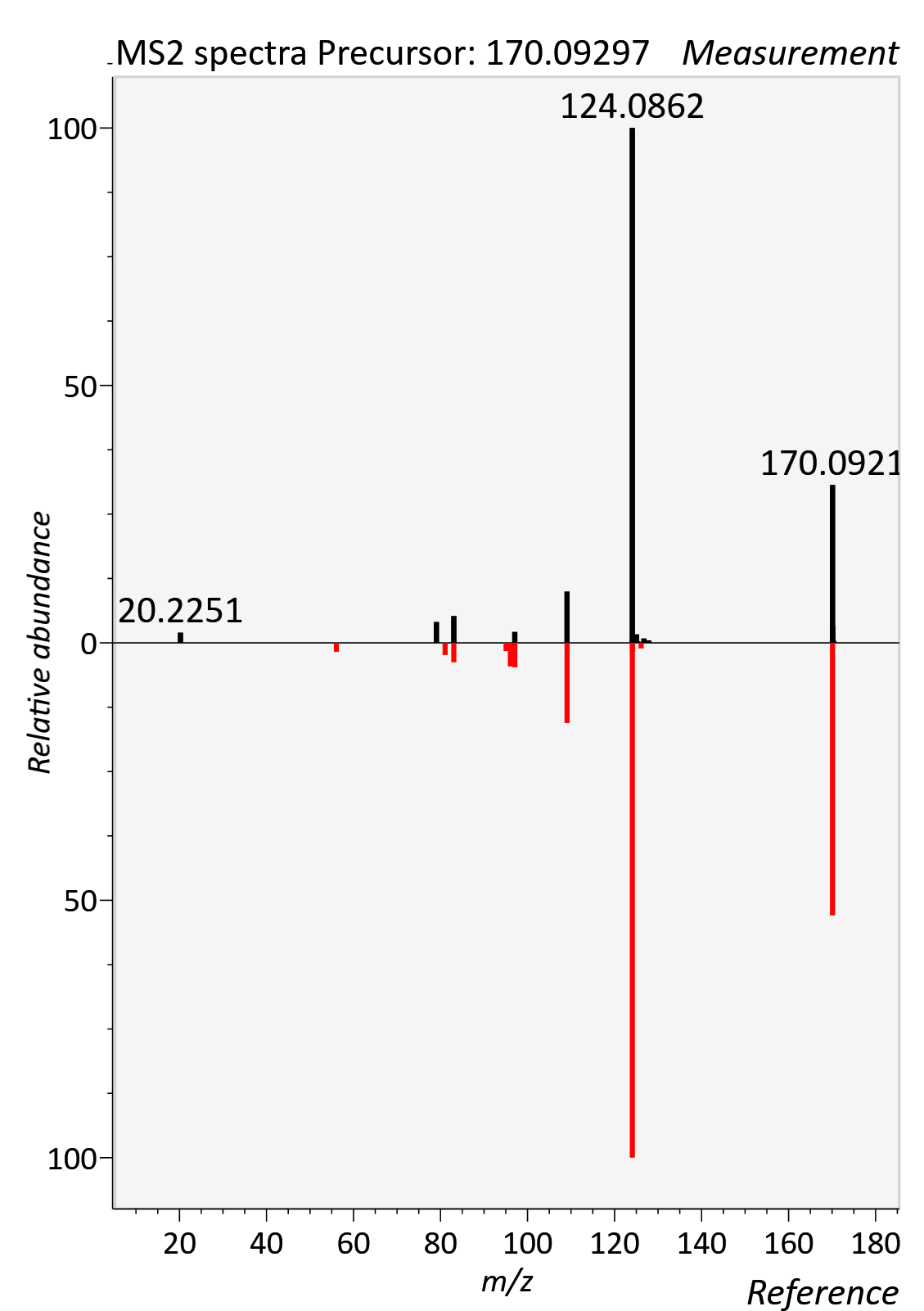

1-Methylnicotincamide

RT std: 2.18min, RT experimental: 2.39min, RT Δ 0.21min

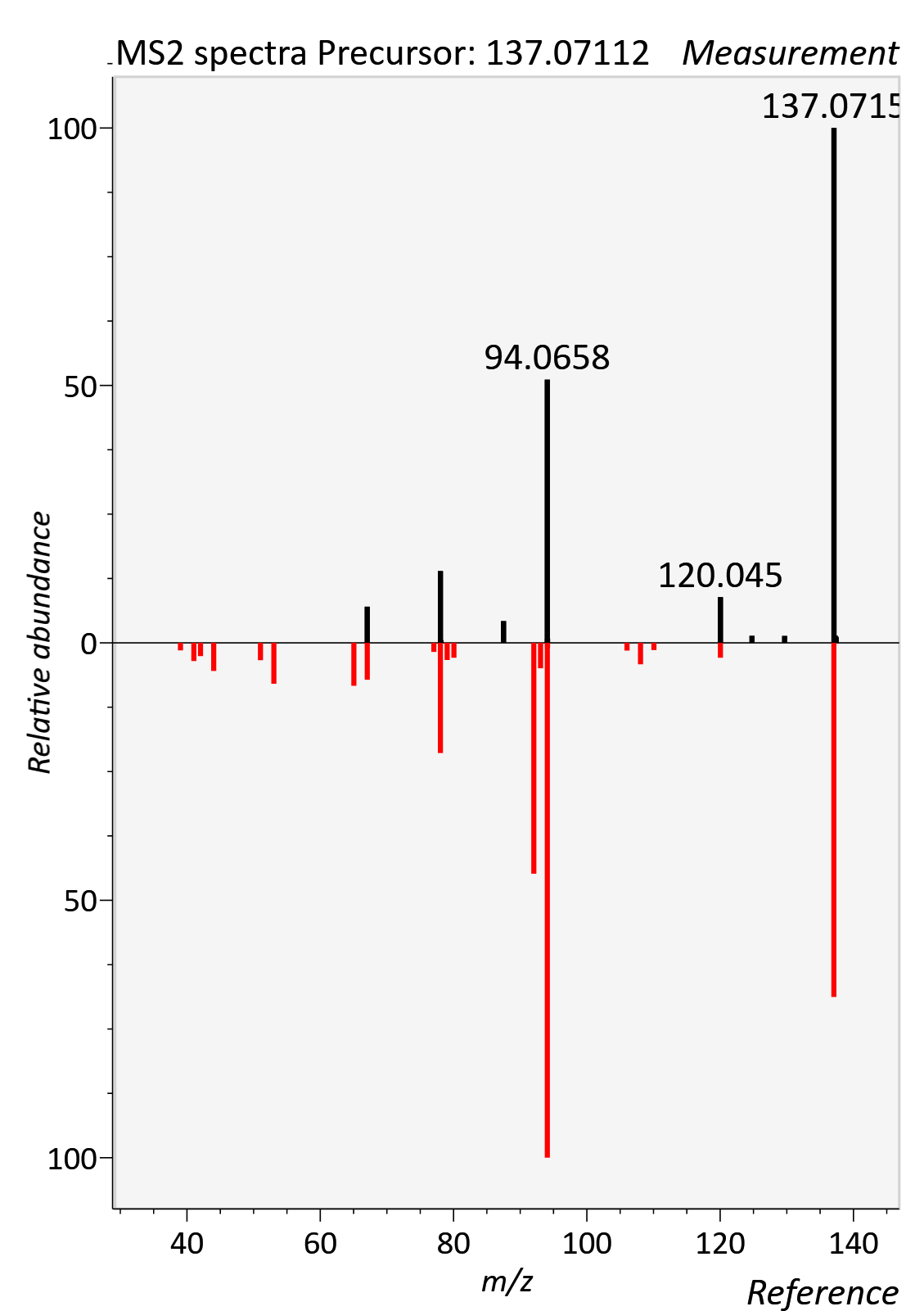

3-Indoleacetic acid

RT std: 5.00min, RT experimental: 5.28min, RT Δ 0.28min

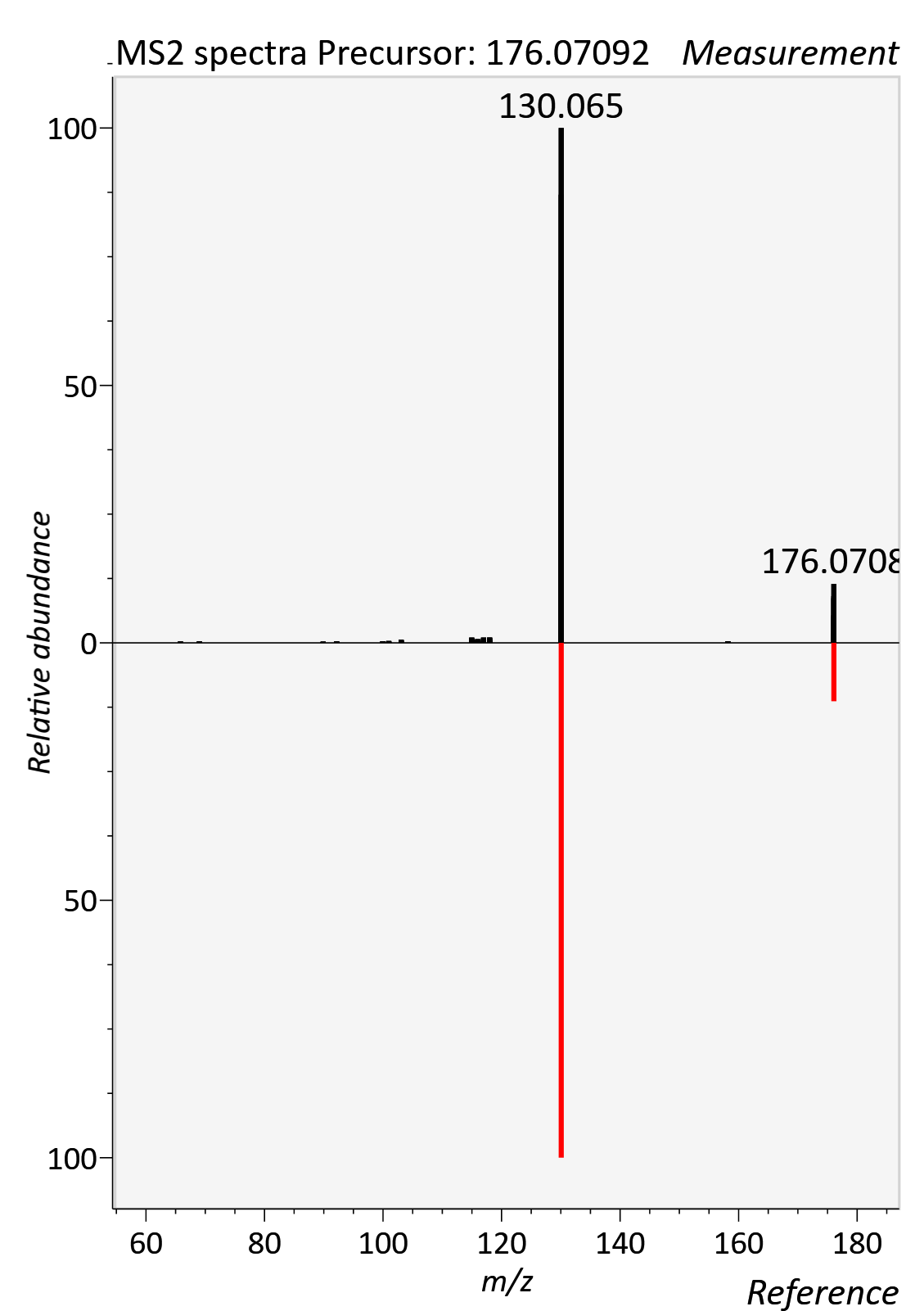

4-Trimethylammoniobutanoic acid

RT std: 3.36min, RT experimental: 3.60min, RT Δ 0.06min

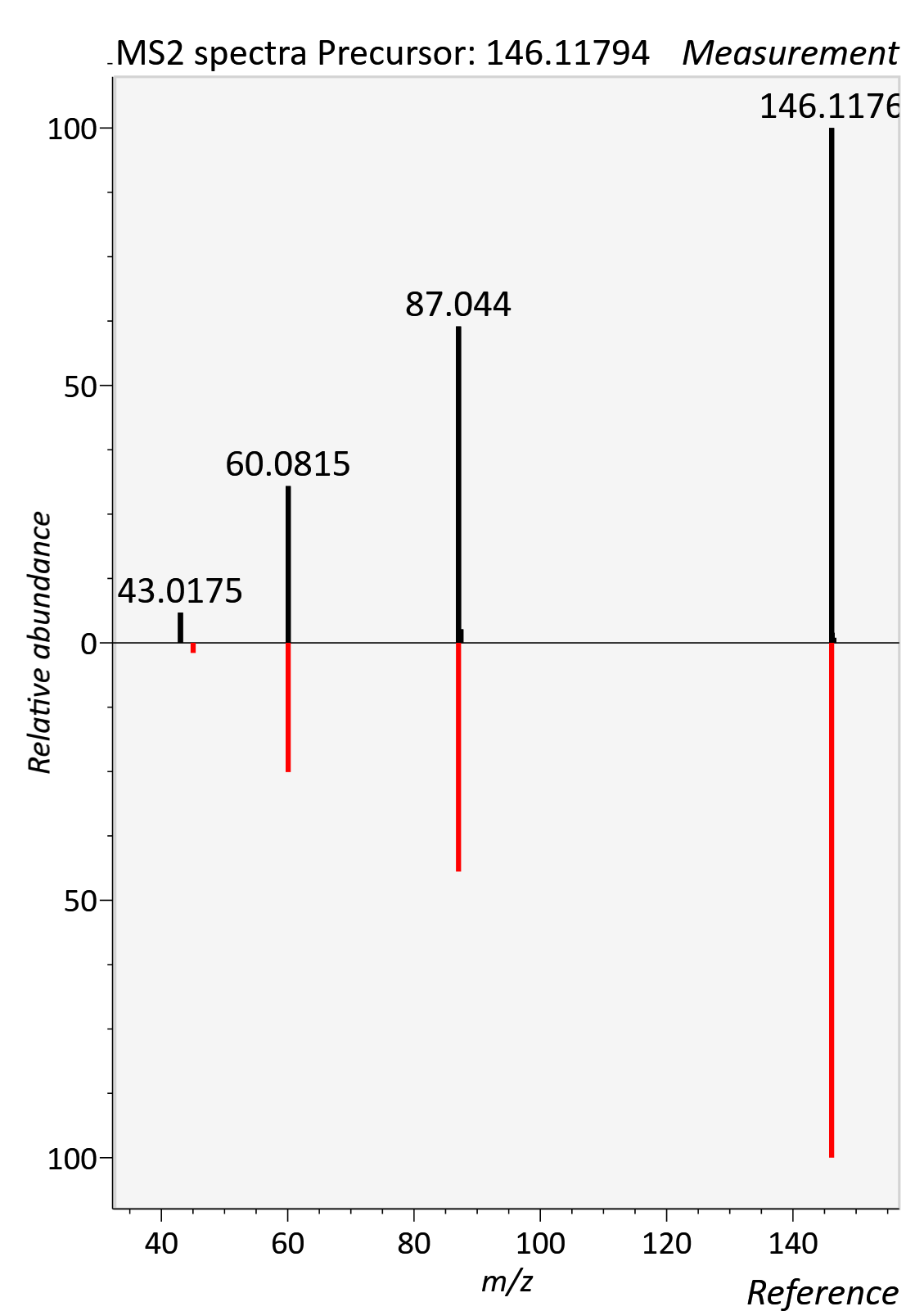

5-AVAB

RT std: 2.39min, RT experimental: 2.27min, RT Δ 0.12min

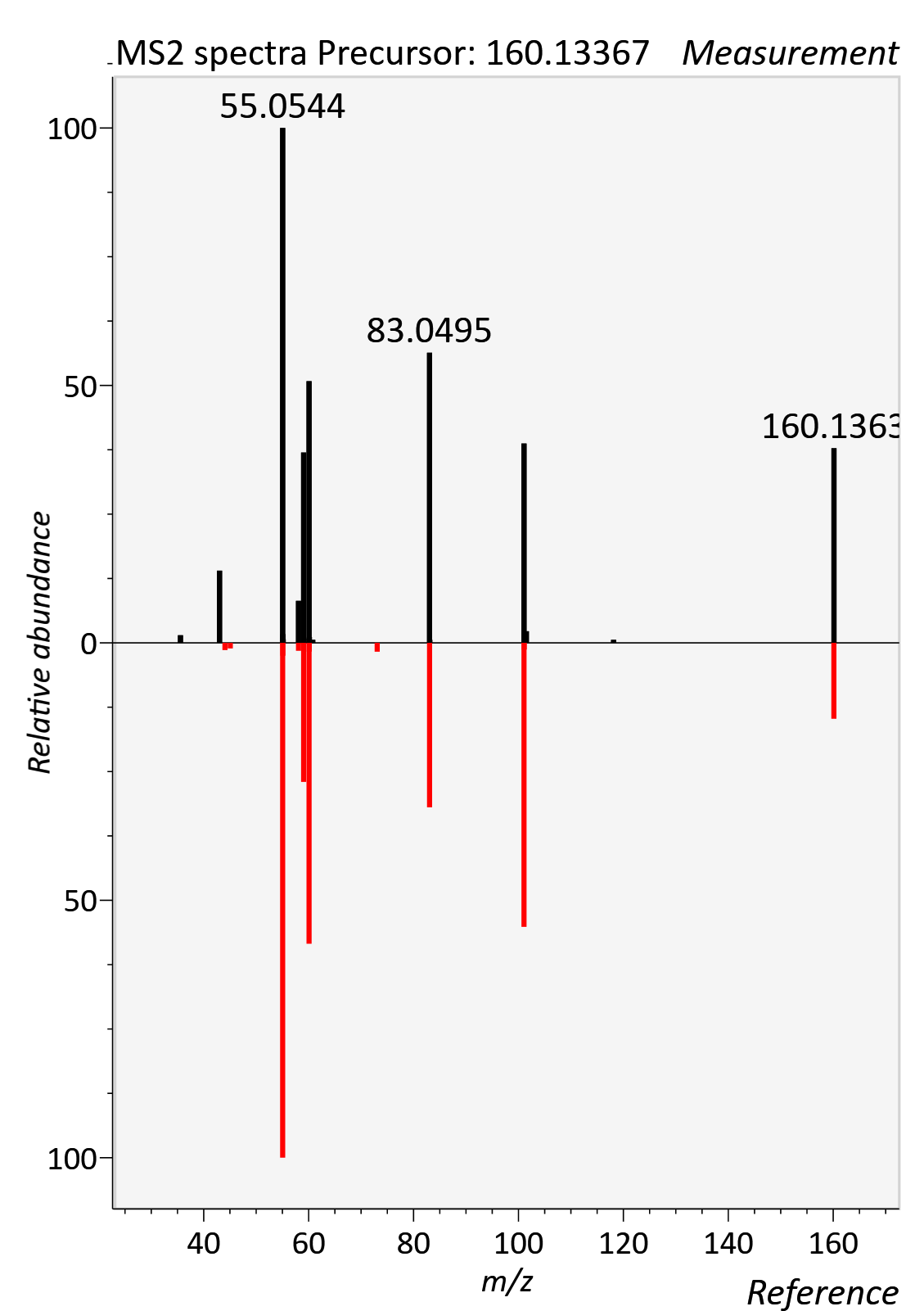

AC 05:0 (Isovalerylcarnitine)

RT std: 1.28min, RT experimental: 1.40min, RT Δ 0.12min

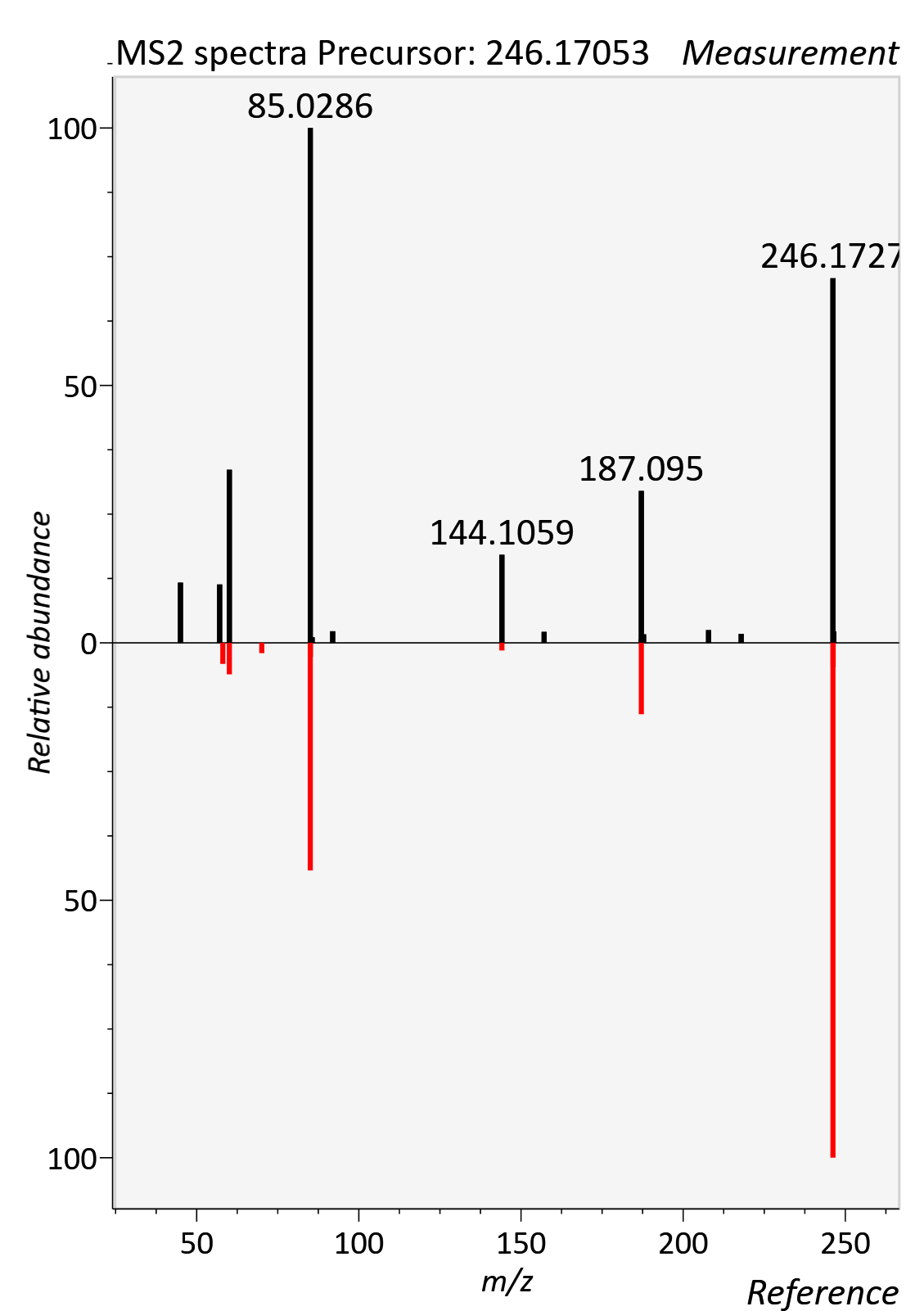

7a-hydroxy-3-oxo-4-cholestenoic acid

RT std: 9.71min, RT experimental: 9.94min, RT Δ 0.23min

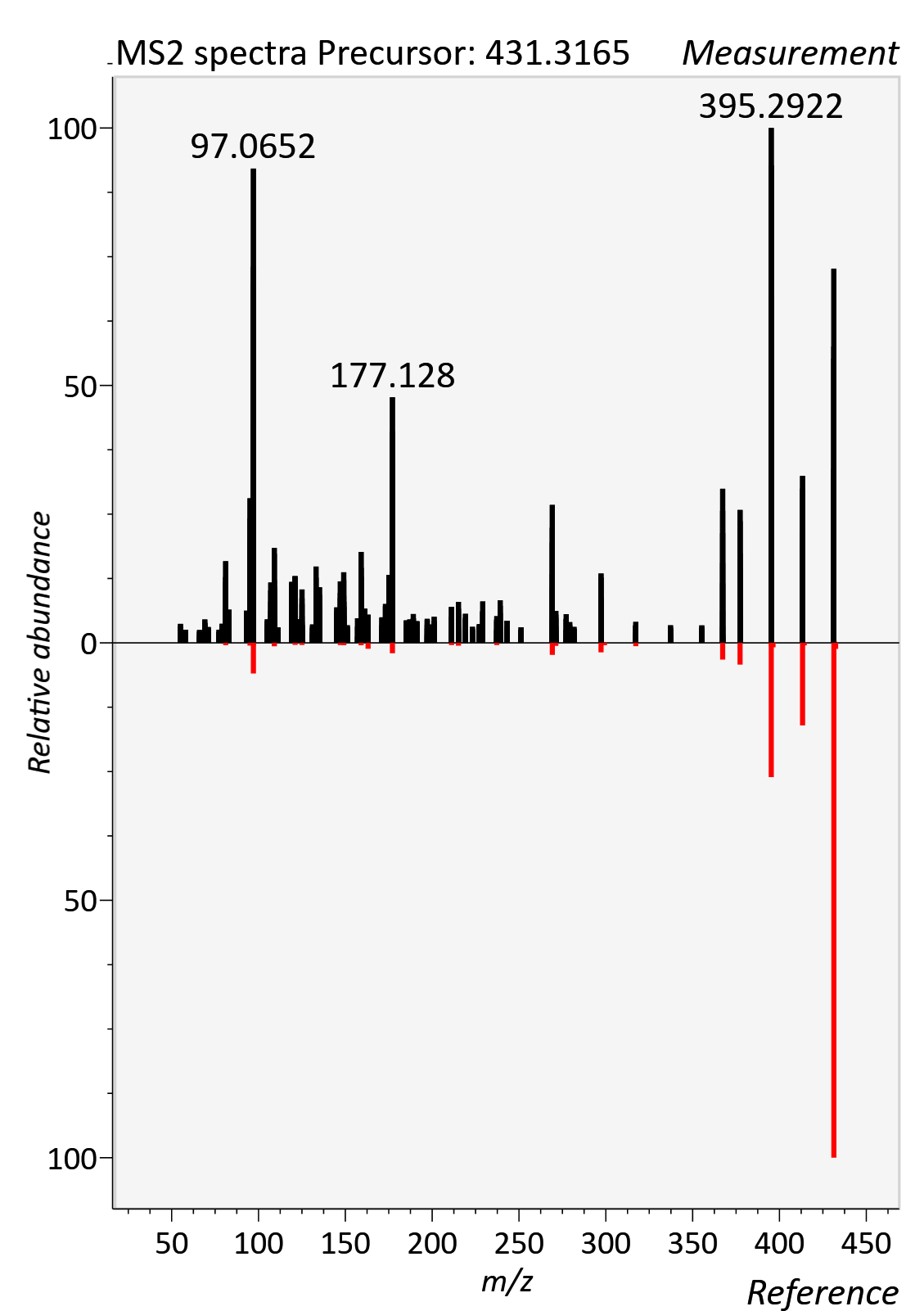

AC 04:0 (Butanoylcarnitine)

RT std: 1.85min, RT experimental: 1.77min, RT Δ 0.08min

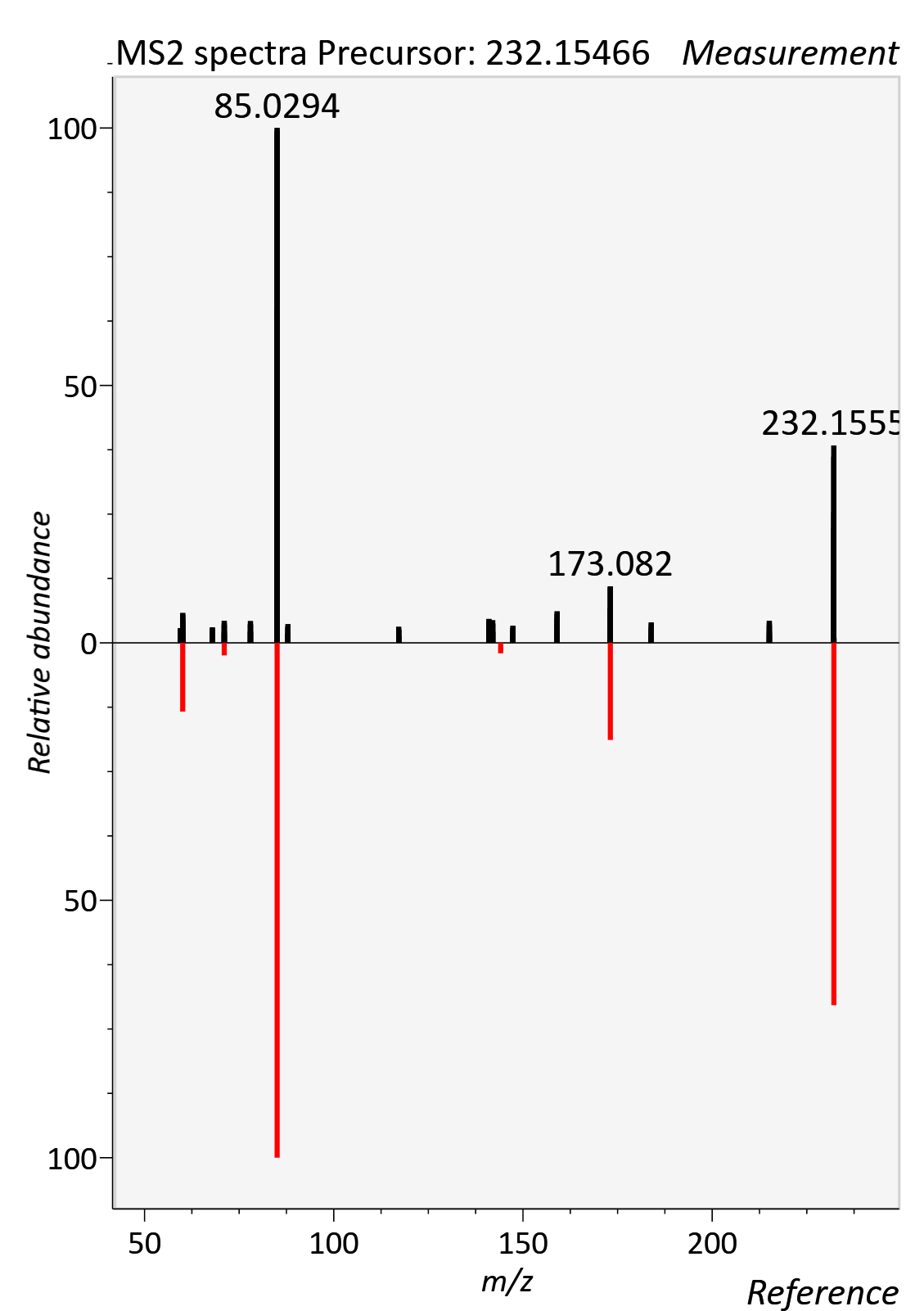

AC 06:0 (Hexanoylcarnitine)

RT std: 4.09min, RT experimental: 4.20min, RT Δ 0.11min

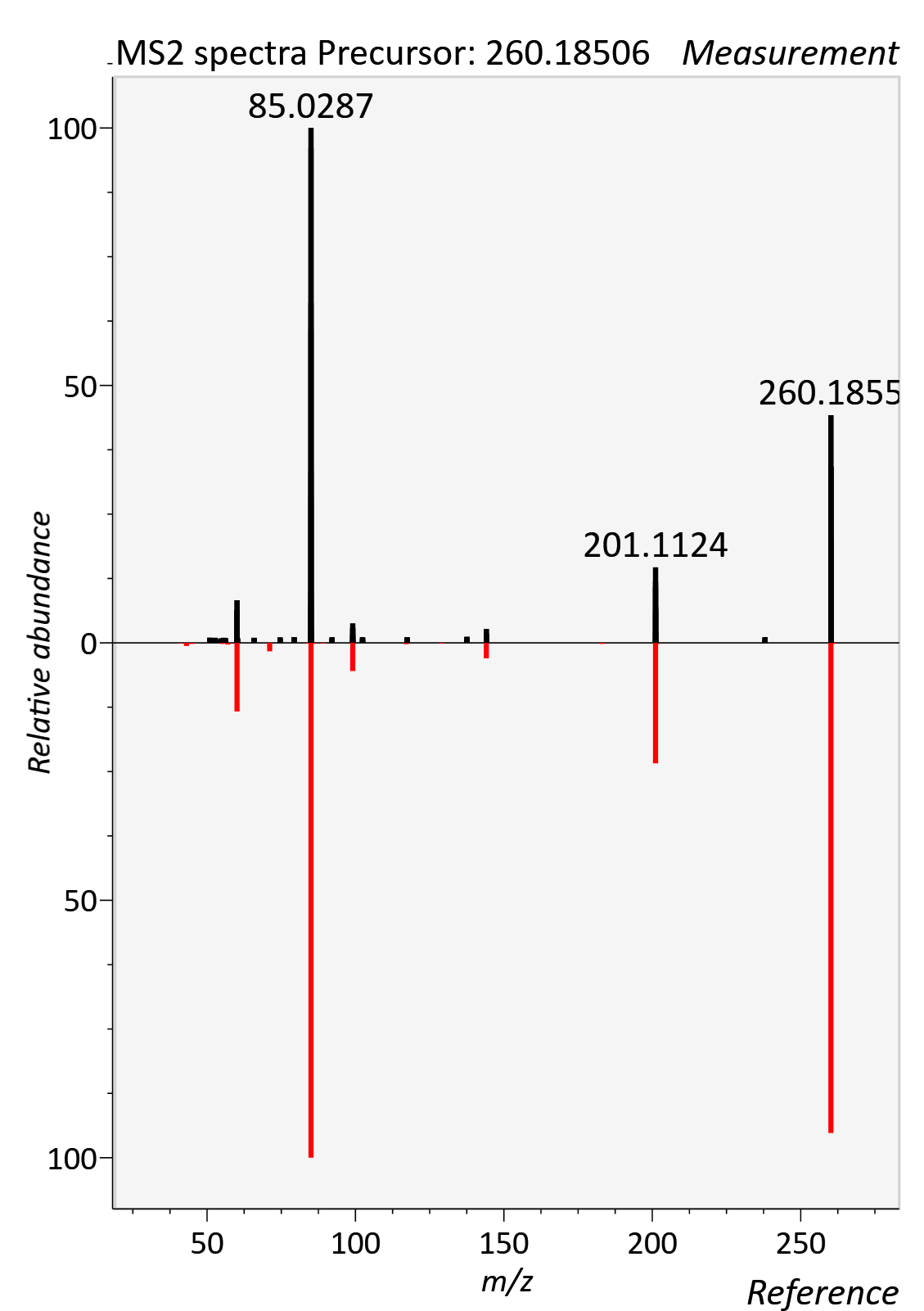

AC 08:0 (Octanoylcarnitine)

RT std: 5.89, RT experimental: 6.02min, RT Δ 0.13min

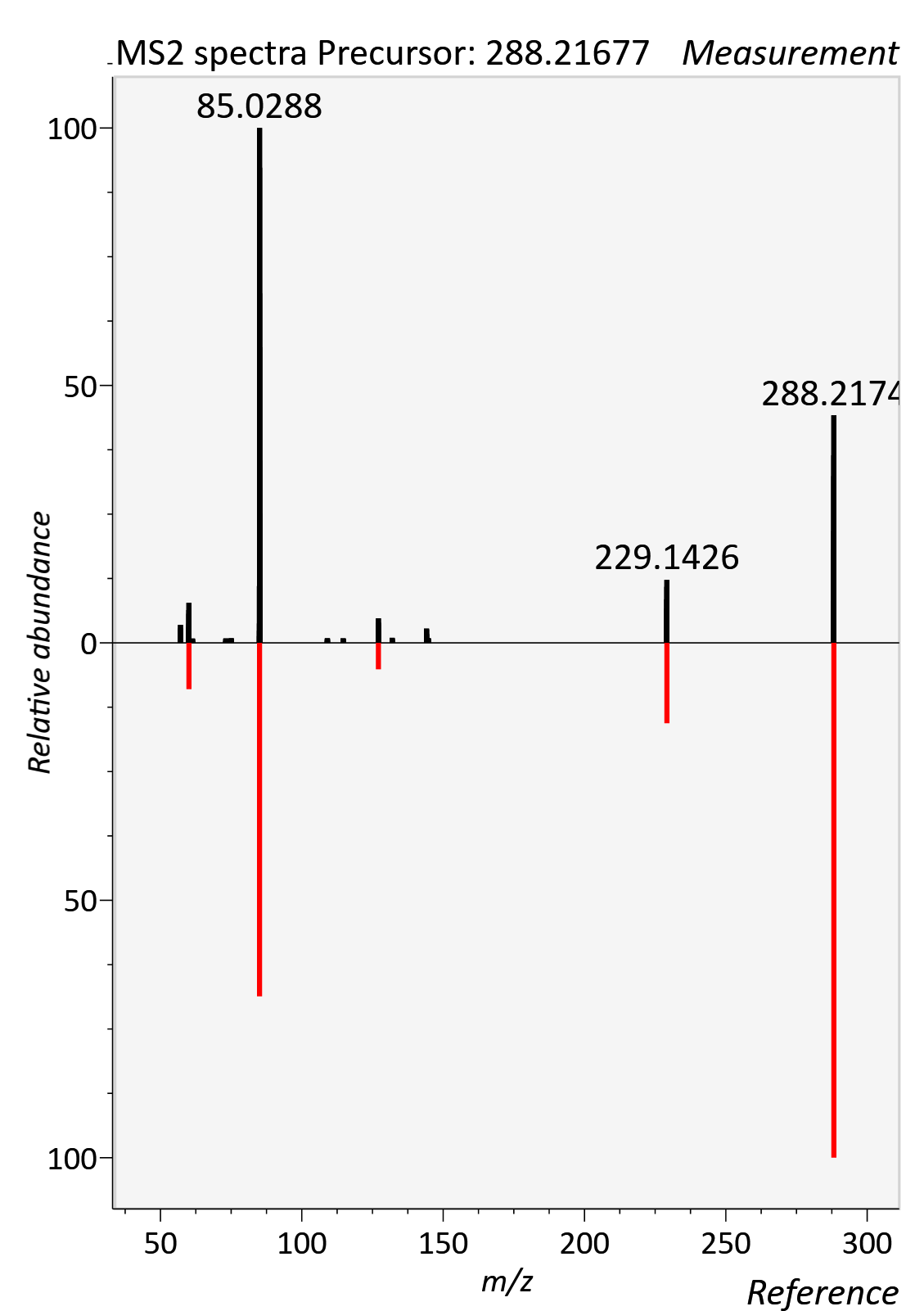

AC 08:1 (Octenoyl-L-carnitine)

RT std: 5.56min, RT experimental: 5.20min, RT Δ 0.23min

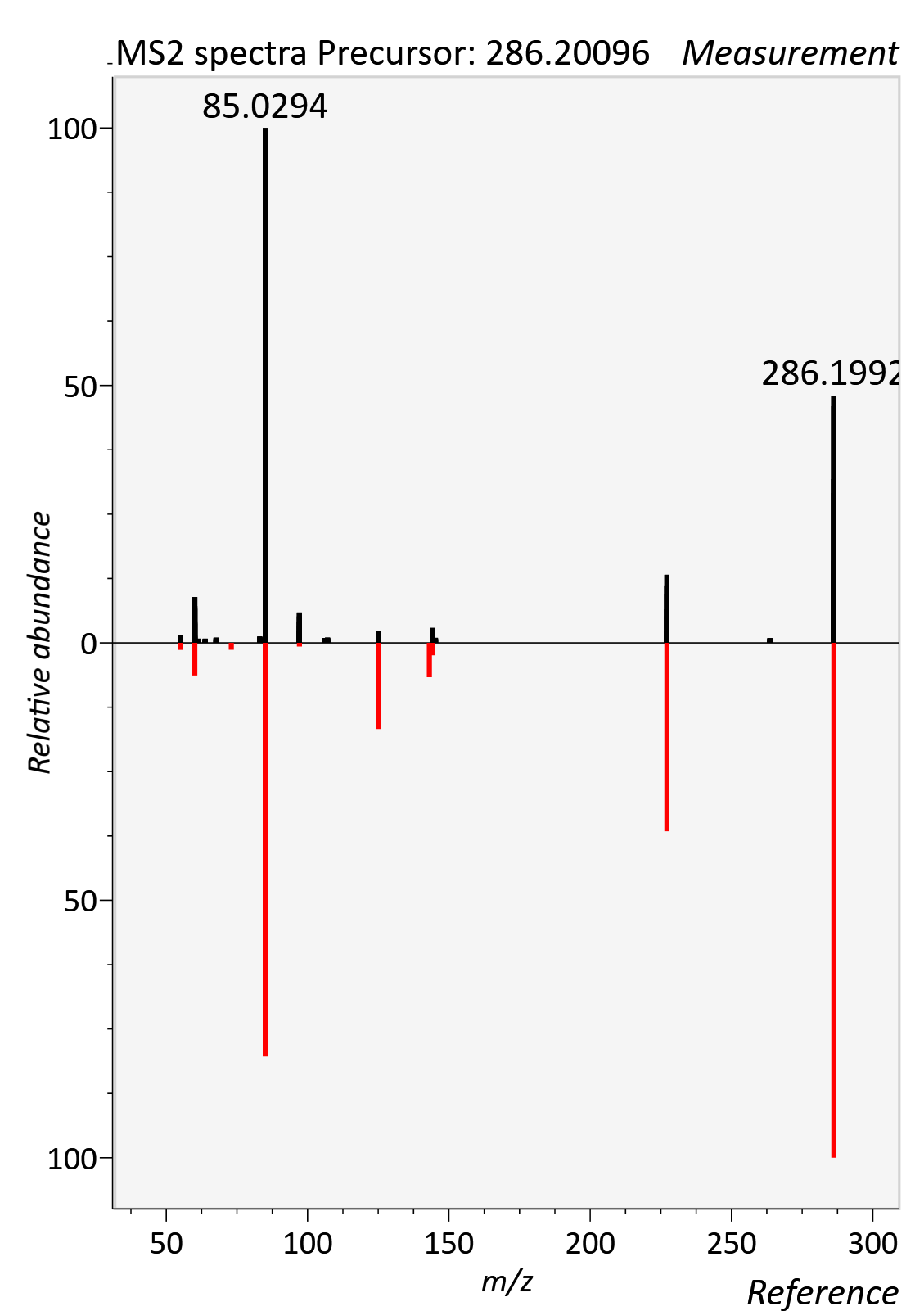

AC 10:0 (Decanoylcarnitine)

RT std: 7.17min, RT experimental: 7.28min, RT Δ 0.11min

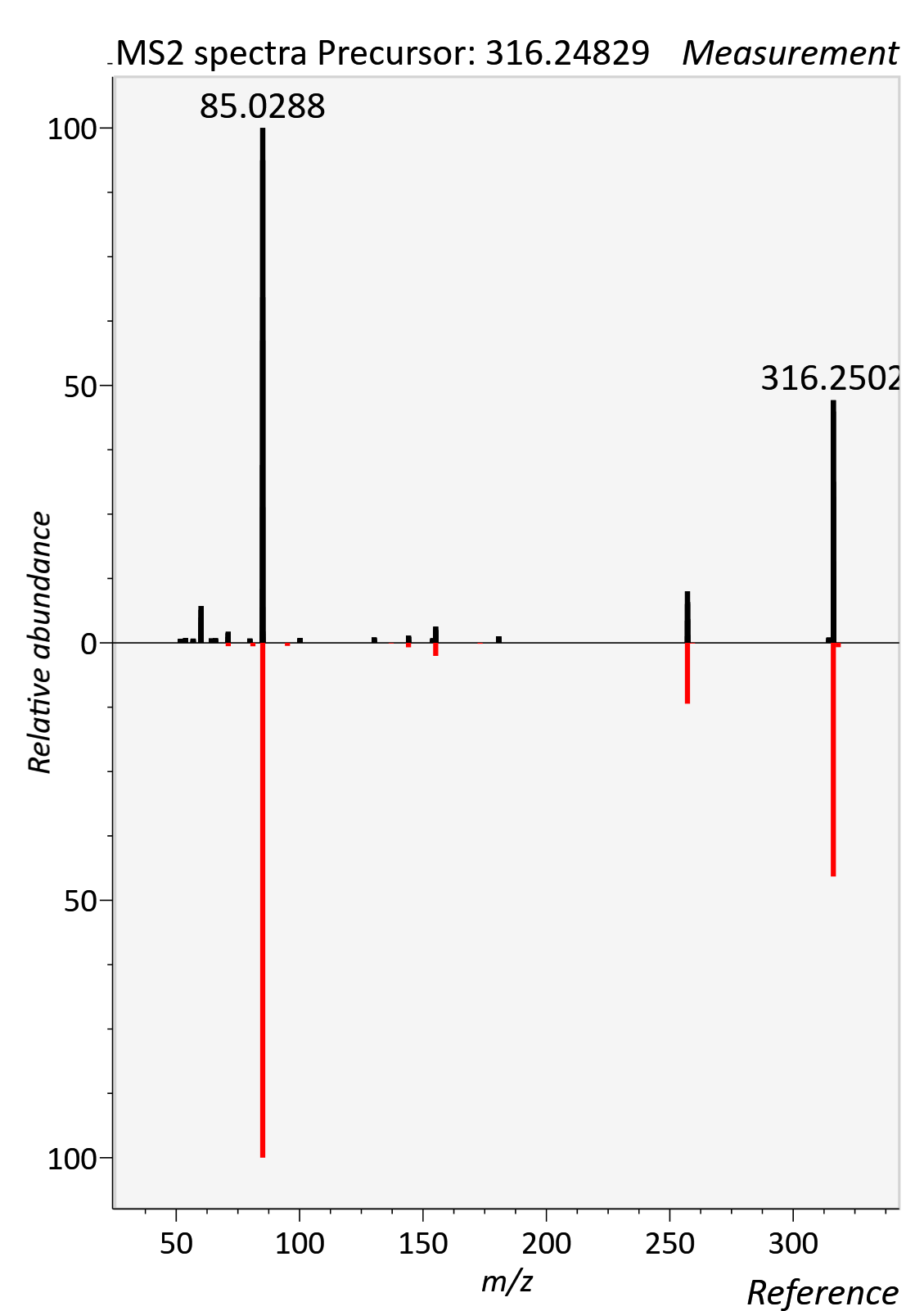

AC 12:0 (Dodecanoylcarnitine)

RT std: 8.09min, RT experimental: 8.17min, RT Δ 0.08min

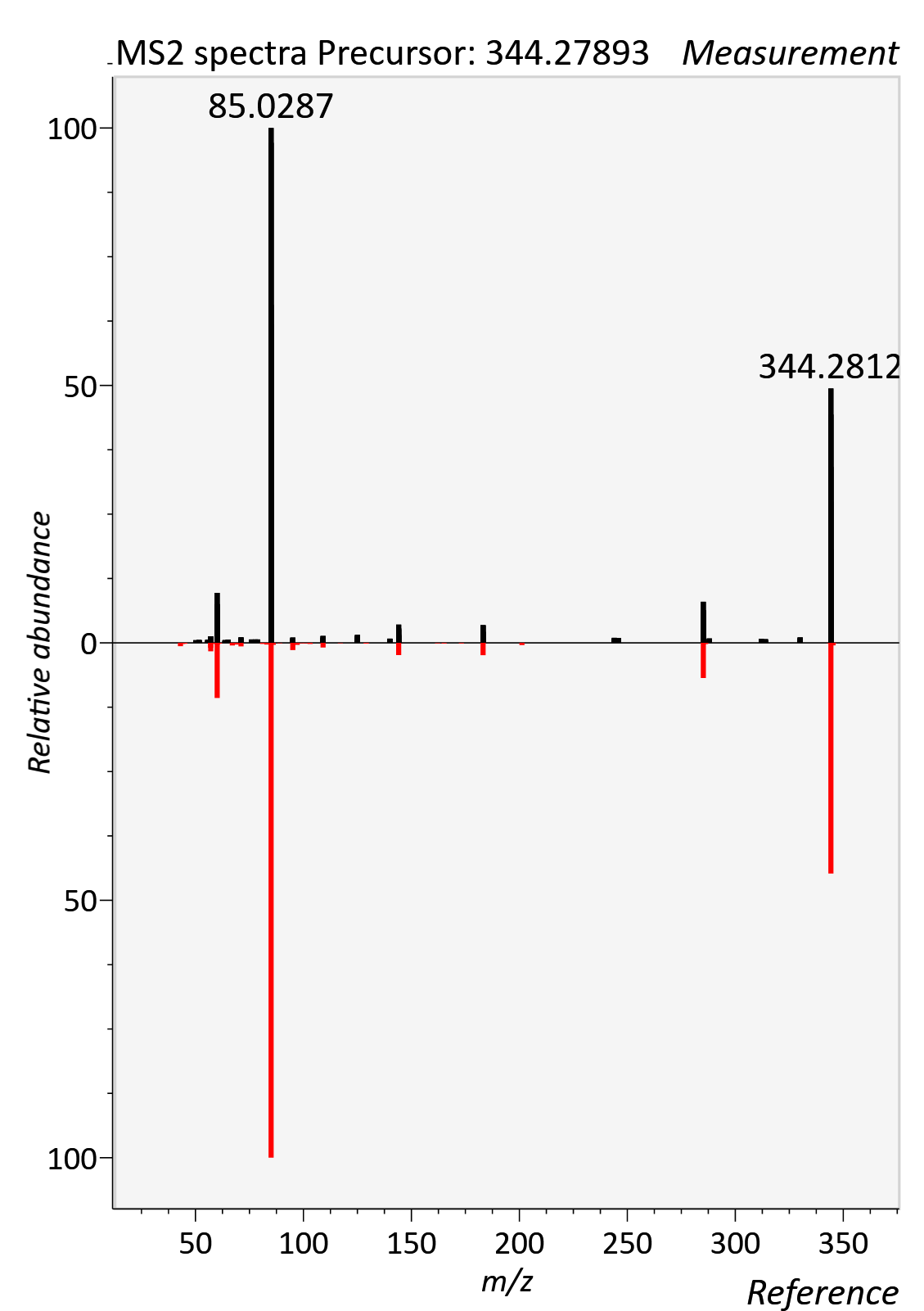

AC 12:1 (Dodecenoylcarnitine)

RT std: 7.91min, RT experimental: 7.76min, RT Δ 0.15min

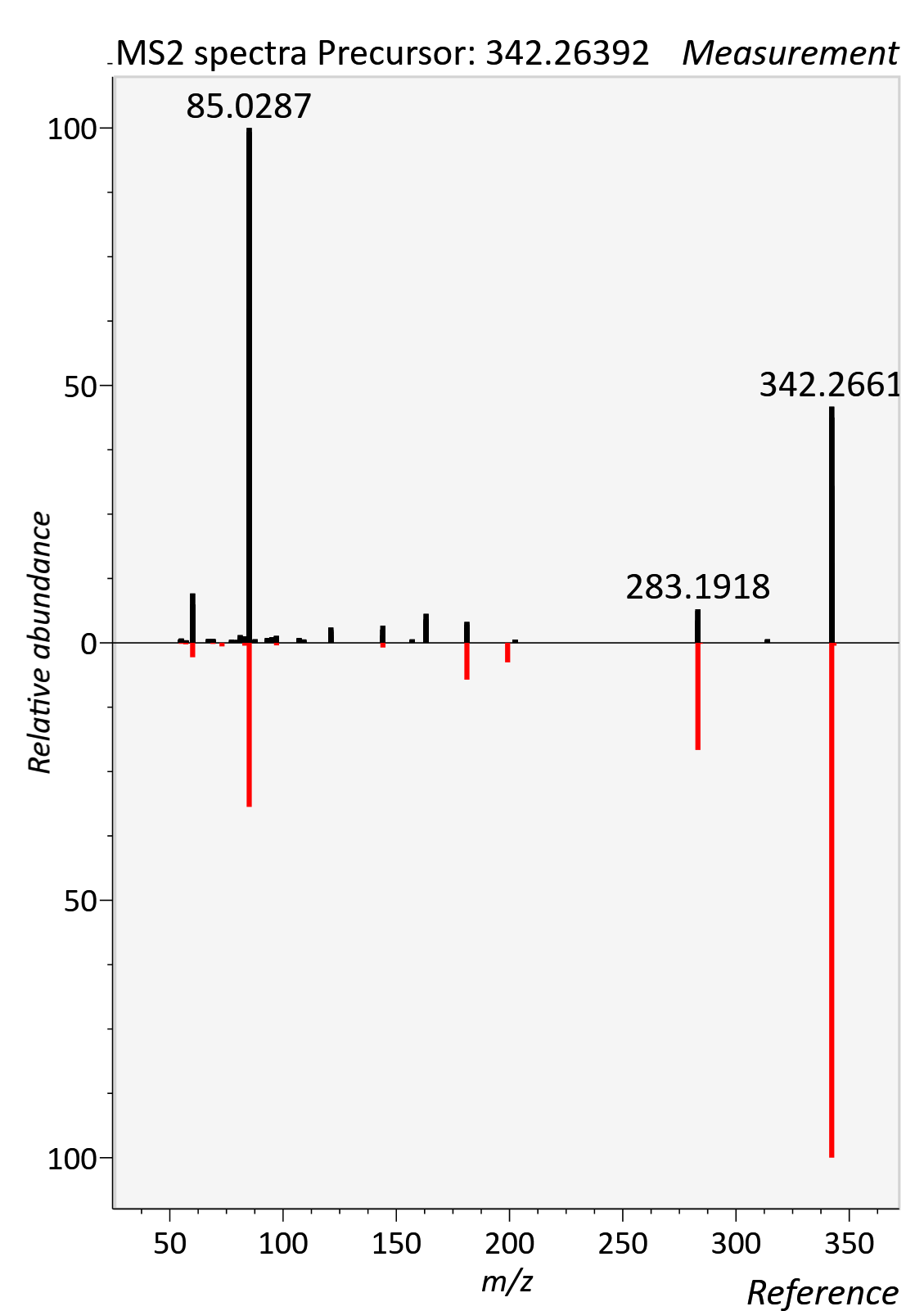

AC 14:0 (Tetradecanoyl-L-carnitine)

RT std: 8.60min, RT experimental: 8.80min, RT Δ 0.20min

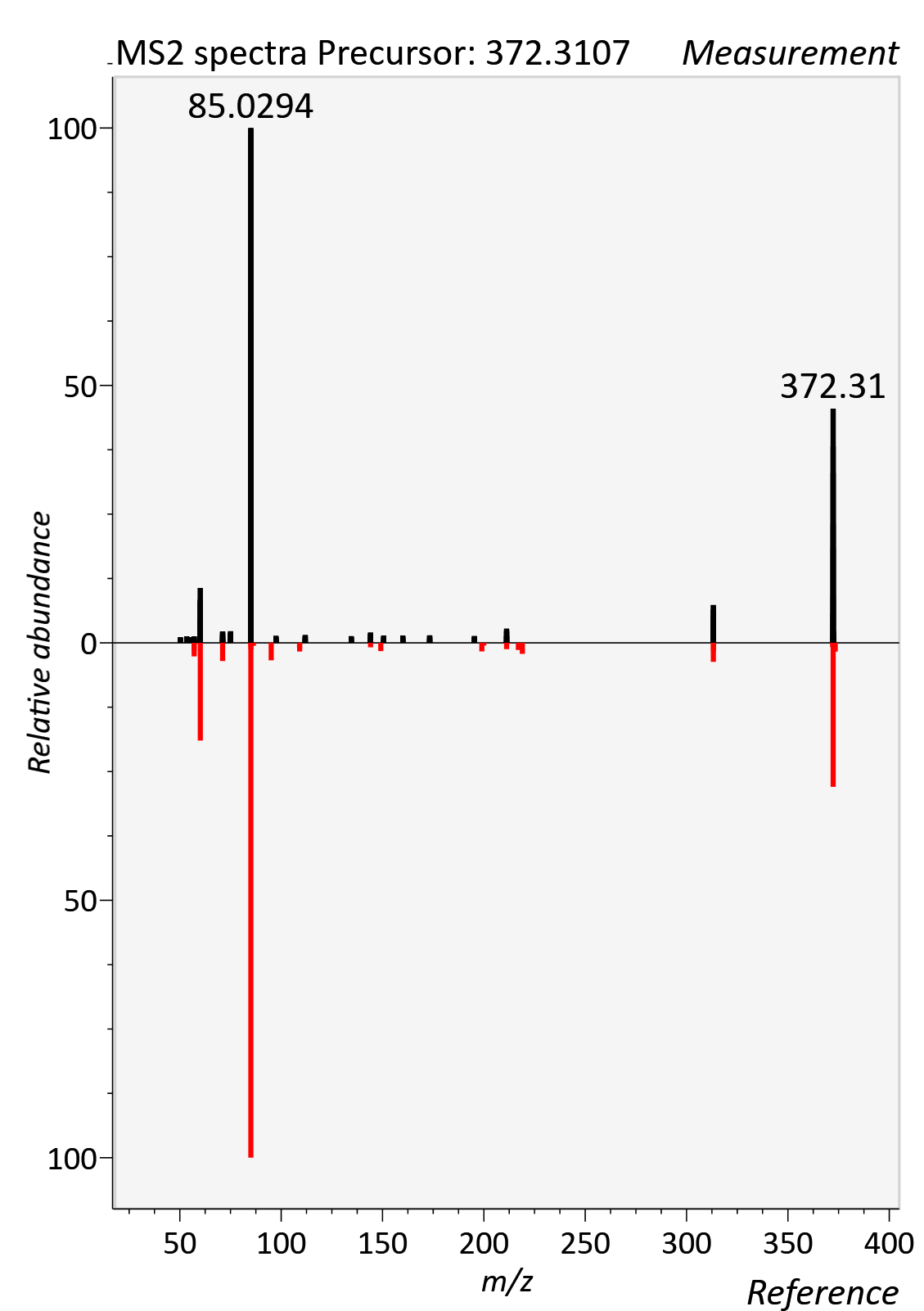

AC 14:1 (Tetradecenoylcarnitine)

RT std: 8.65min, RT experimental: 8.48min, RT Δ 0.23min

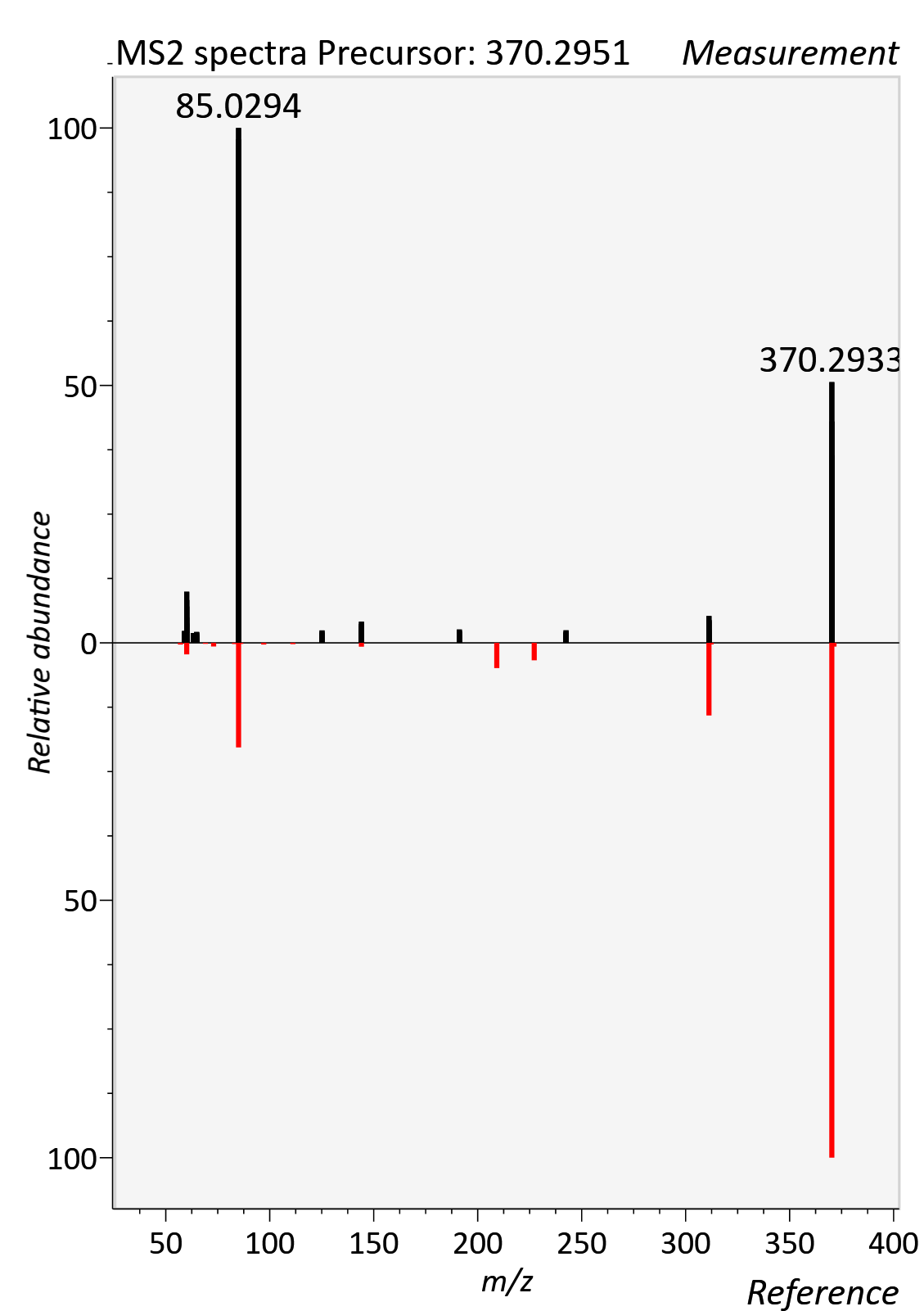

AC 16:0 (Palmitoylcarnitine)

RT std: 9.13min, RT experimental: 9.30min, RT Δ 0.20min

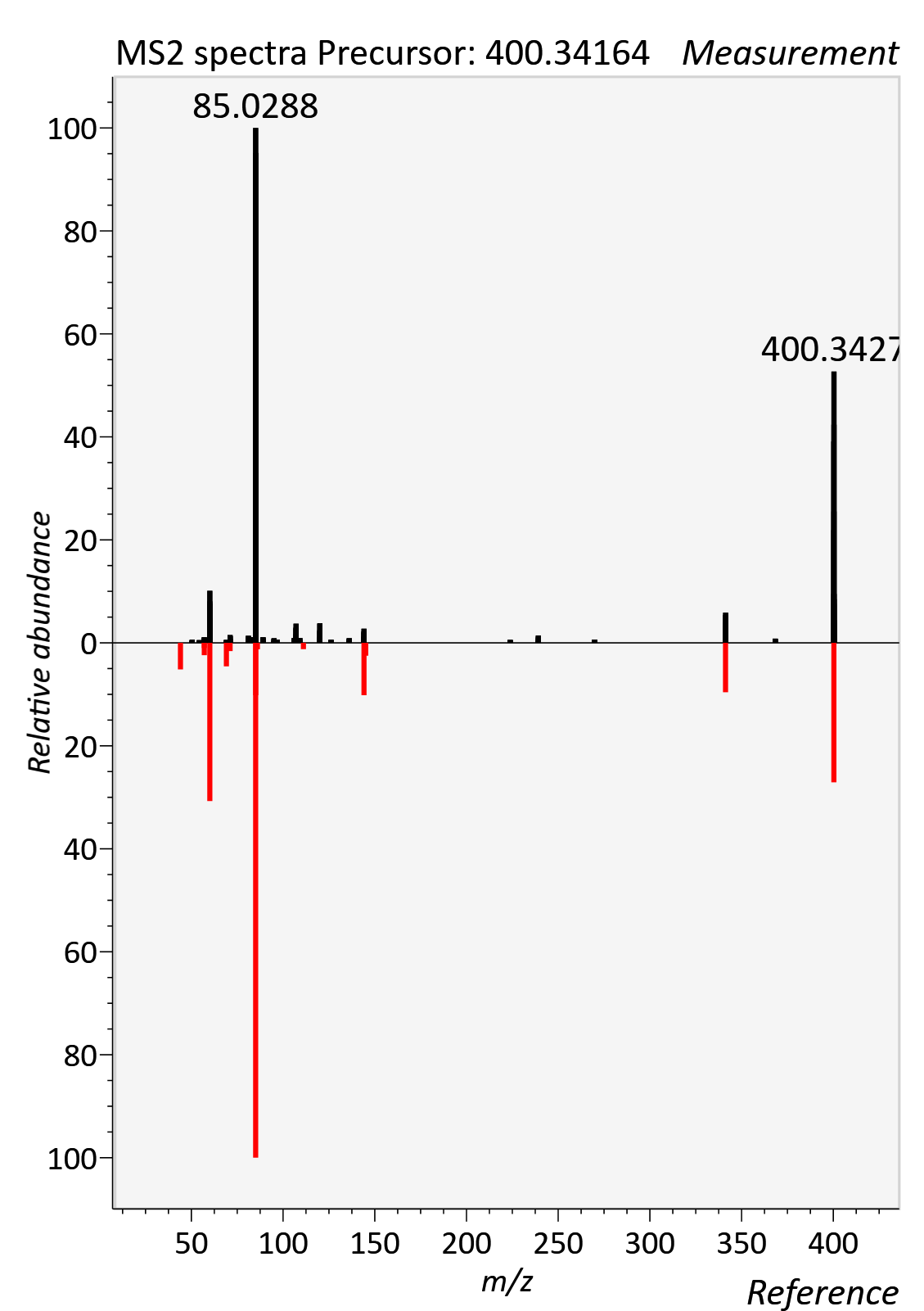

AC 18:1 (Oleylcarnitine)

RT std: 9.34min, RT experimental: 9.44min, RT Δ 0.10min

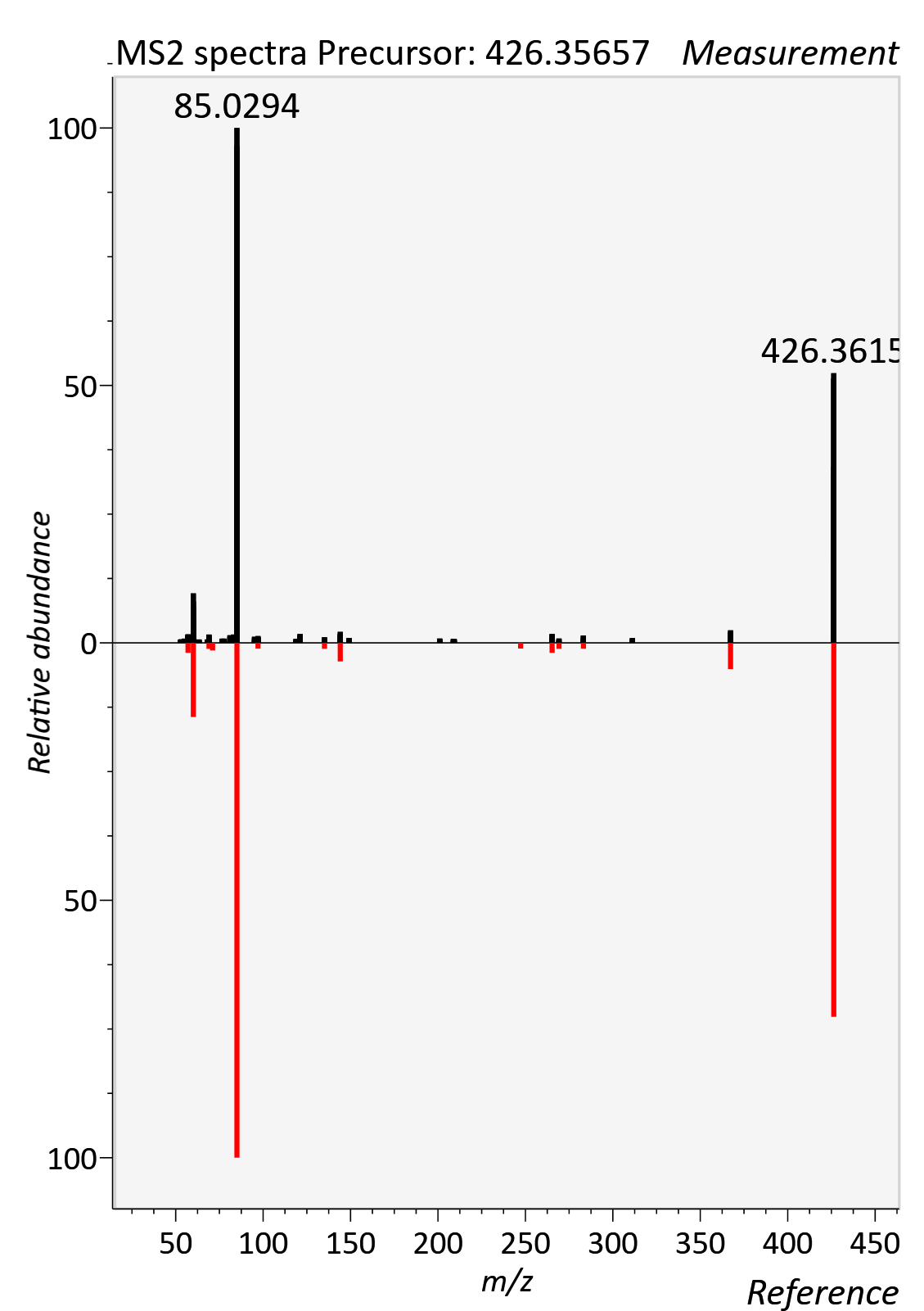

AC 18:2 (Linoleylcarnitine)

RT std: 9.15min, RT experimental: 9.17min, RT Δ 0.02min

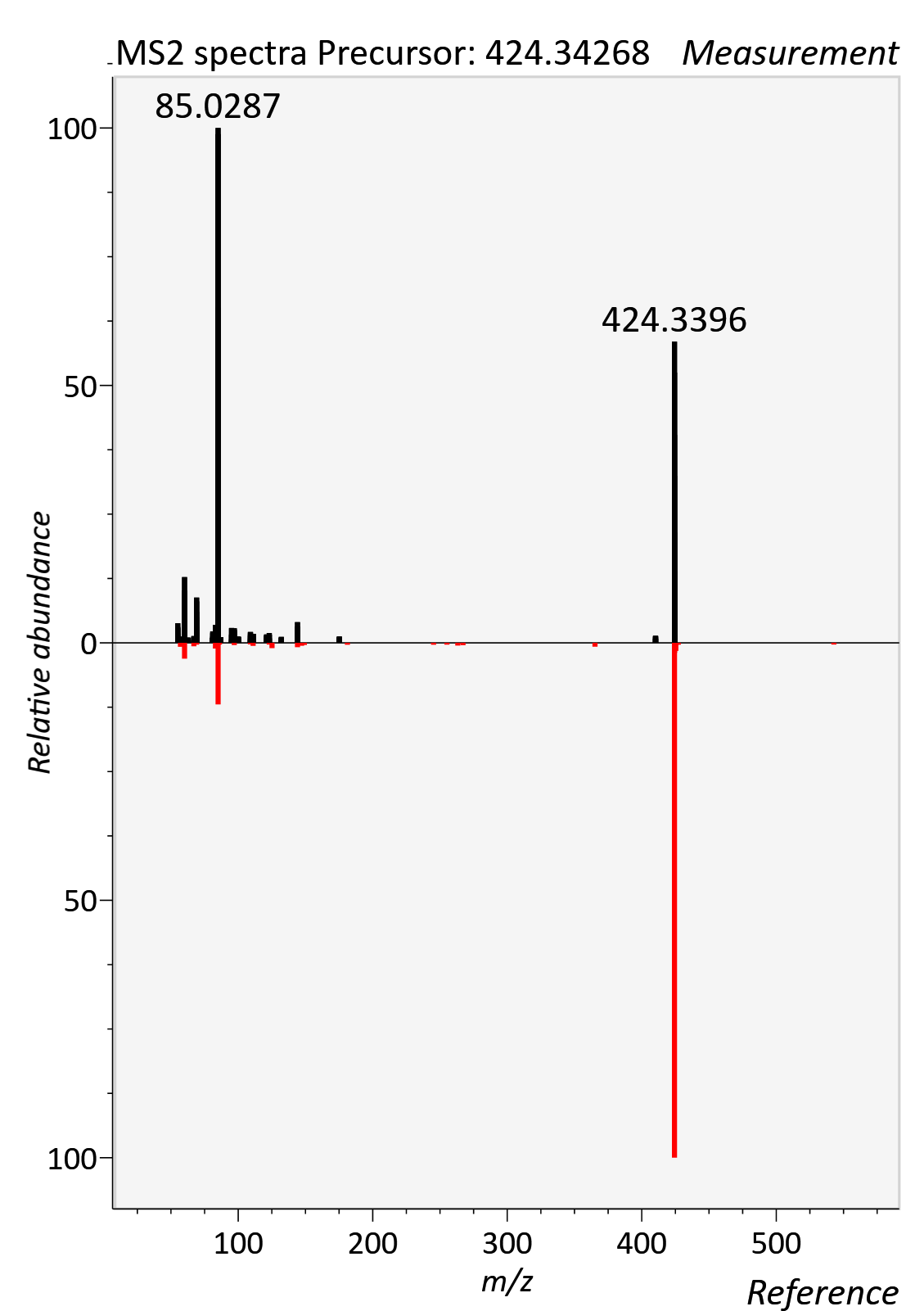

Acetylcarnitine

RT std: 2.71min, RT experimental: 3.25min, RT Δ 0.54min

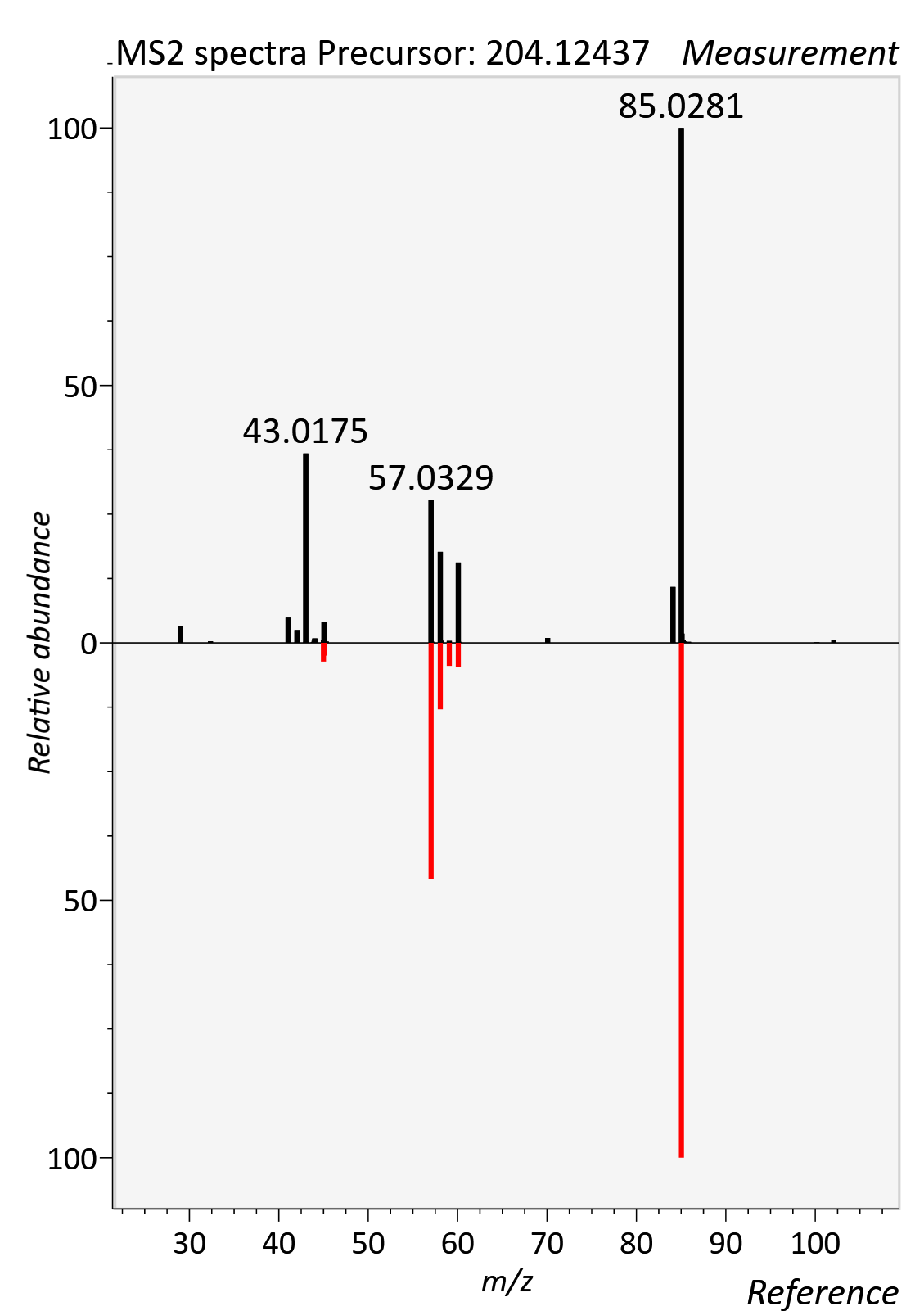

Caffeine

RT std: 3.67min, RT experimental: 3.80min, RT Δ 0.13min

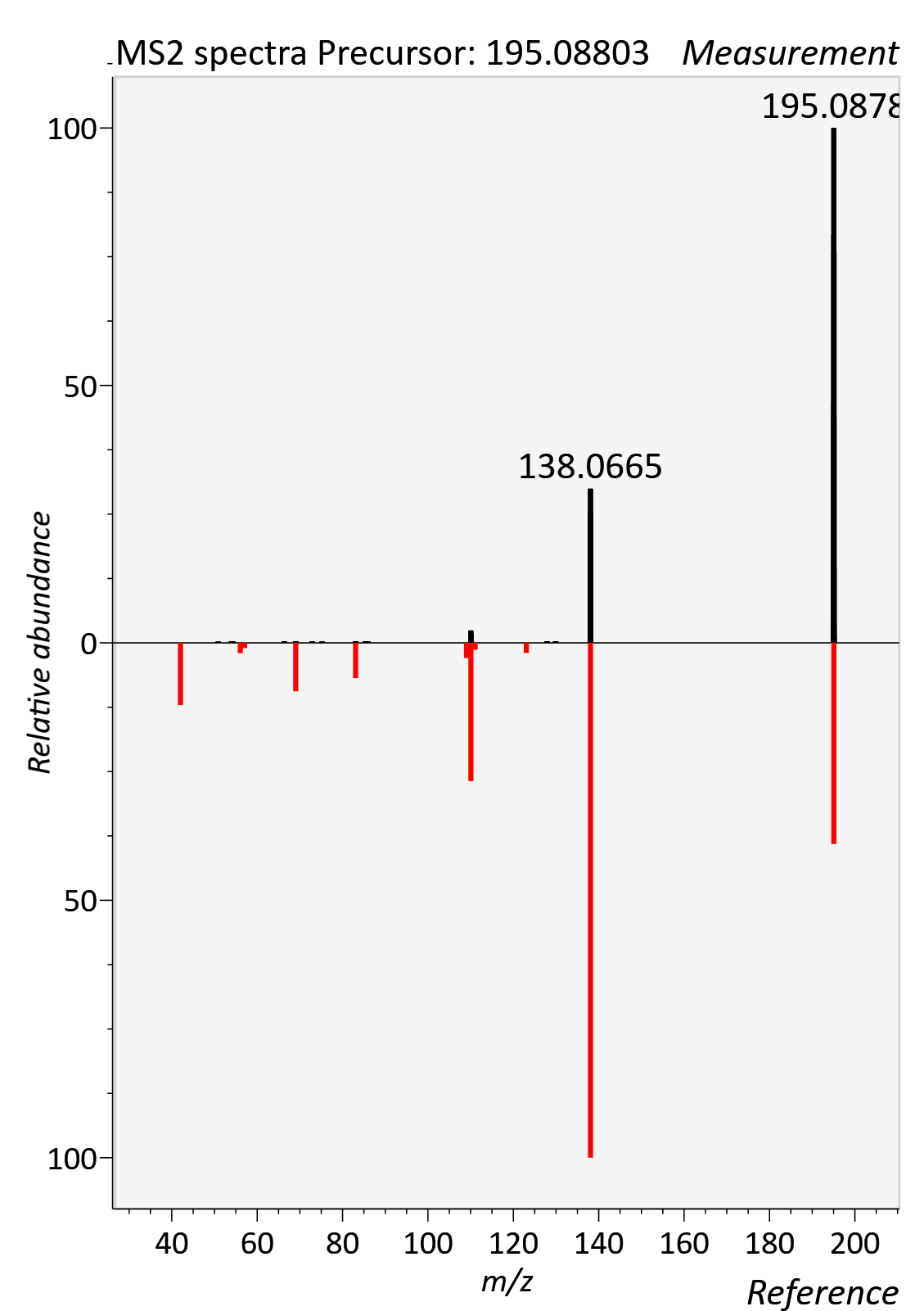

Citrulline

RT std: 6.35min, RT experimental: 6.50min, RT Δ 0.15min

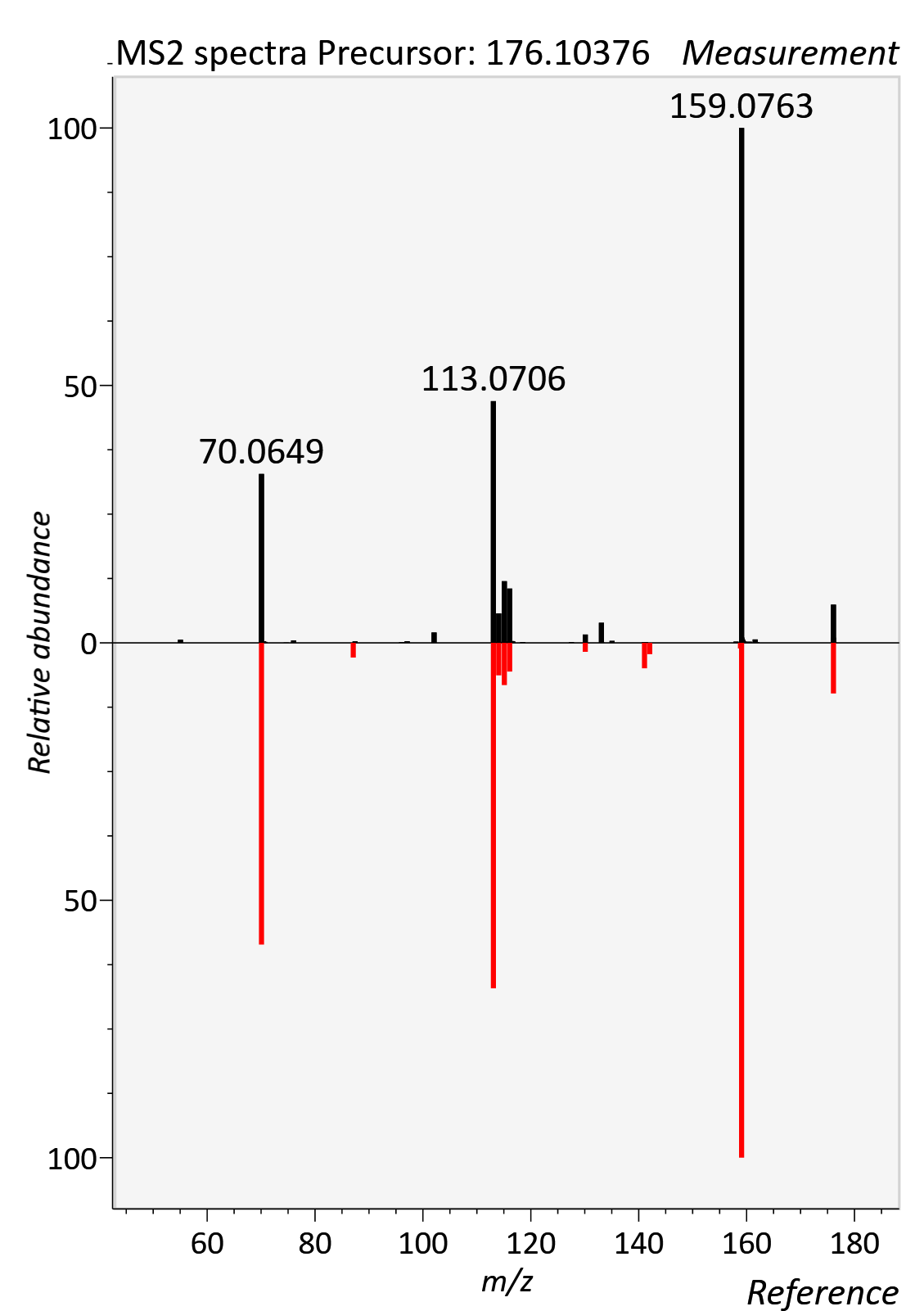

Cholic acid

RT std: 9.08min, RT experimental: 9.28min, RT Δ 0.20min

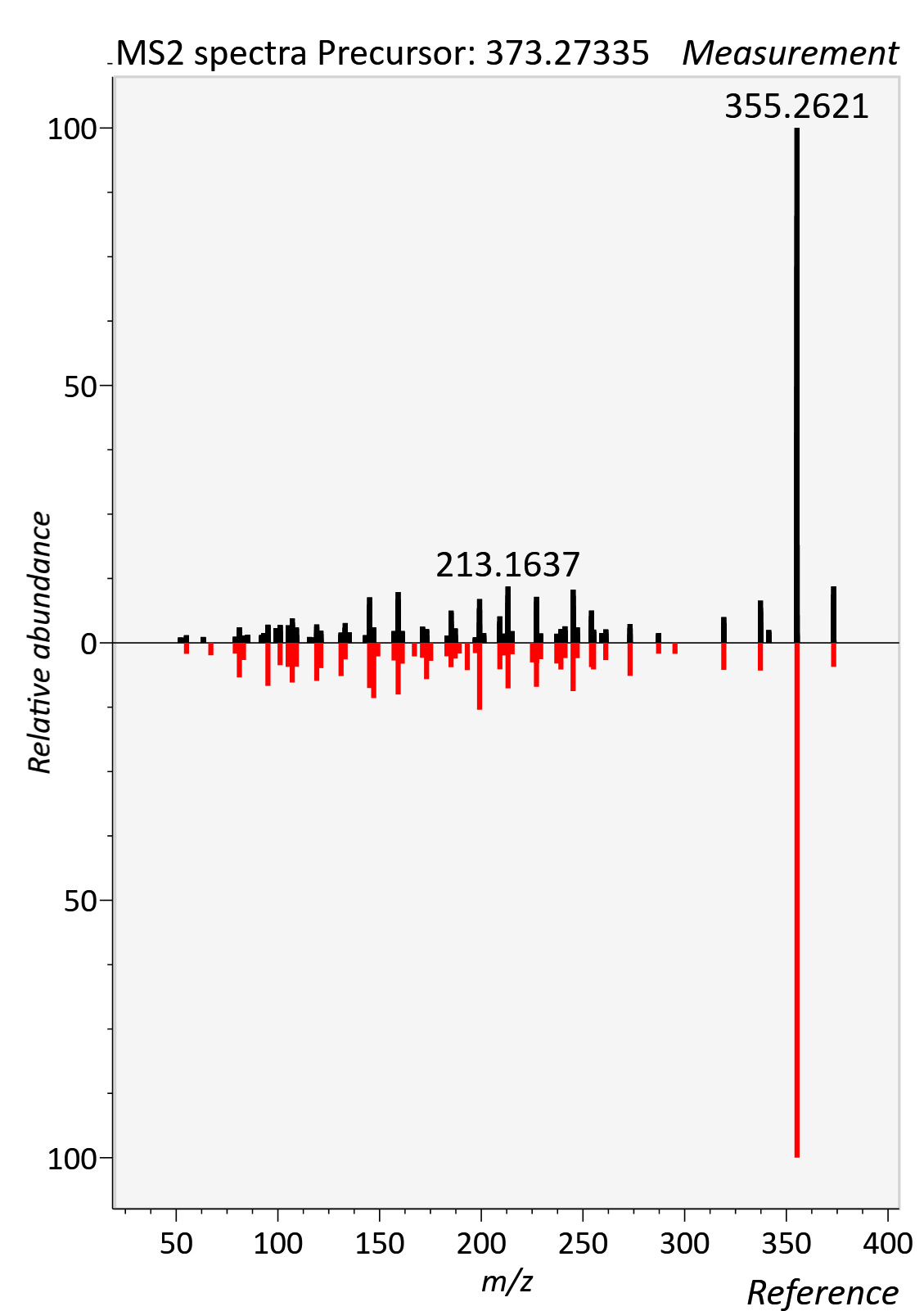

Cortisol

RT std: 6.79min, RT experimental: 6.99min, RT Δ 0.20min

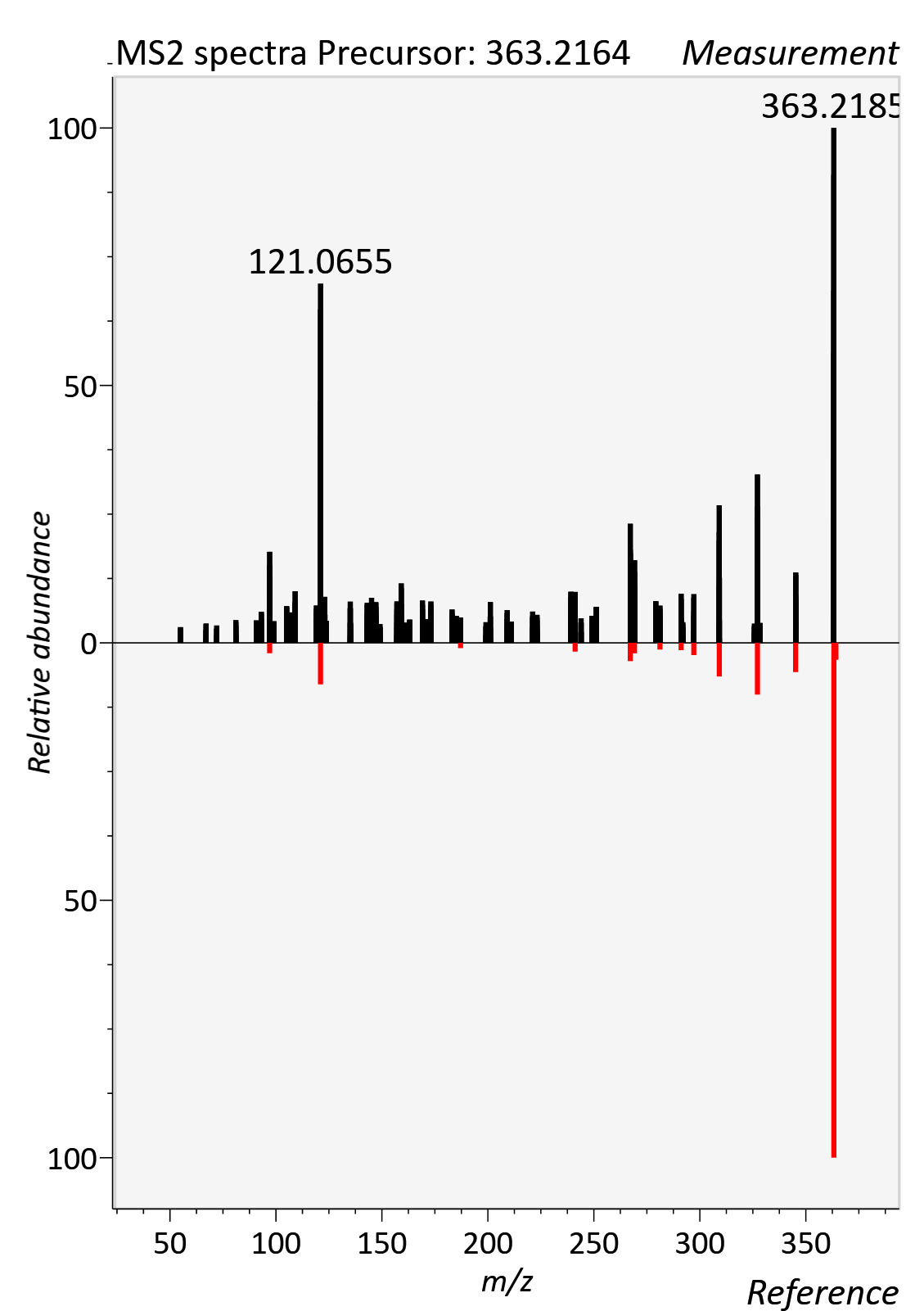

Cortisone, [M+FA-H]-

RT std: 6.55min, RT experimental: 6.73min, RT Δ 0.22min

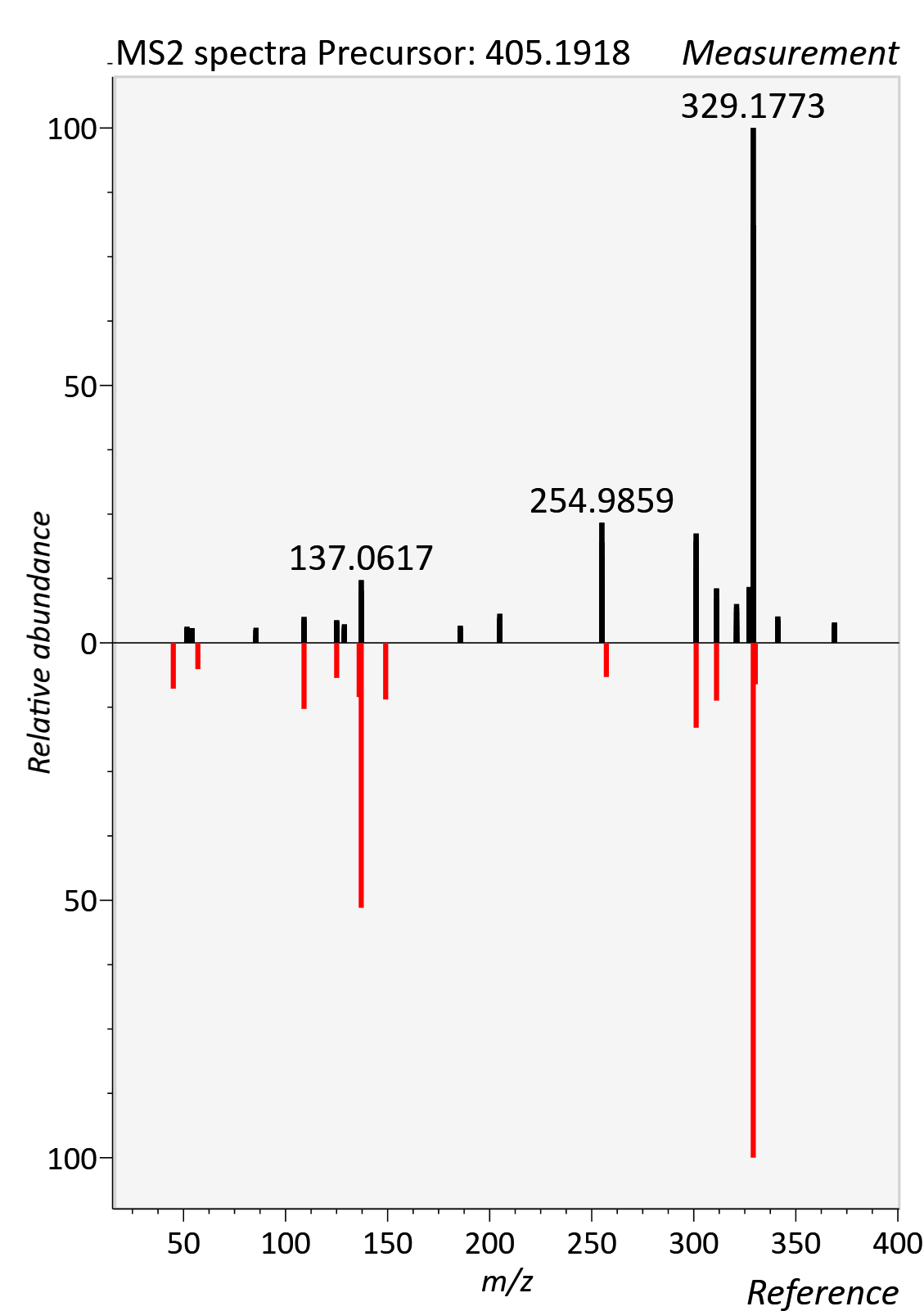

Creatinine

RT std: 1.22min, RT experimental: 1.35min, RT Δ 0.13min

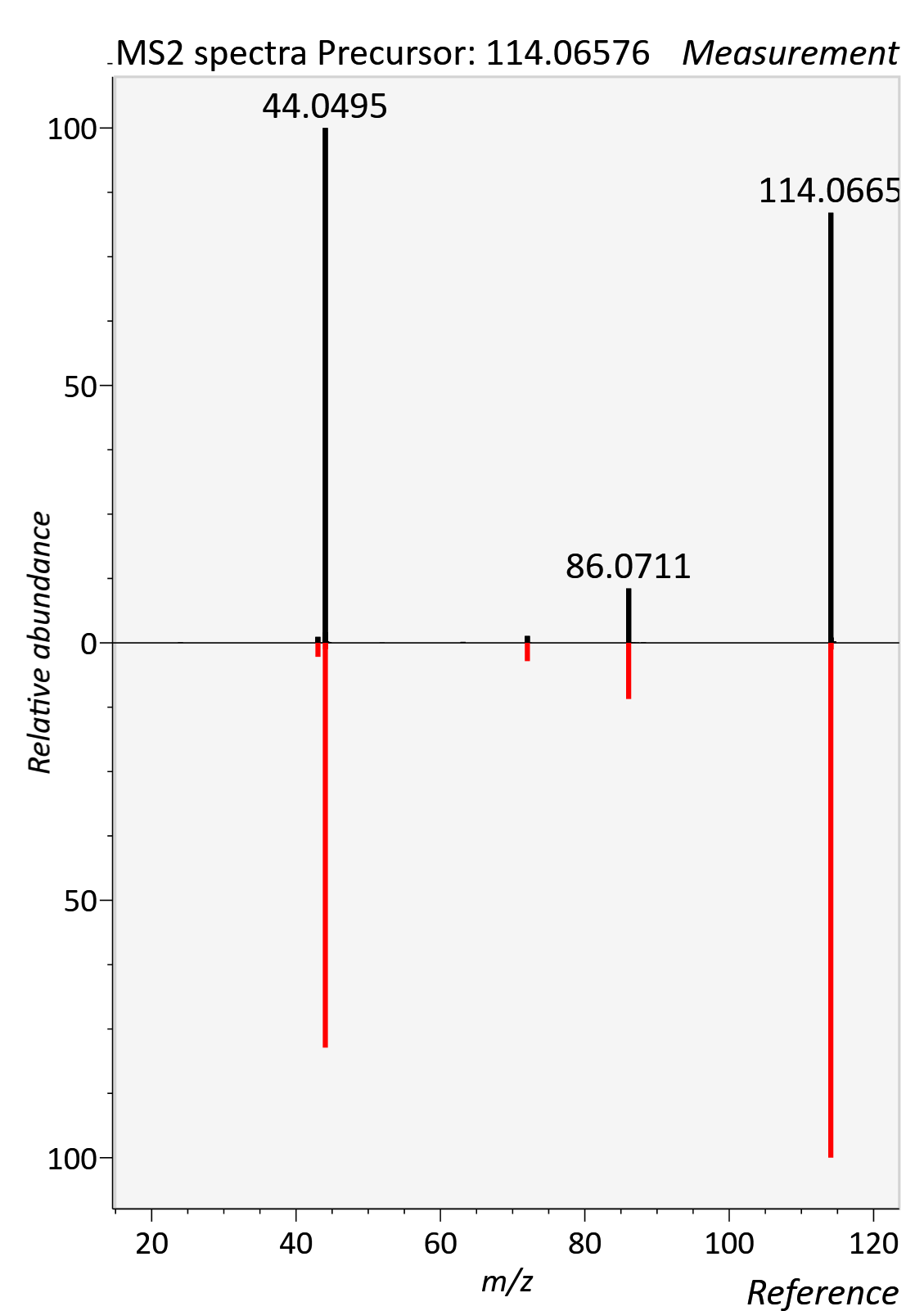

Deoxycholic acid

RT std: 9.71min, RT experimental: 9.90min, RT Δ 0.19min

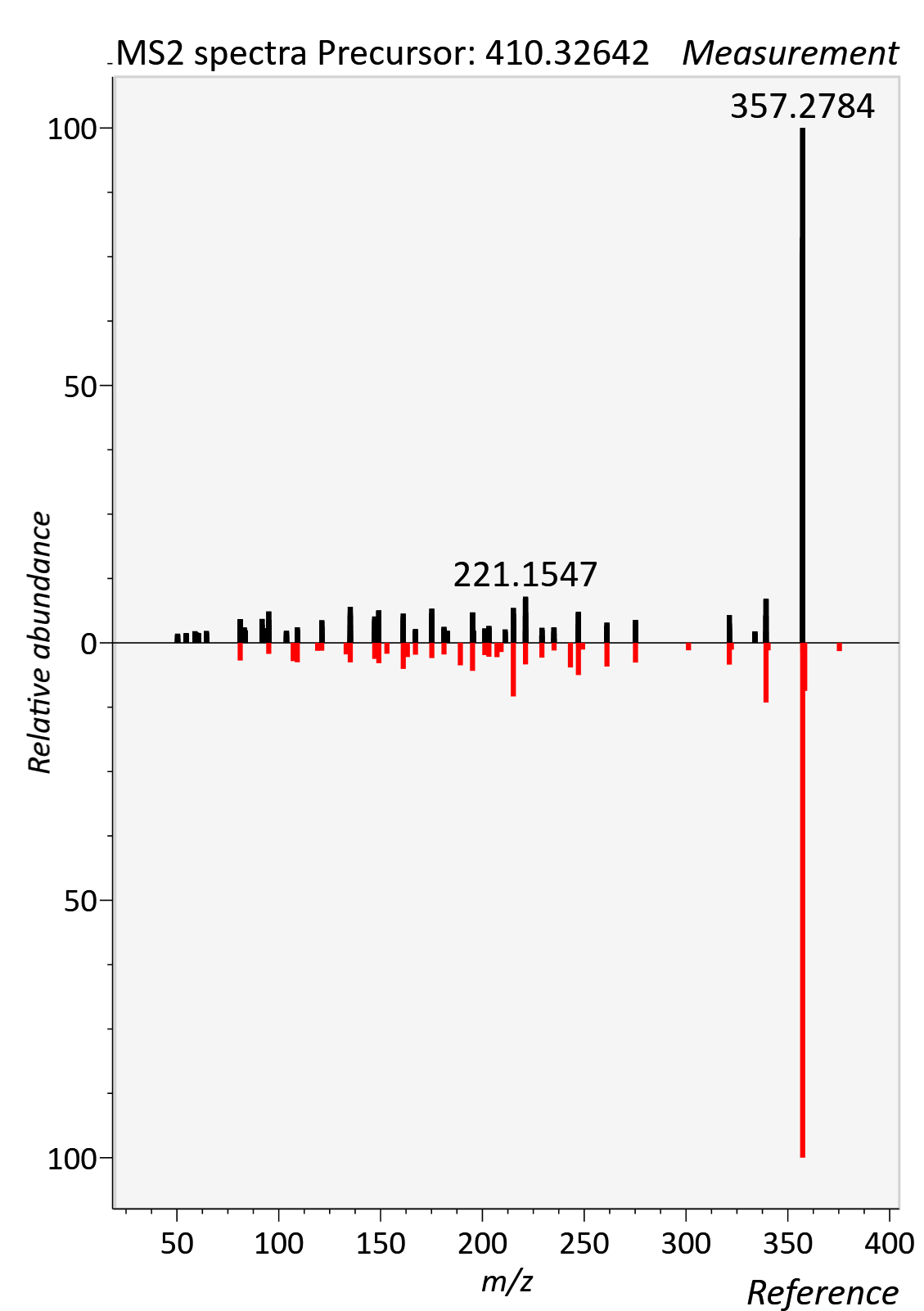

FA 18:1 (Octadecenoic acid)

RT std: 11.23min, RT experimental: 10.92min, RT Δ 0.31min

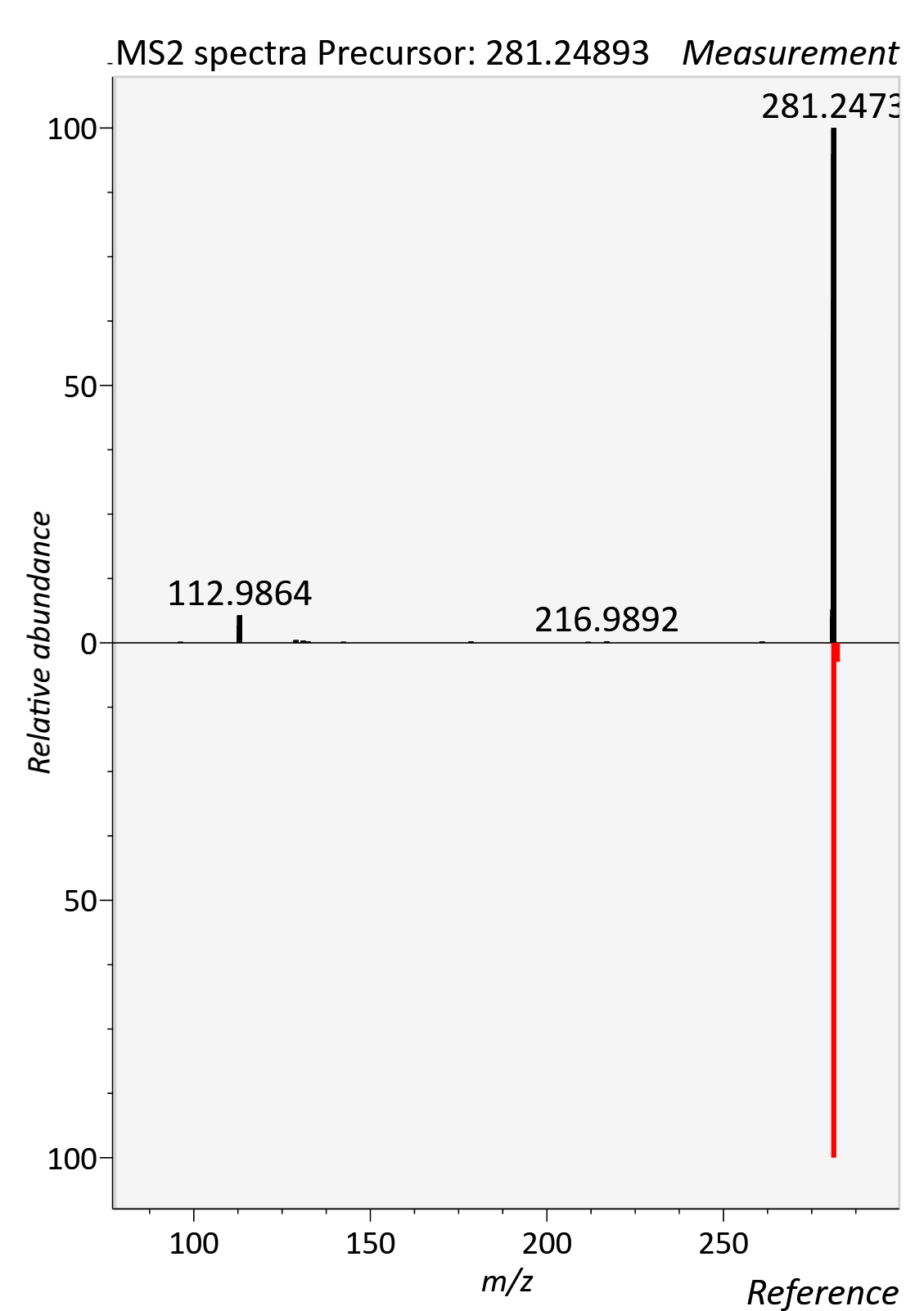

FA 22:6 (DHA)

RT std: 10.66min, RT experimental: 10.89min, RT Δ 0.23min

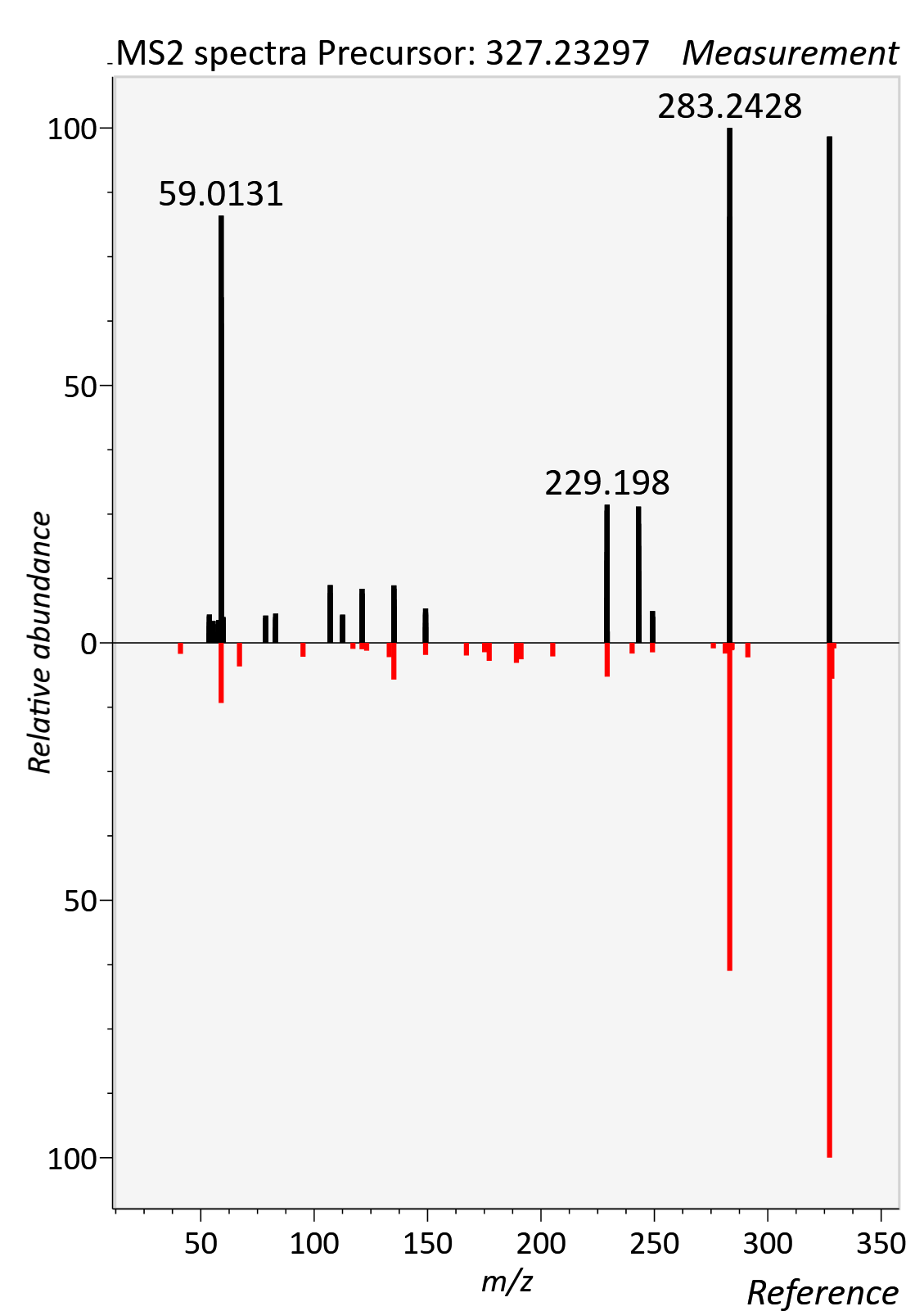

Glutamine

RT std: 6.13min, RT experimental: 6.27min, RT Δ 0.14min

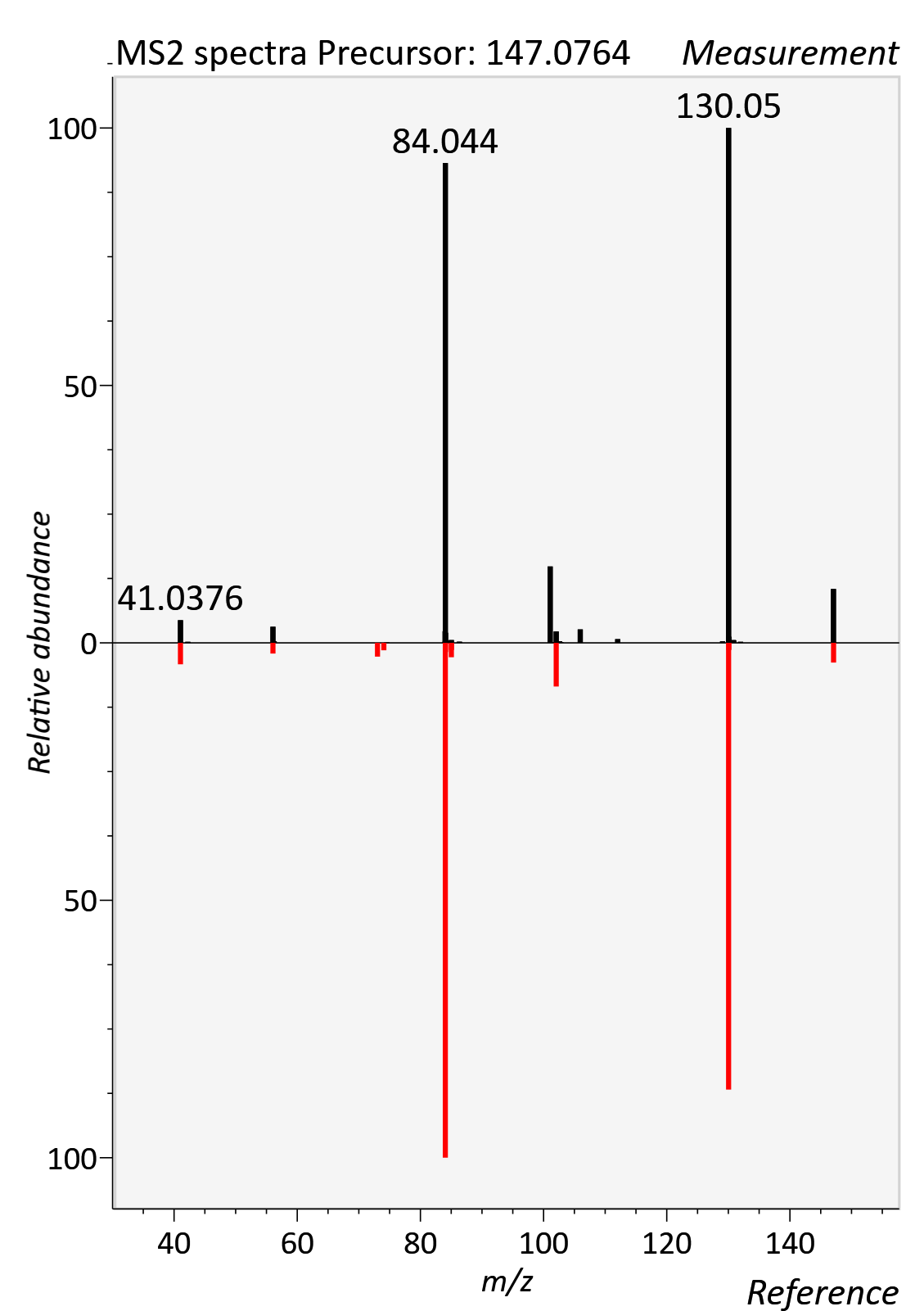

Glycine Betaine

RT std: 3.53min, RT experimental: 4.04min, RT Δ 0.49min

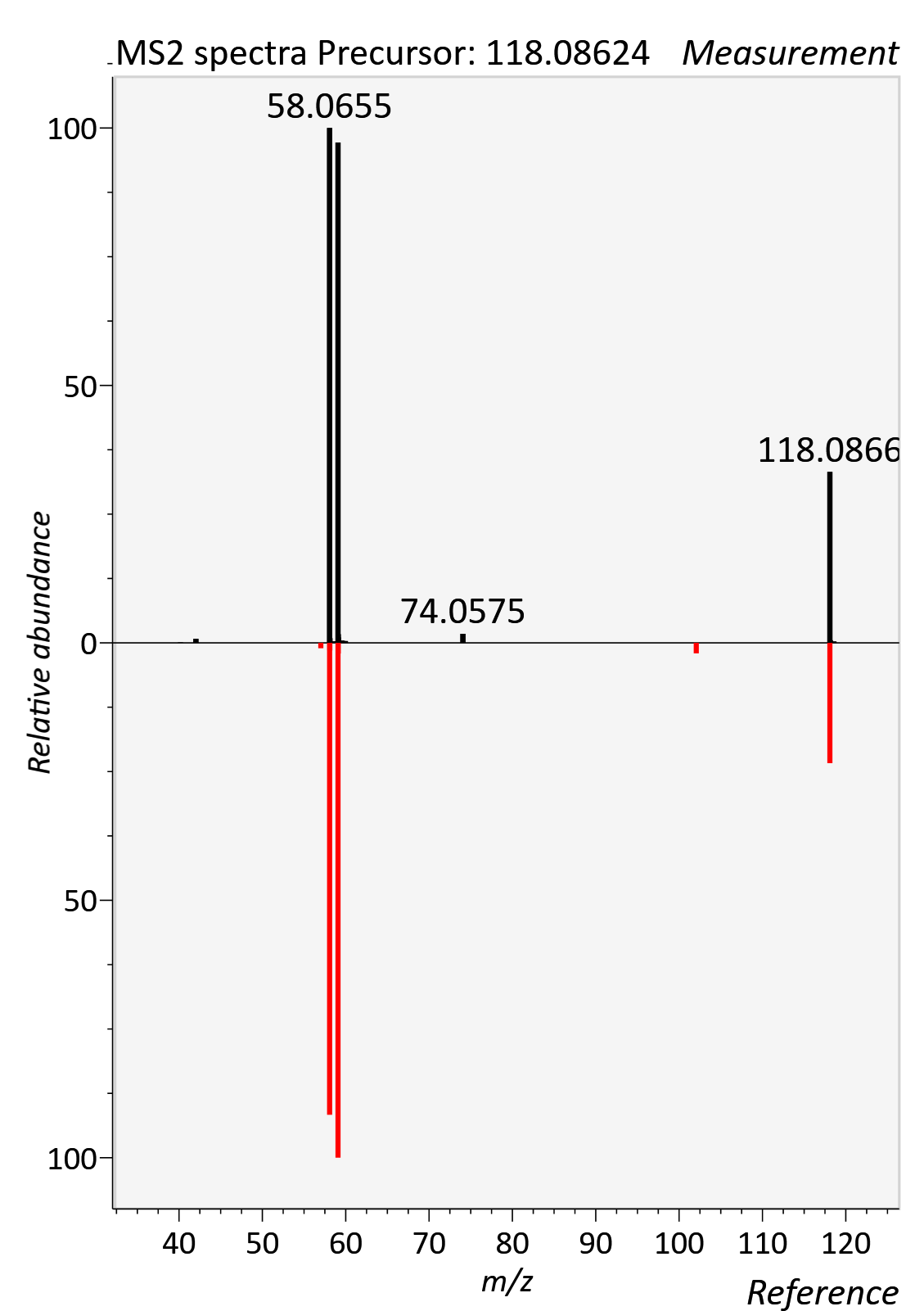

Glycochenodeoxycholic acid

RT std: 9.03min, RT experimental: 9.23min, RT Δ 0.20min

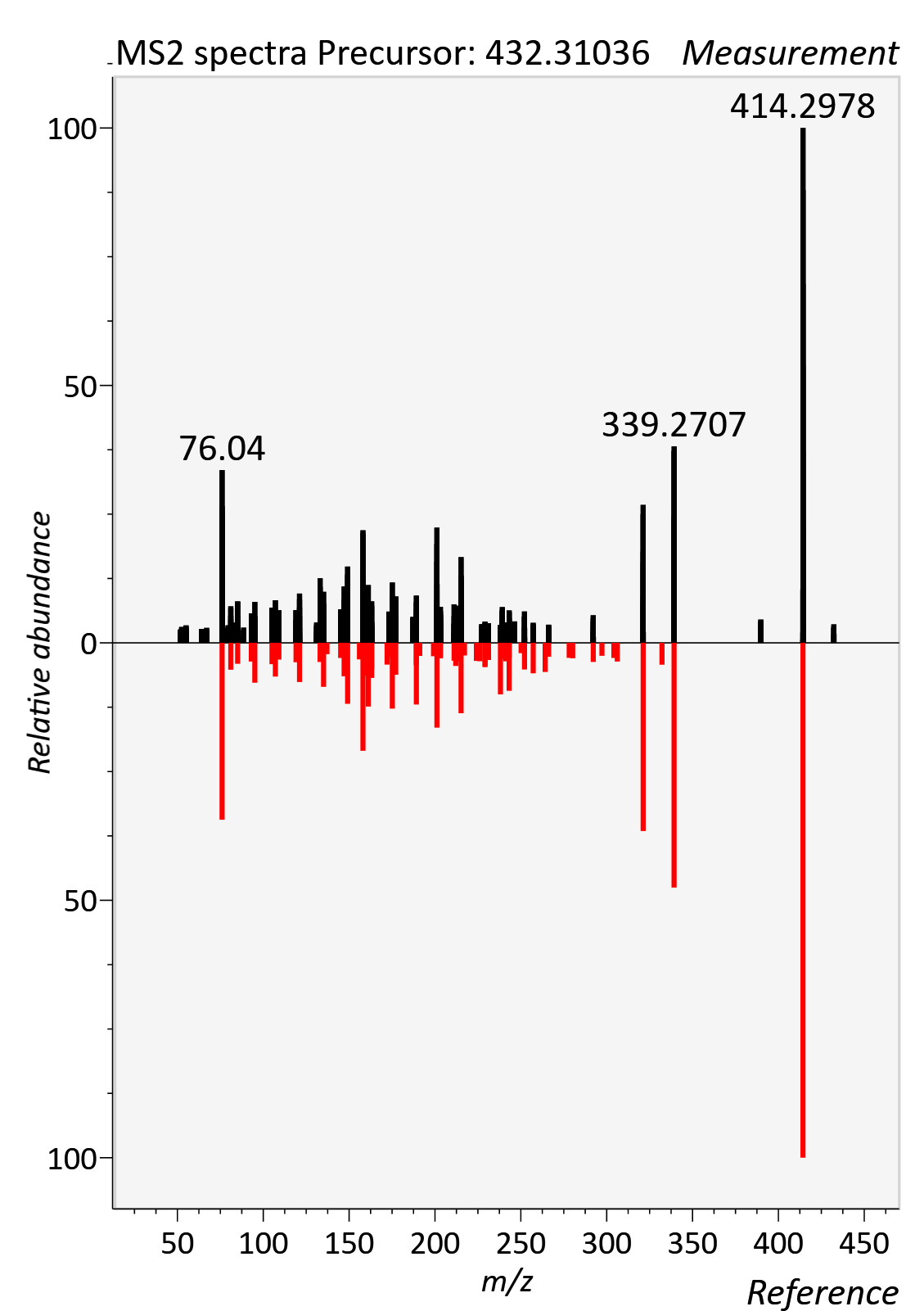

Glycocholic acid

RT std: 8.74min, RT experimental: 8.59min, RT Δ 0.15 min

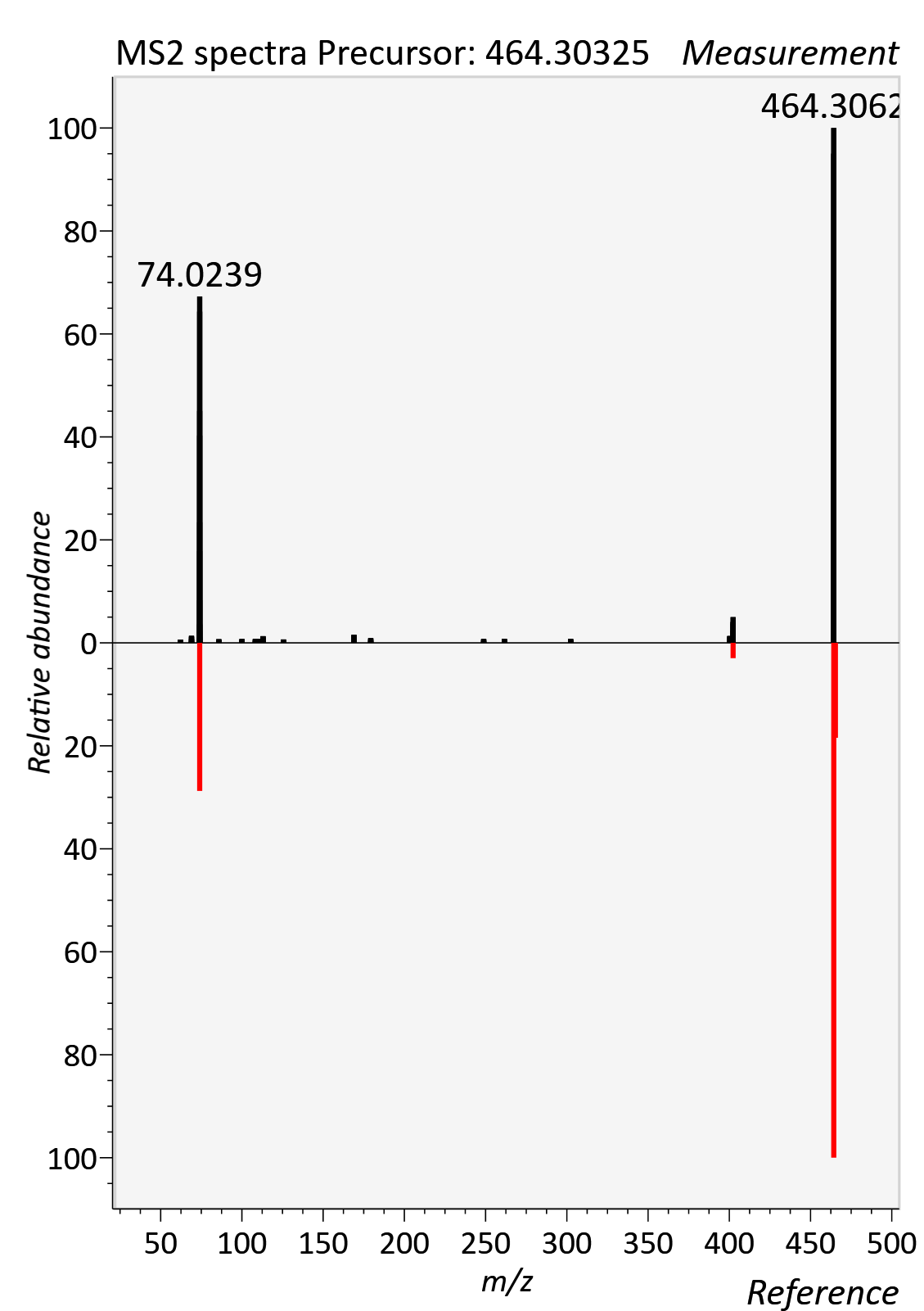

Glycohyodeoxycholic acid

RT std: 8.24min, RT experimental: 8.25min, RT Δ 0.01min

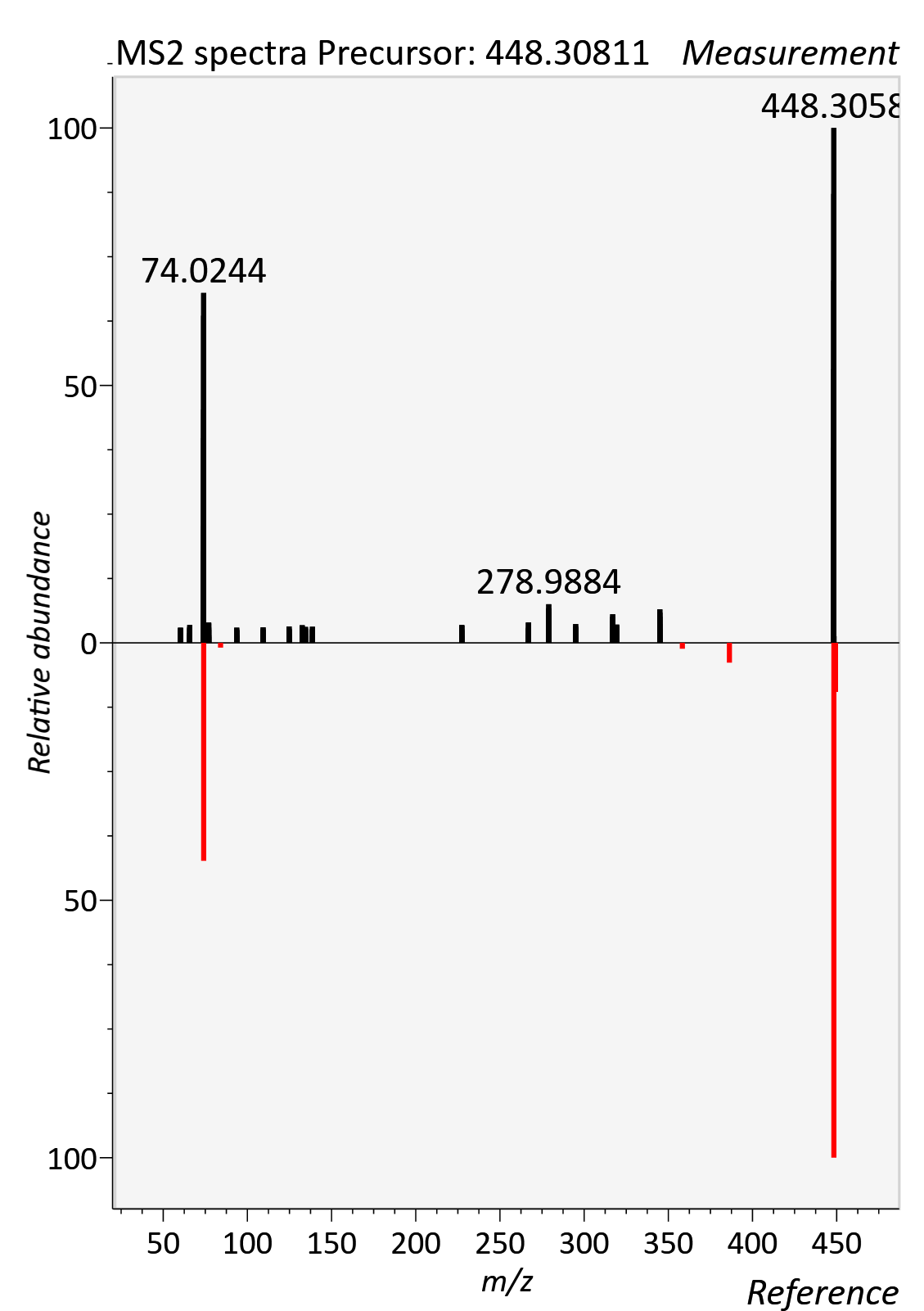

Glycolithocholic aicd

RT std: 9.33min, RT experimental: 9.58min, RT Δ 0.25min

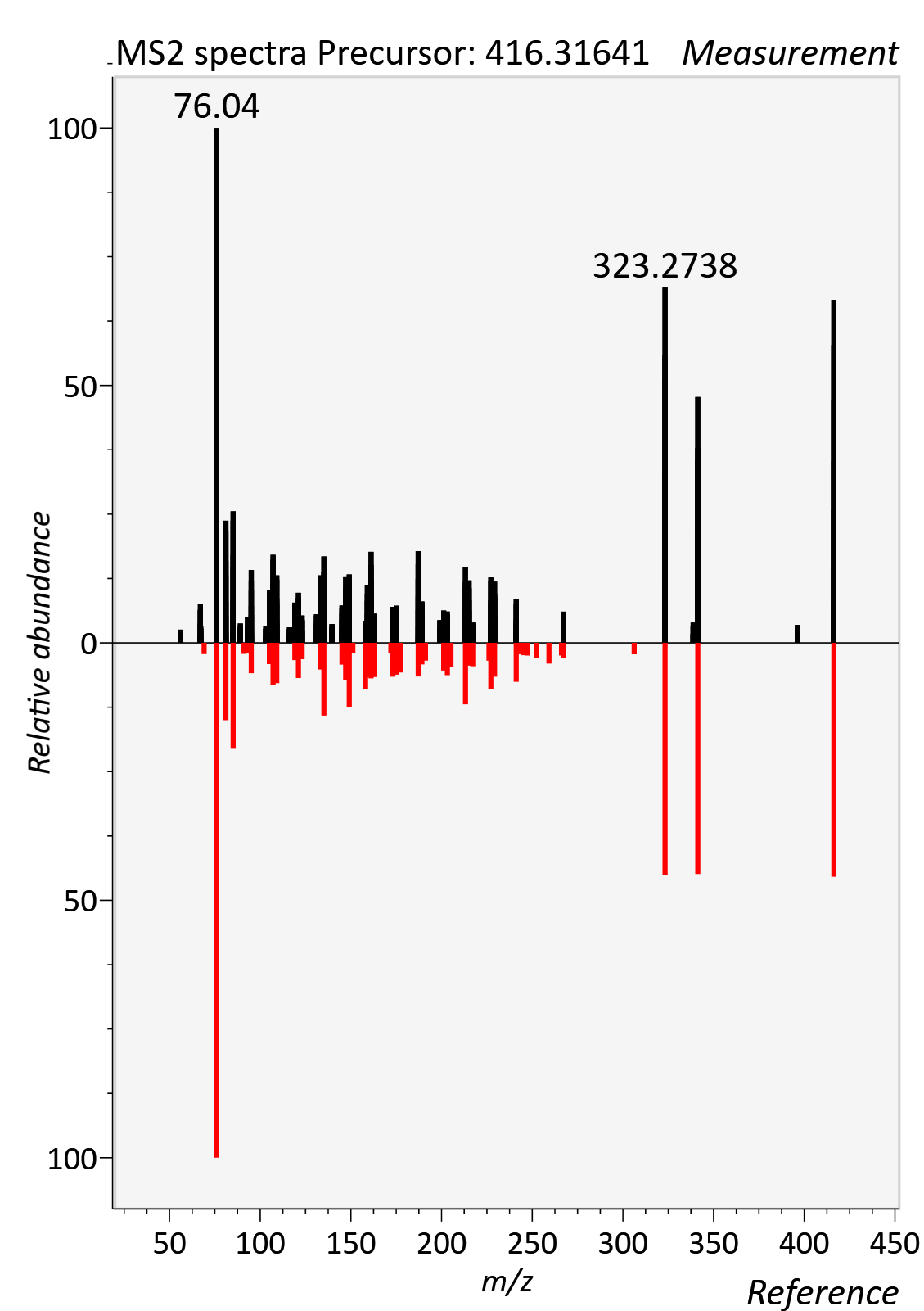

Hippuric acid

RT std: 3.30min, RT experimental: 3.54min, RT Δ 0.24min

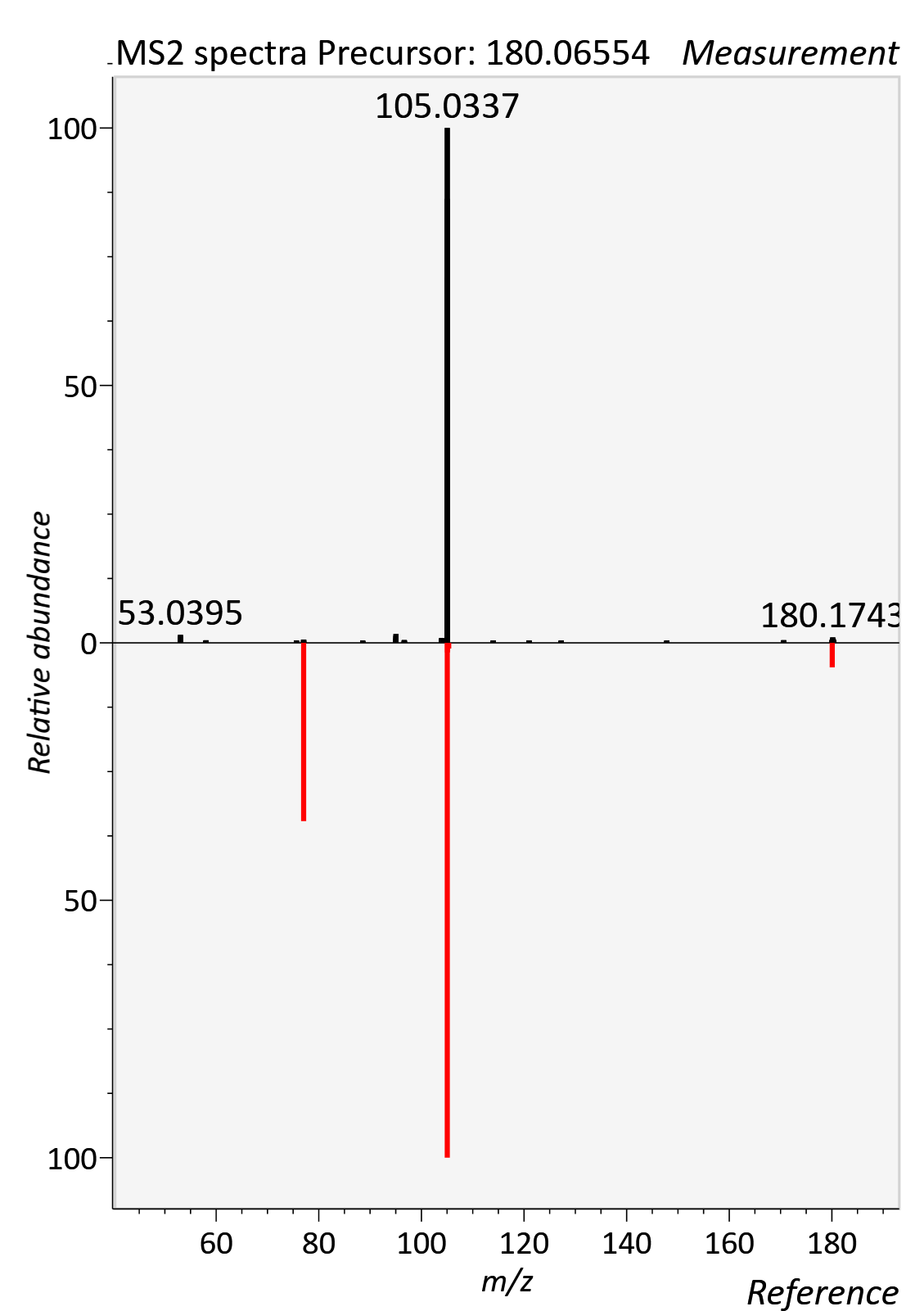

Hydroxyphenyllactic acid

RT std: 2.73min, RT experimental: 3.03min, RT Δ 0.30min

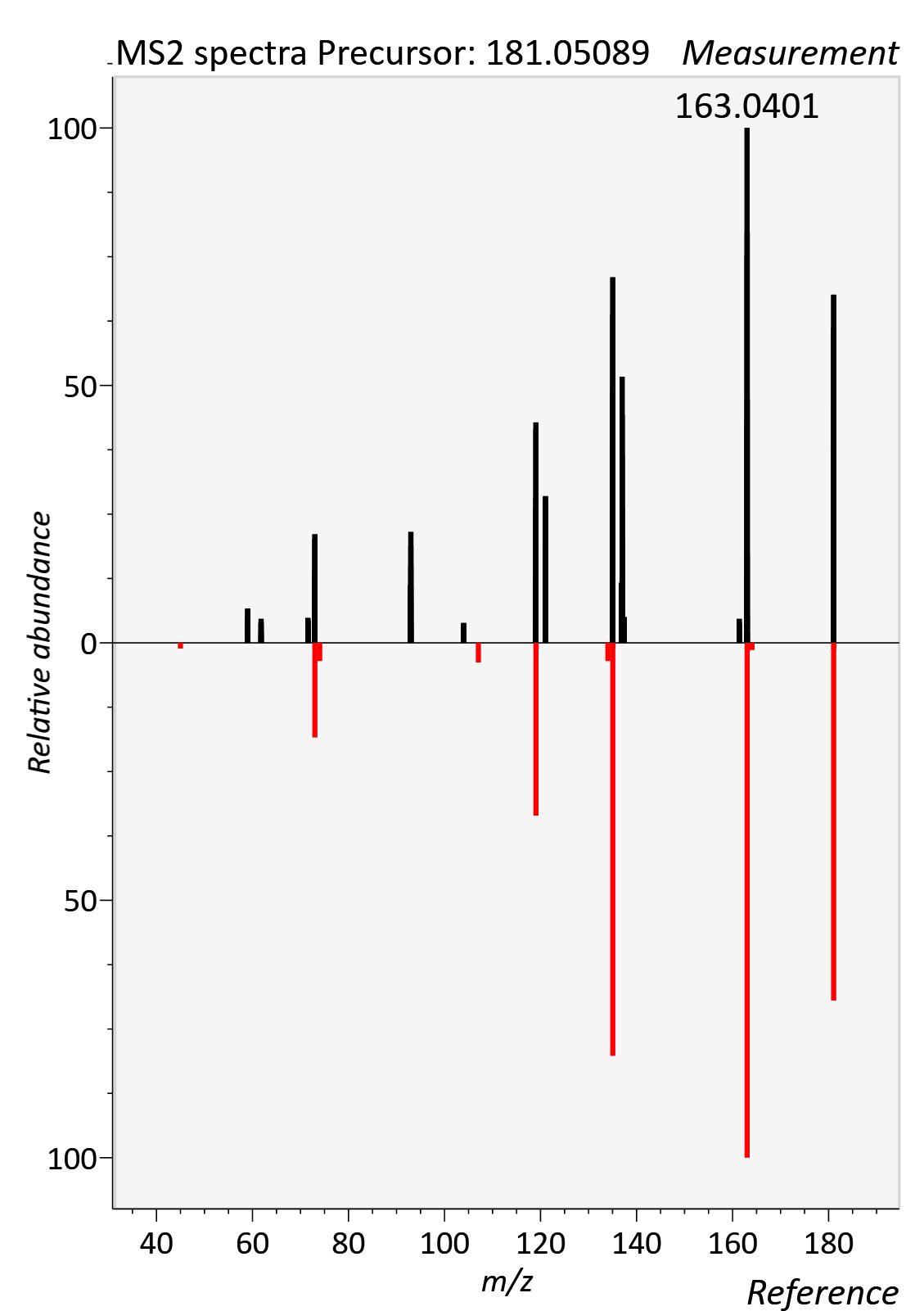

Hypoxanthine

RT std: 1.63min, RT experimental: 1.55min, RT Δ 0.08min

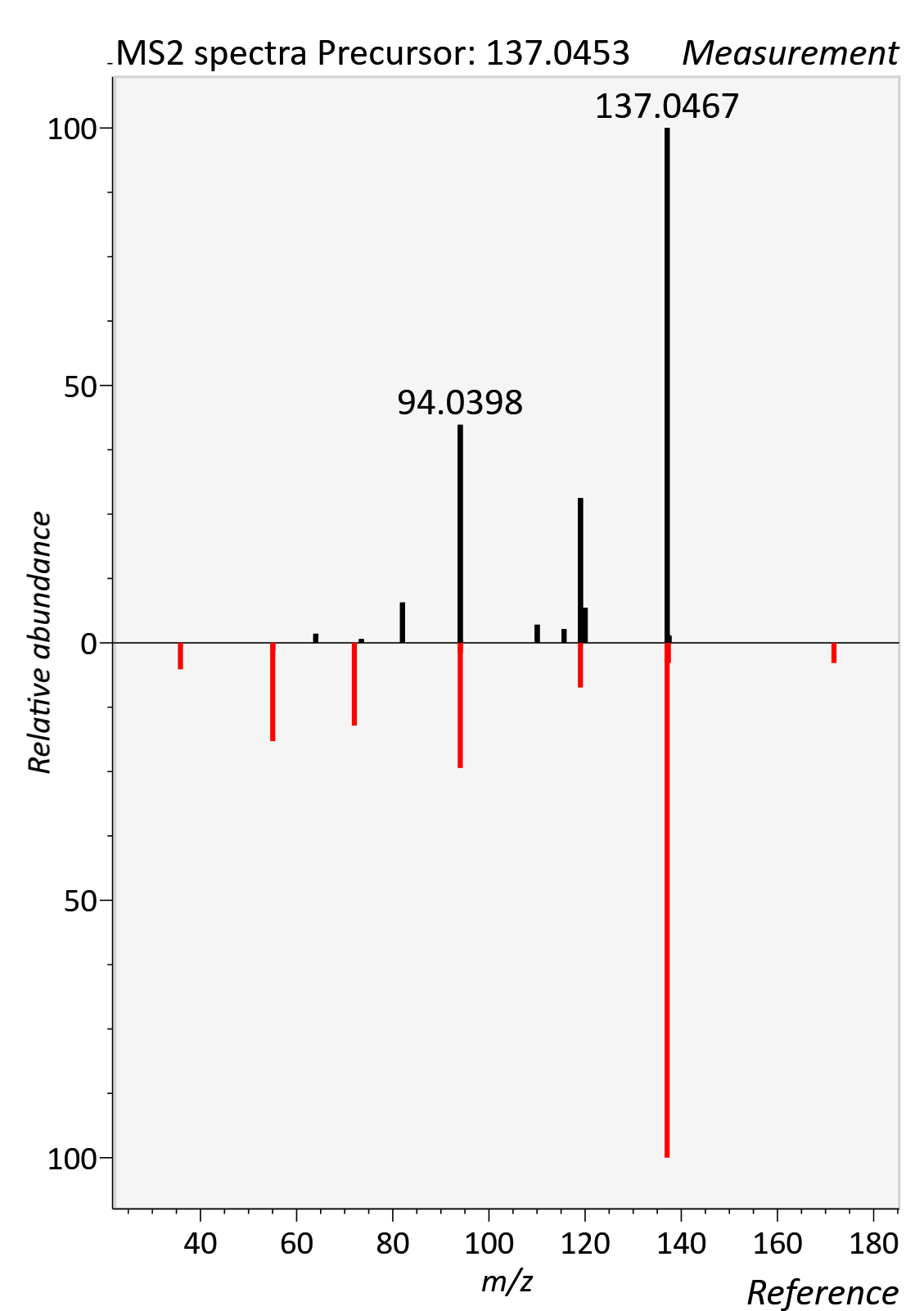

Indolelactic acid

RT std: 4.29min, RT experimental: 4.82min, RT Δ 0.53min

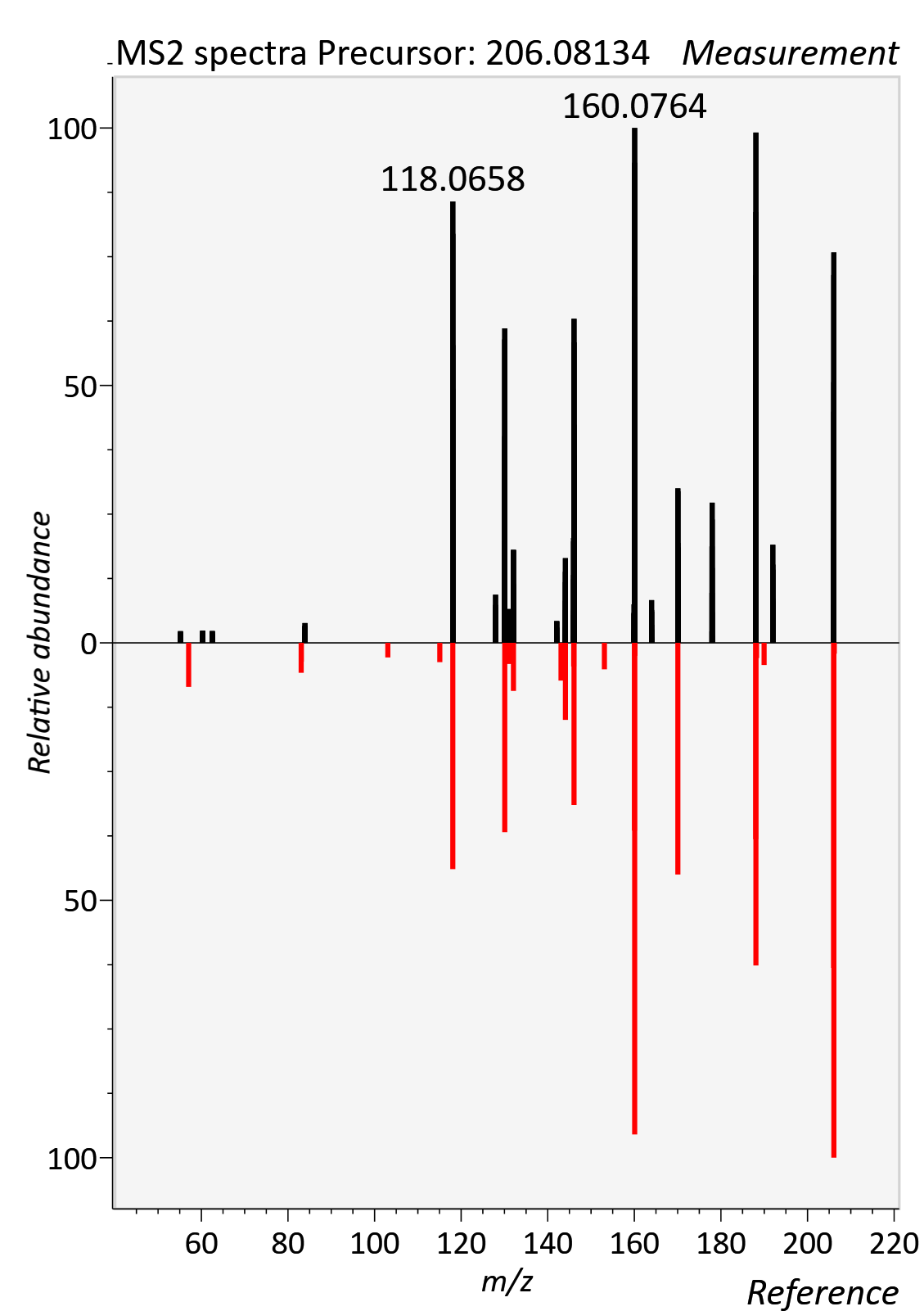

Indoxyl sulfate

RT std: 2.64min, RT experimental: 2.73min, RT Δ 0.09min

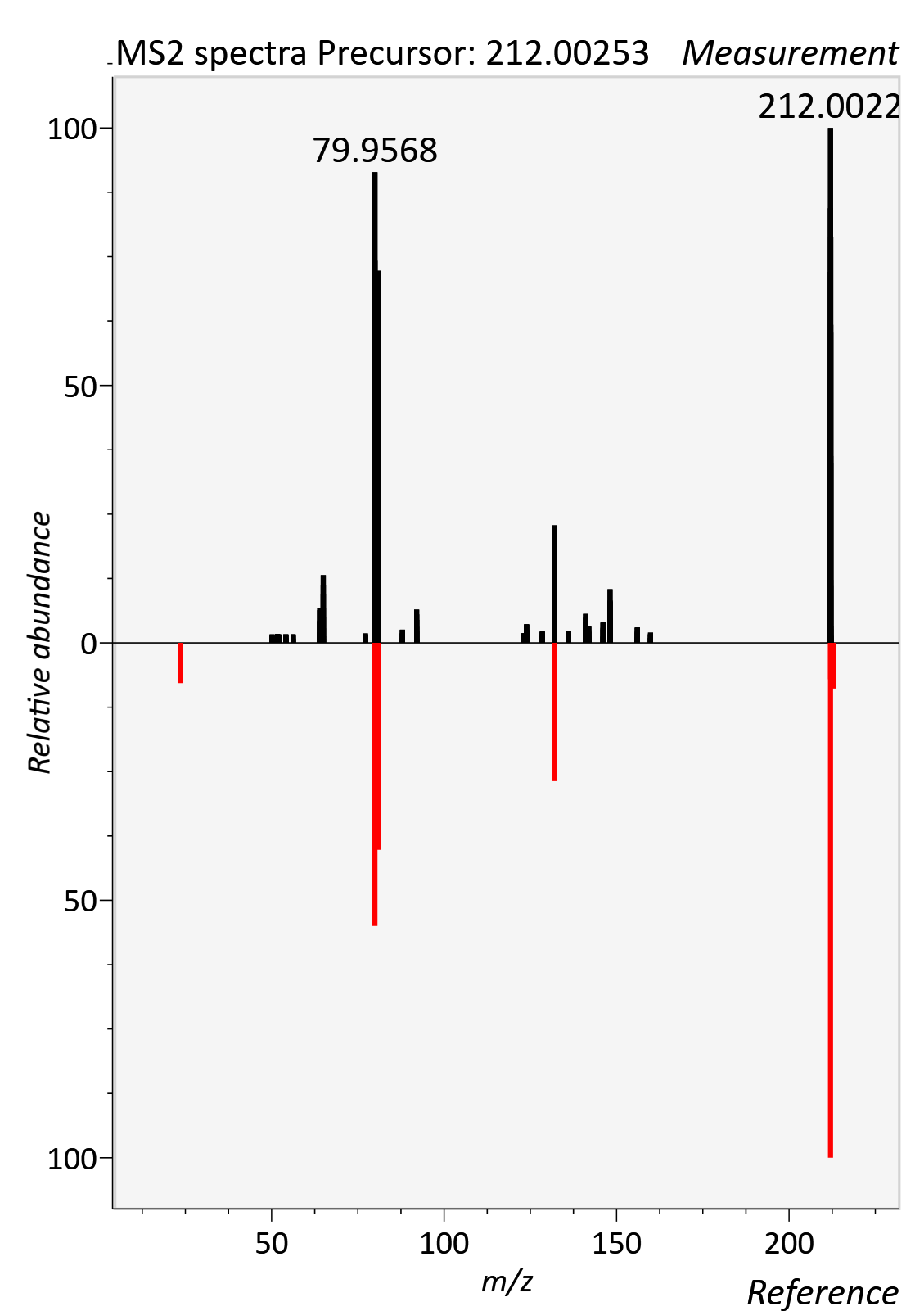

Isoleucine

RT std: 4.14min, RT experimental: 4.53min, RT Δ 0.39min

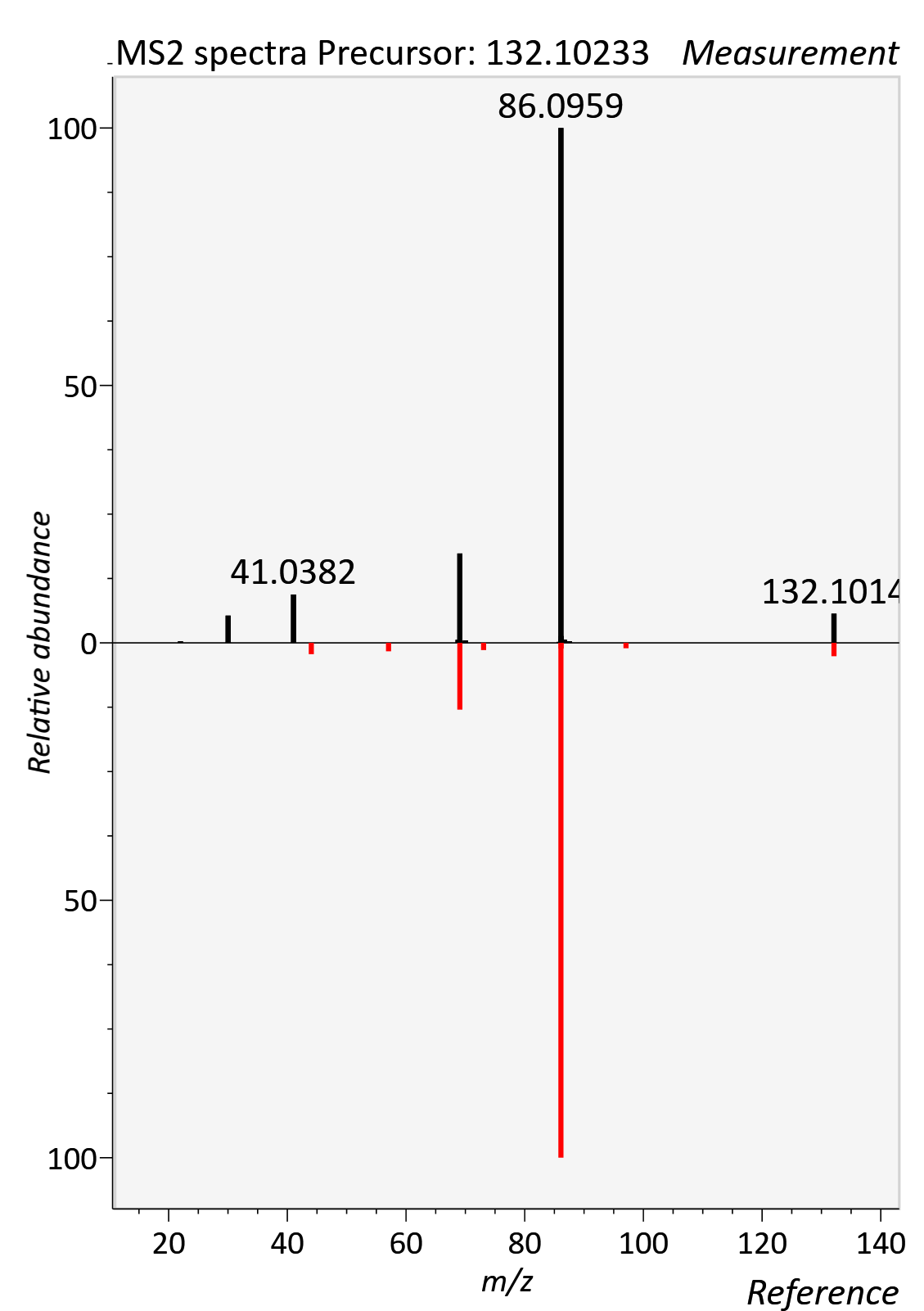

L-Arginine

RT std: 6.94min, RT experimental: 7.11min, RT Δ 0.17min

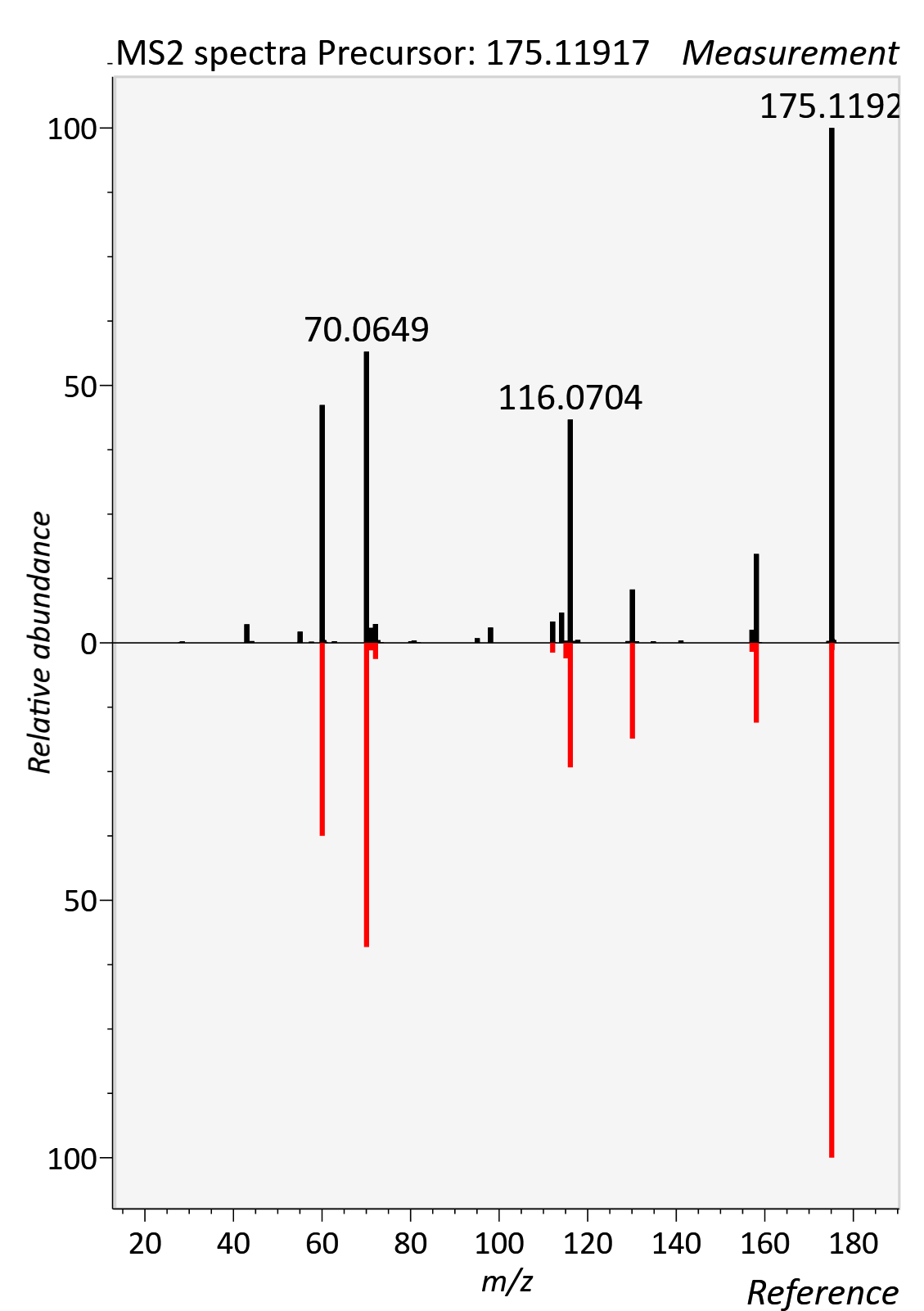

L-Carnitine

RT std: 4.77min, RT experimental: 5.10min, RT Δ 0.37min

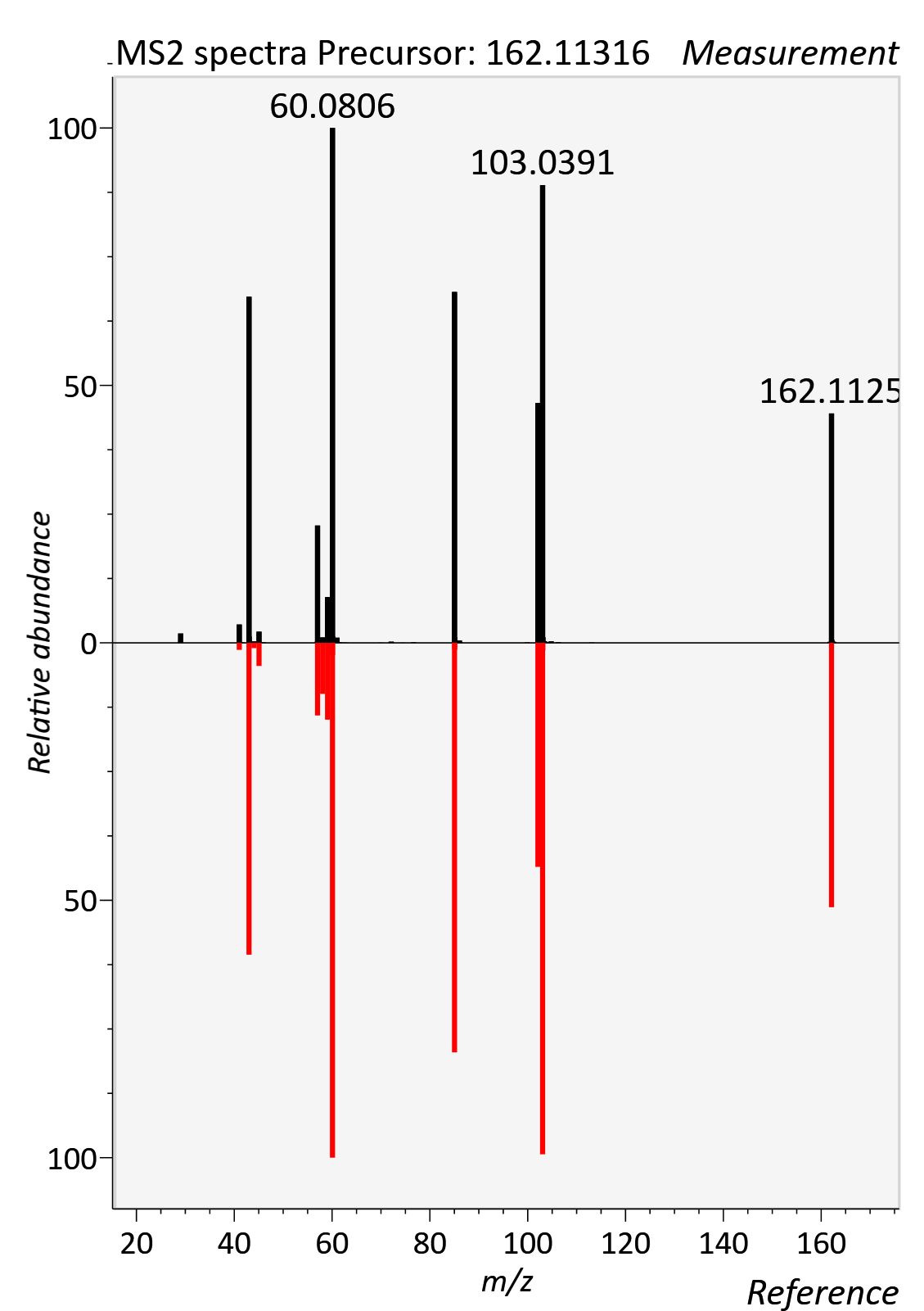

Leucine

RT std: 3.80, RT experimental: 4.28min, RT Δ 0.48min

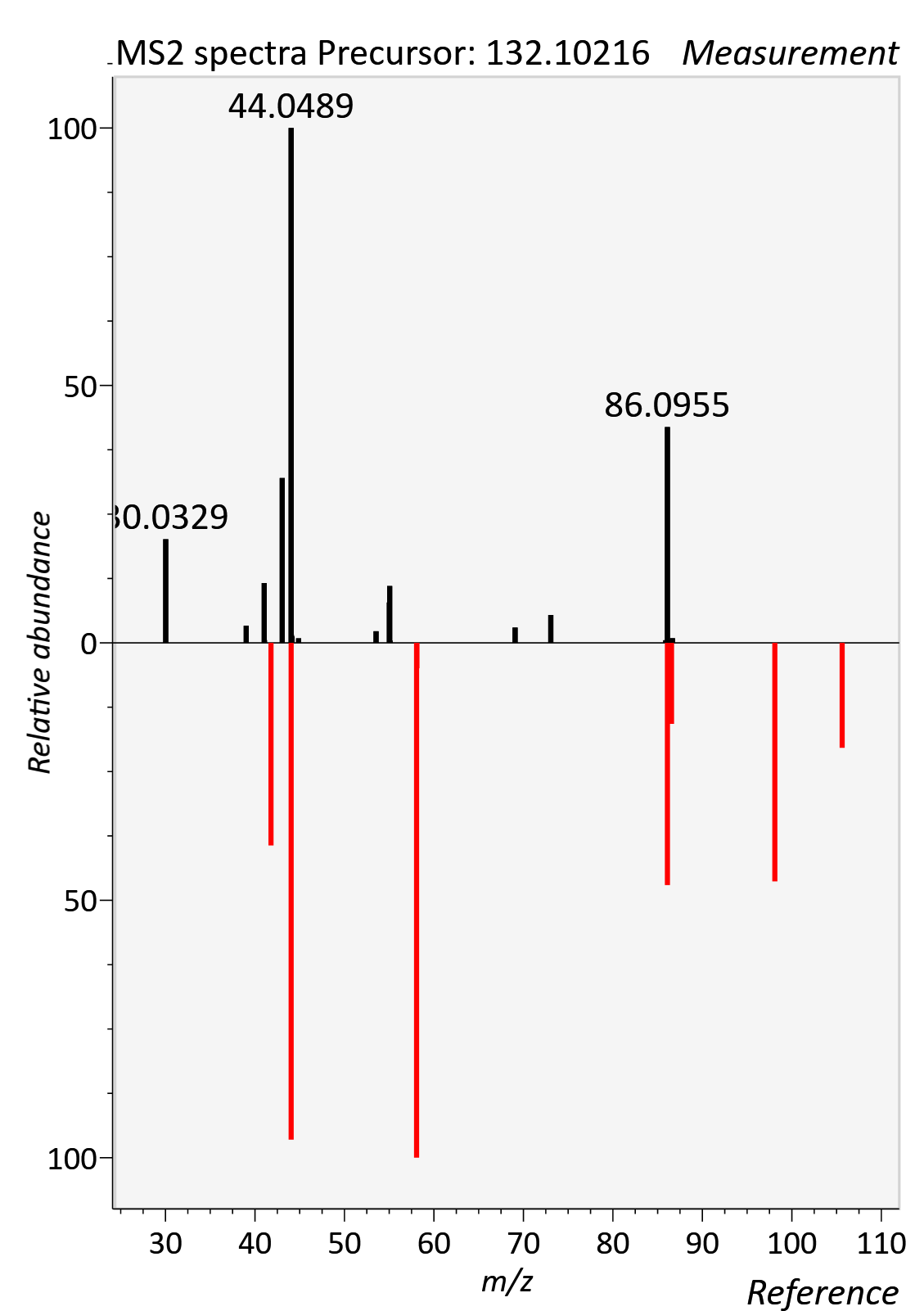

LysoPC 15:0_0:0

RT std: 10.04min, RT experimental: 10.21min, RT Δ 0.16min

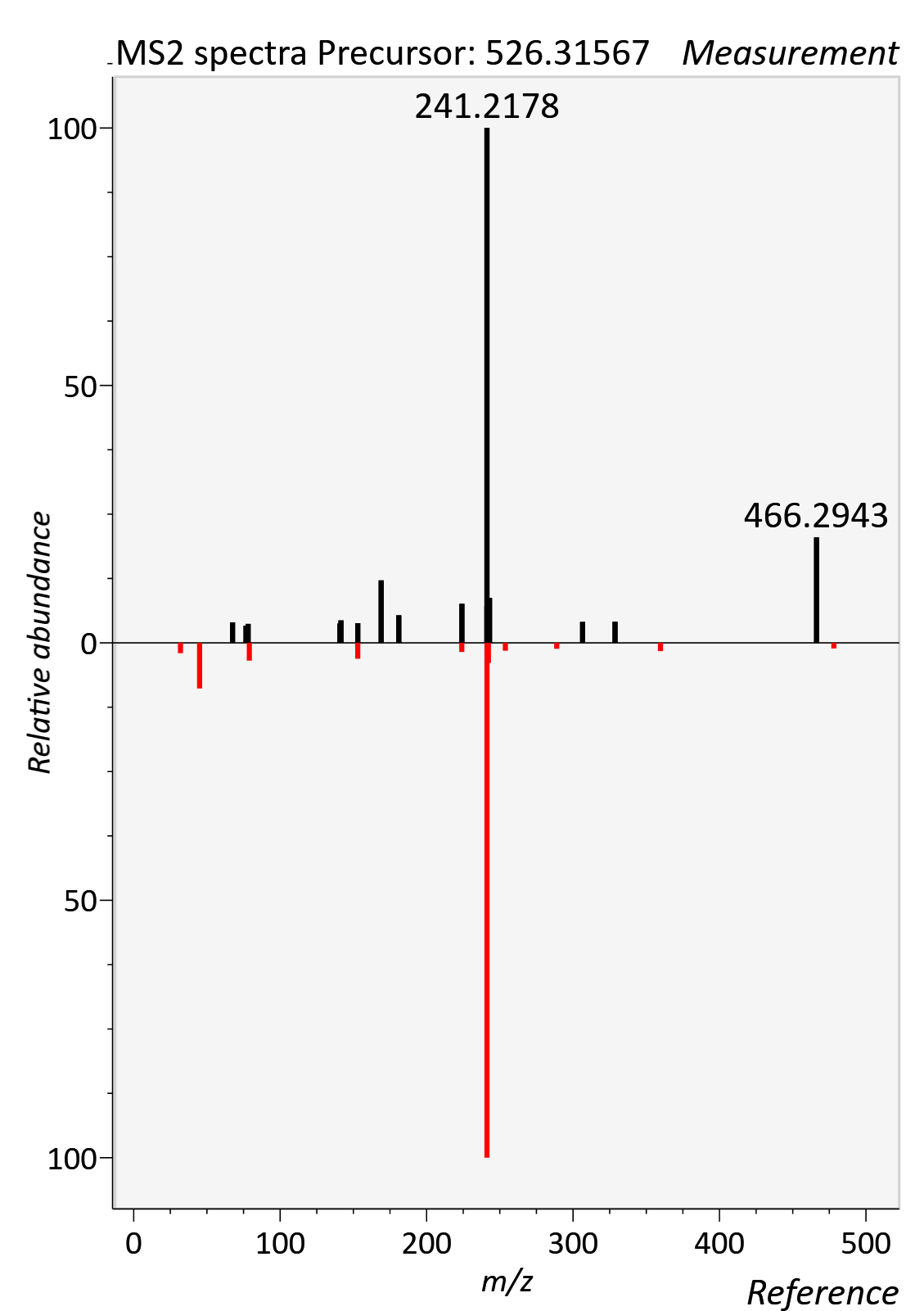

LysoPC 17:0_0:0

RT std: 10.42min, RT experimental: 10.70min, RT Δ 0.28min

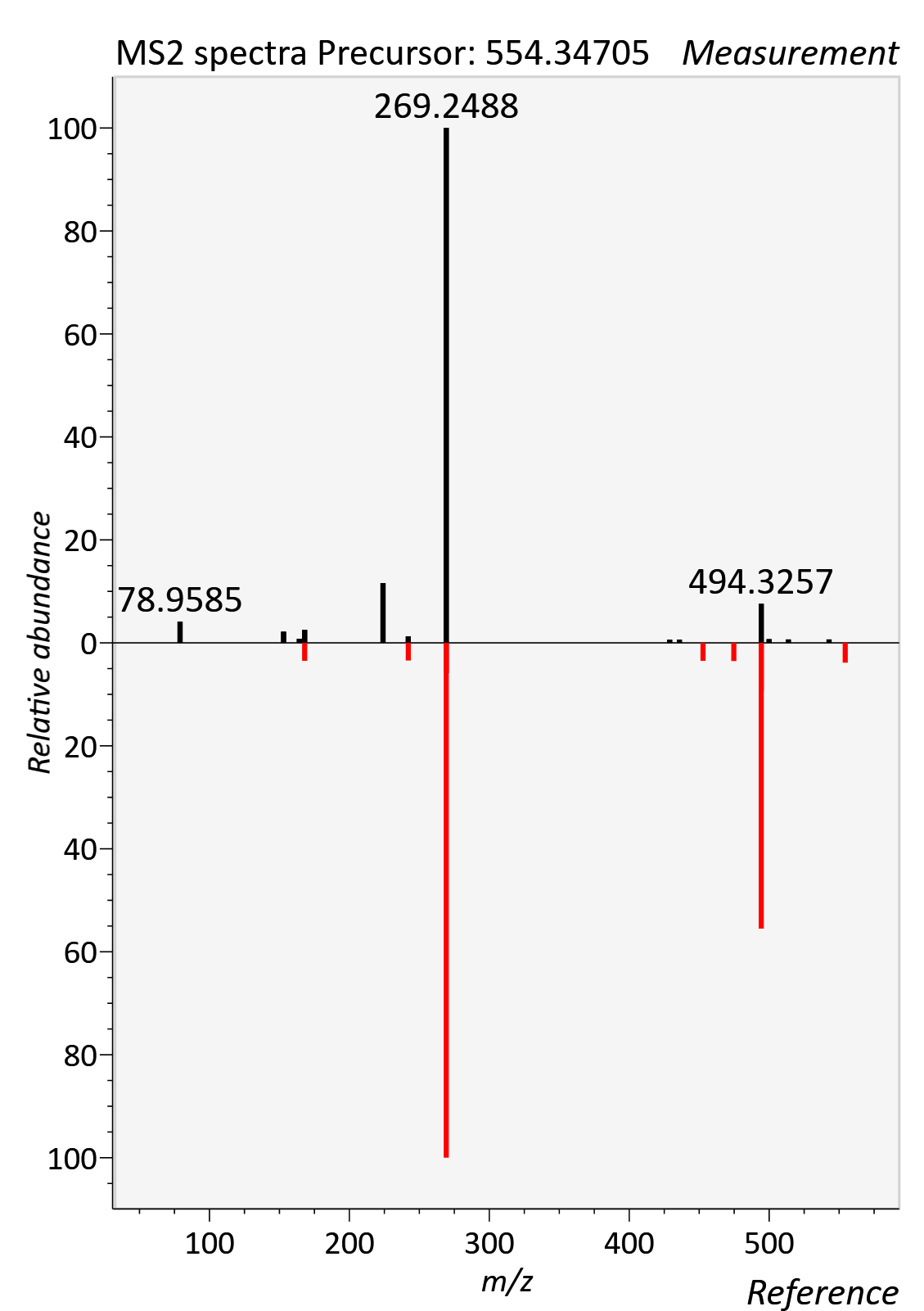

Kynurenine

RT std: 1.73min, RT experimental: 1.90min, RT Δ 0.17min

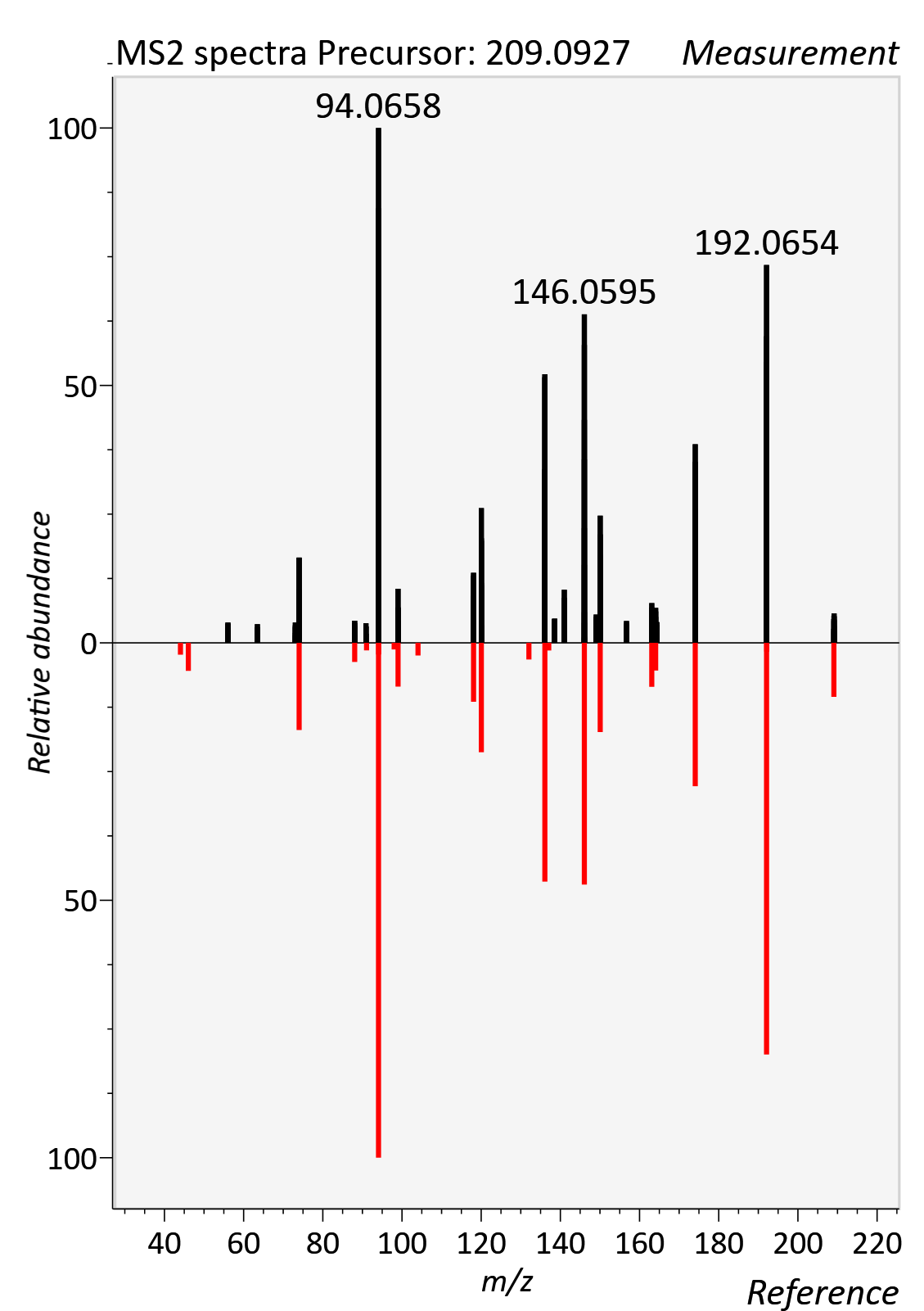

Ornithine

RT std: 7.17min, RT experimental: 7.30min, RT Δ 0.13min

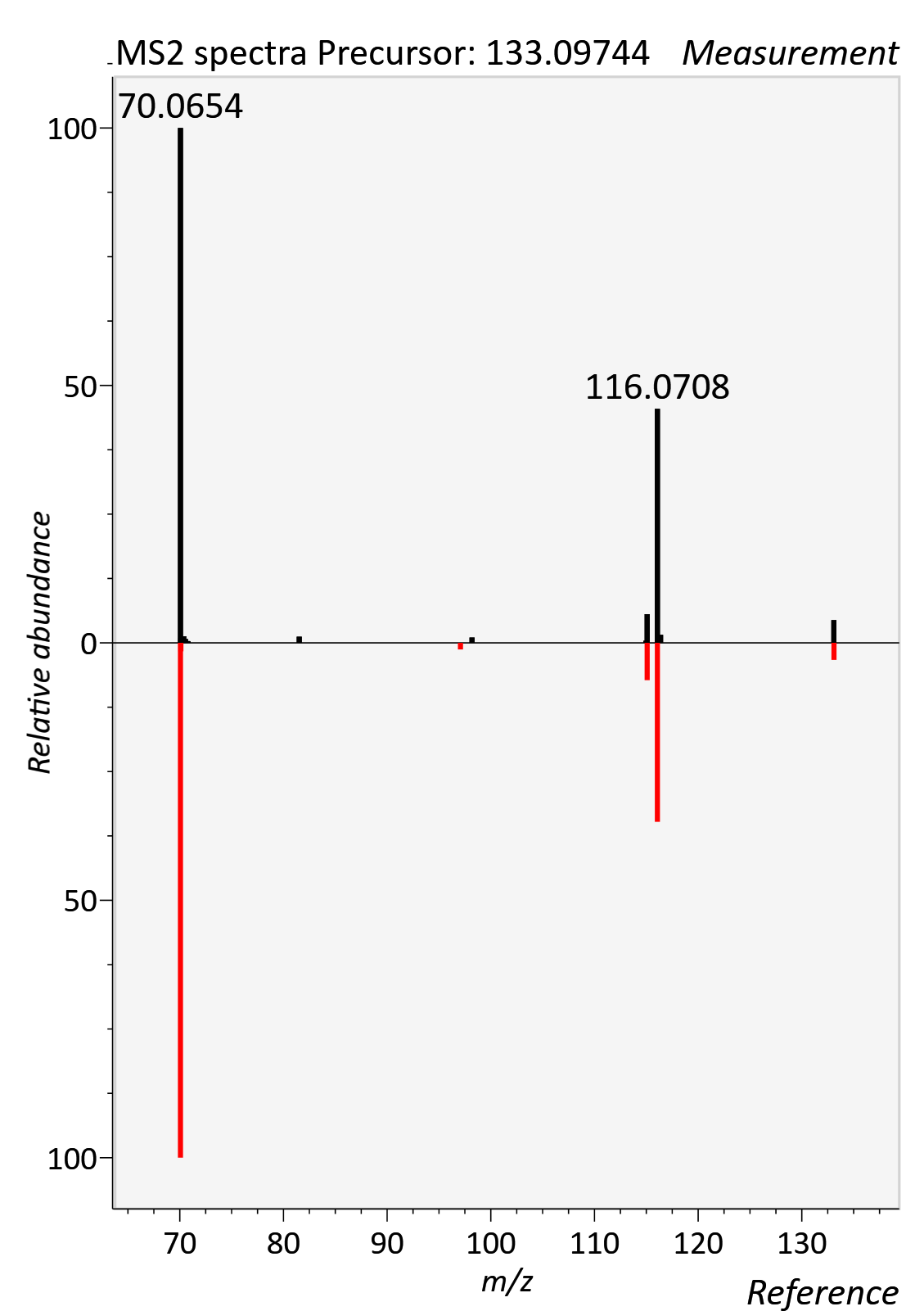

Paraxanthine

RT std: 2.93min, RT experimental: 2.61min, RT Δ 0.32min

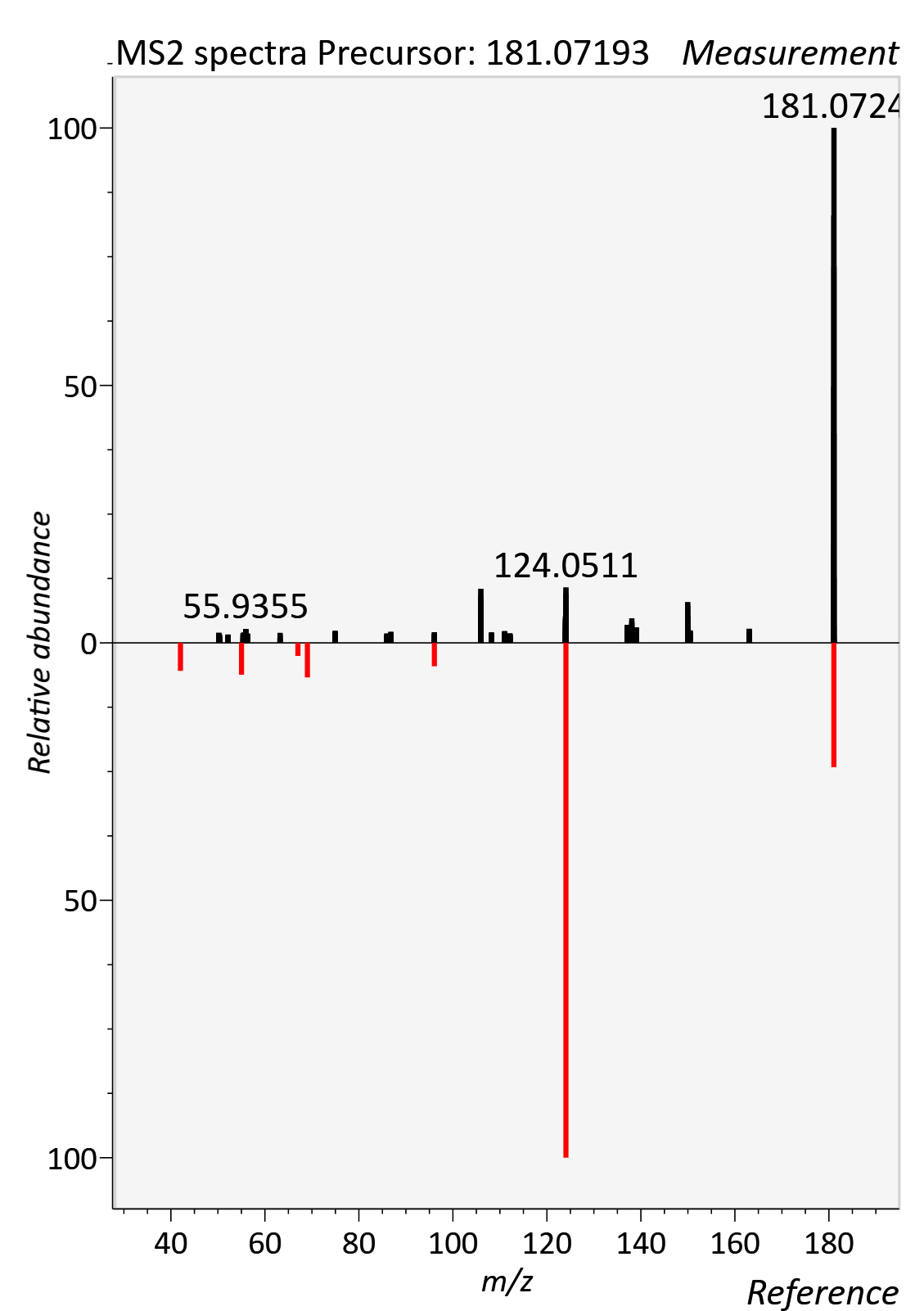

Phenylalanine

RT std: 3.71min, RT experimental: 4.22min, RT Δ 0.51min

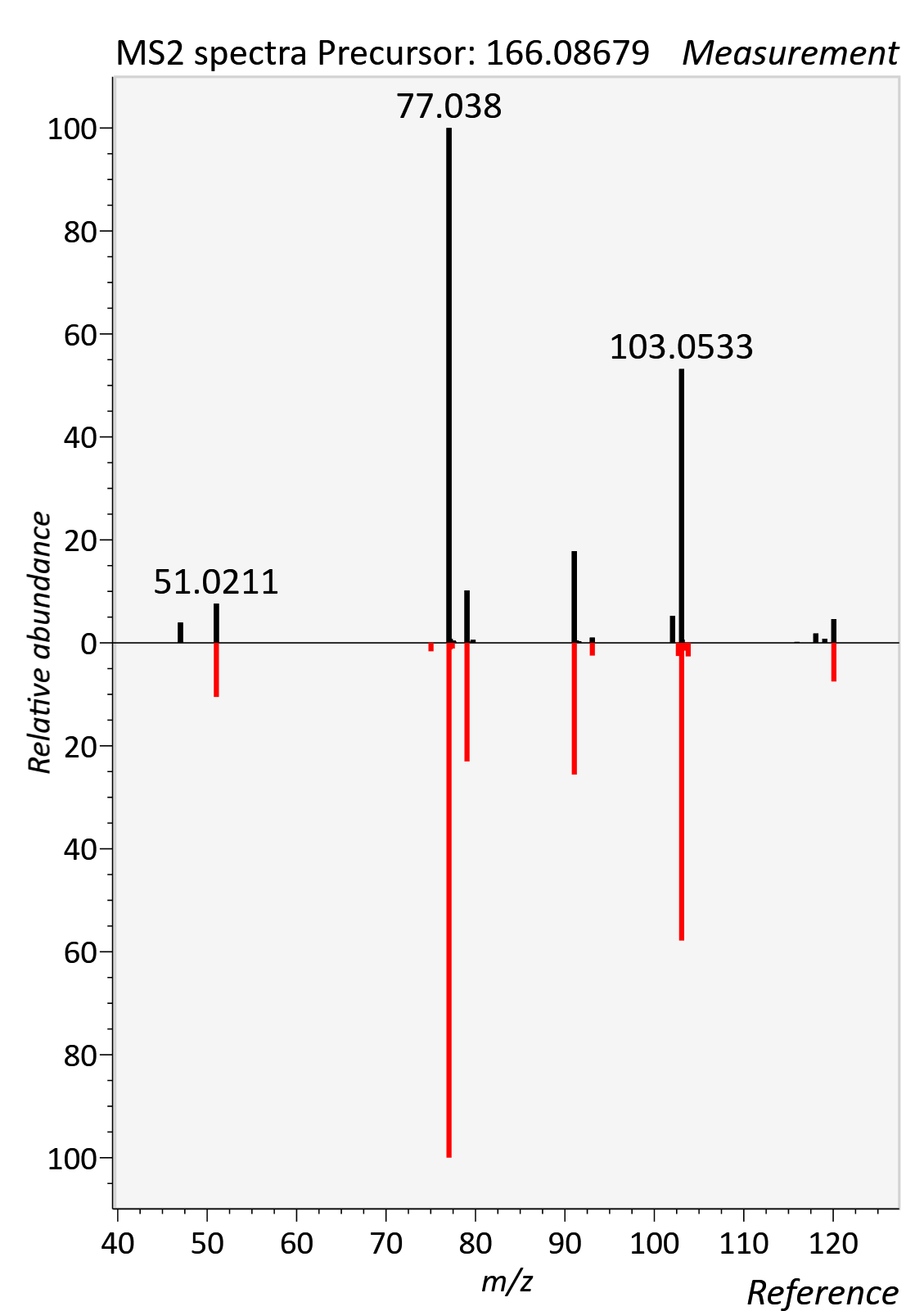

Pipecolic acid

RT std: 4.96min, RT experimental: 4.19min, RT Δ 0.75min

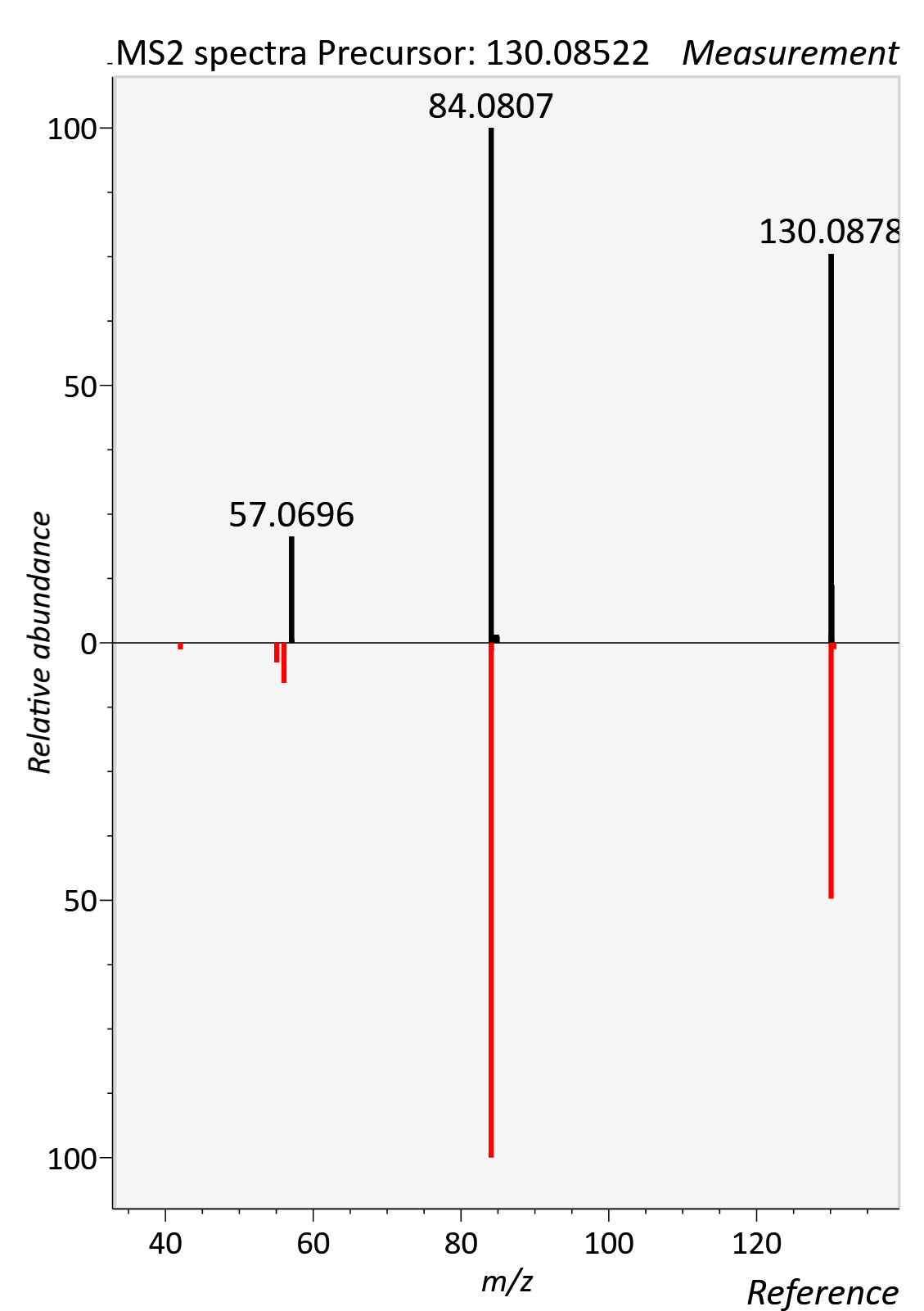

Proline Betaine

RT std: 3.46min, RT experimental: 3.85min, RT Δ 0.39min

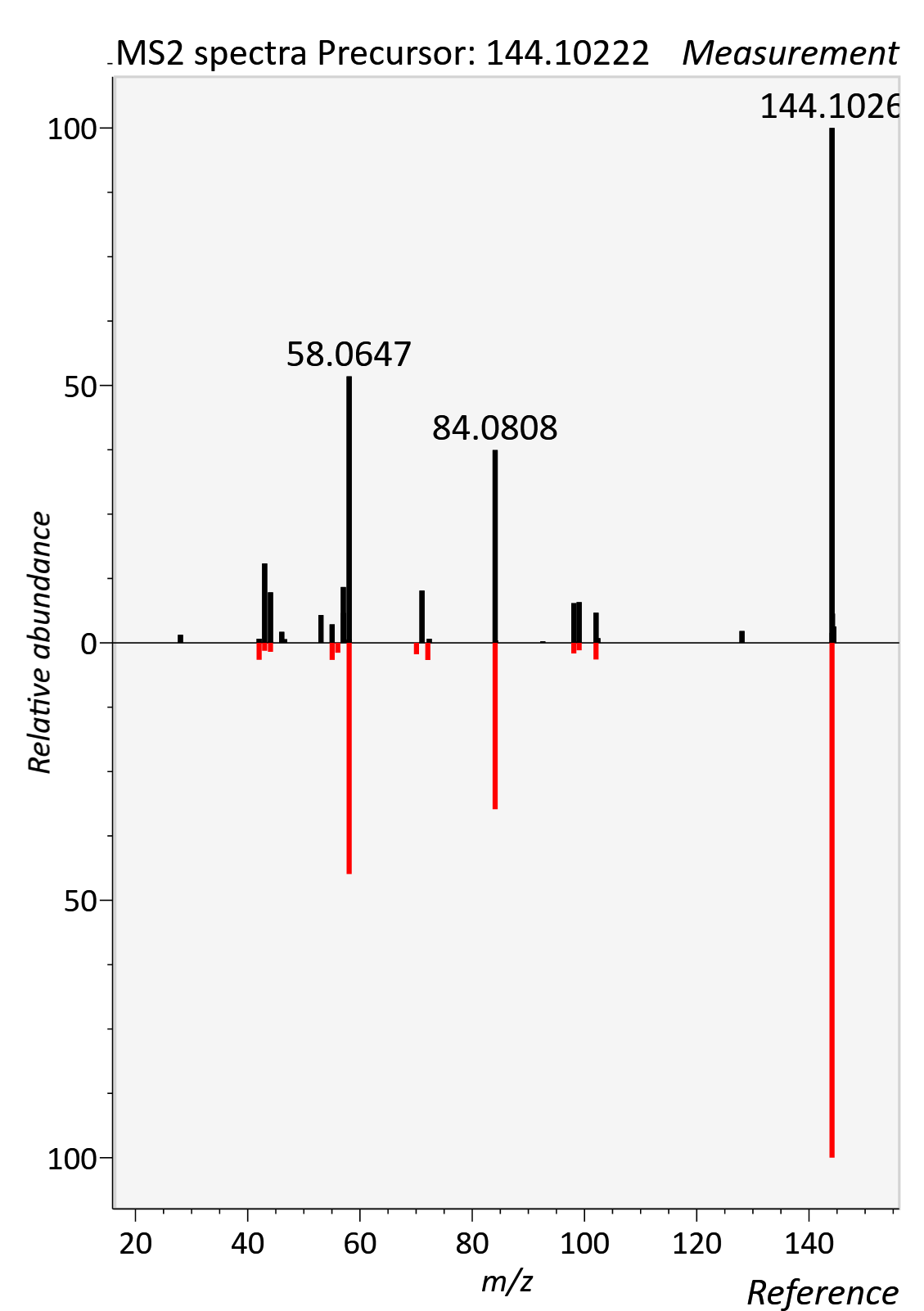

Theobromine

RT std: 2.40min, RT experimental: 2.07min, RT Δ 0.33min

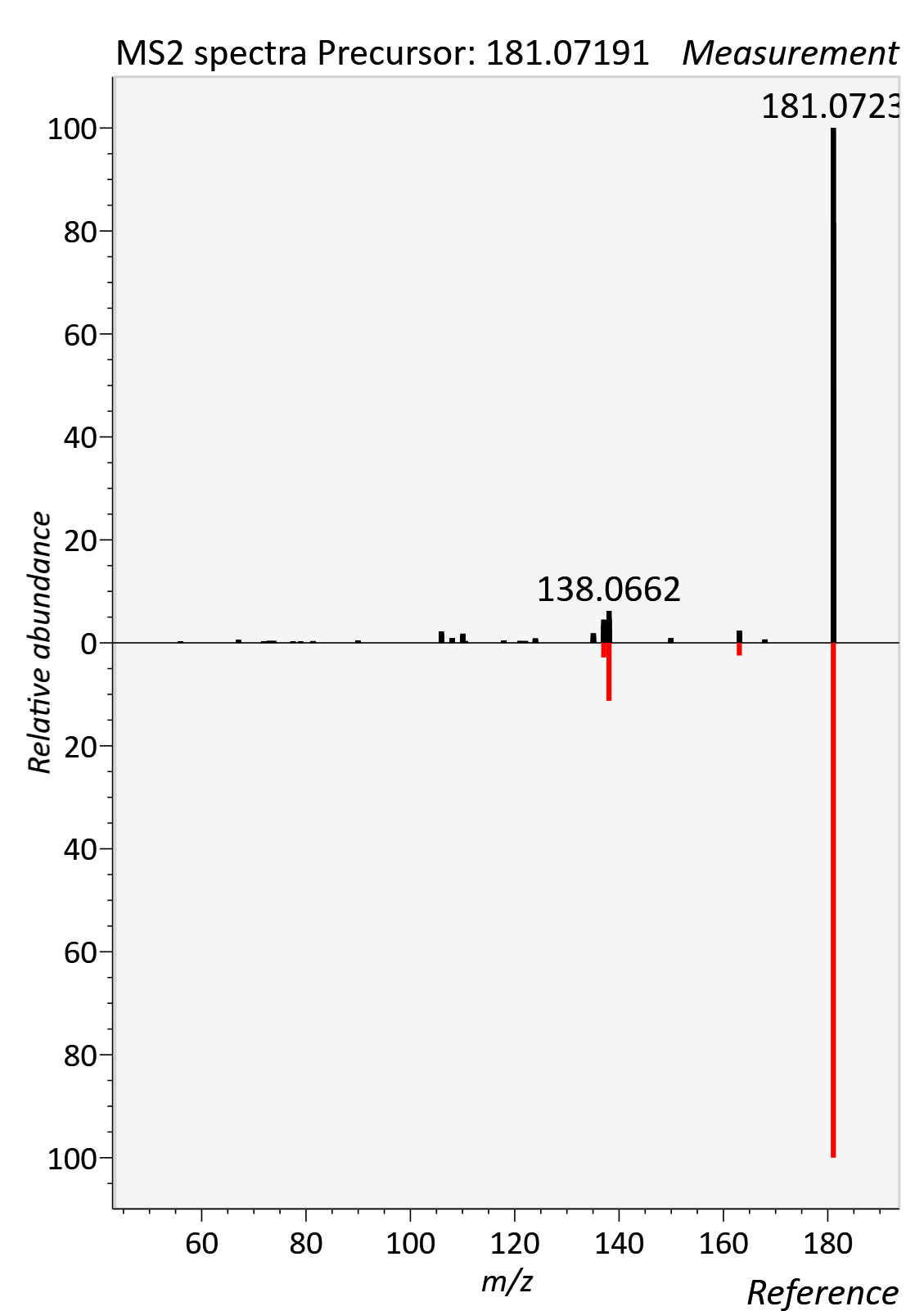

Theophylline

RT std: 2.93min, RT experimental: 3.06min, RT Δ 0.13min

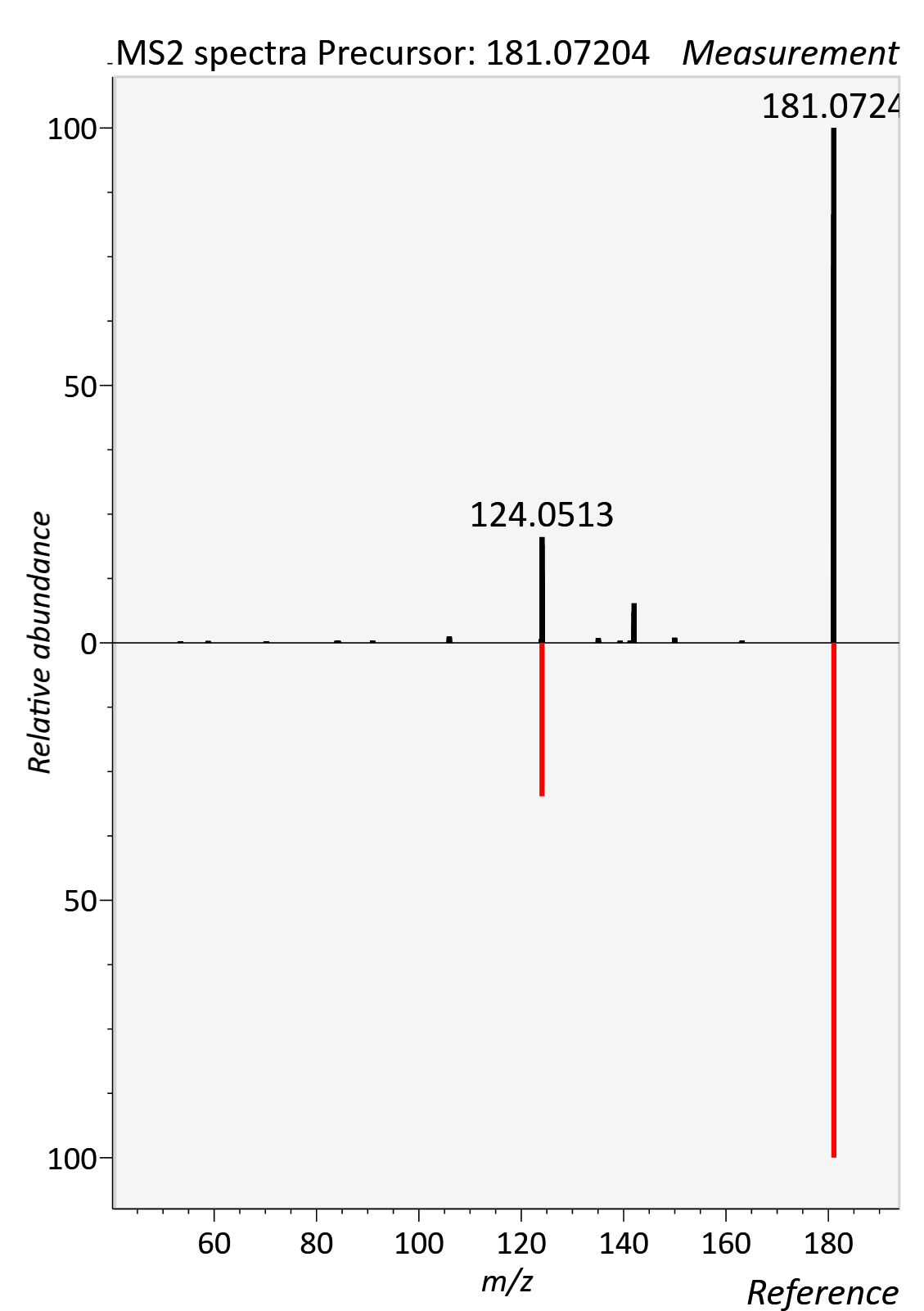

Threonine

RT std: 5.78min, RT experimental: 5.95min, RT Δ 0.17min

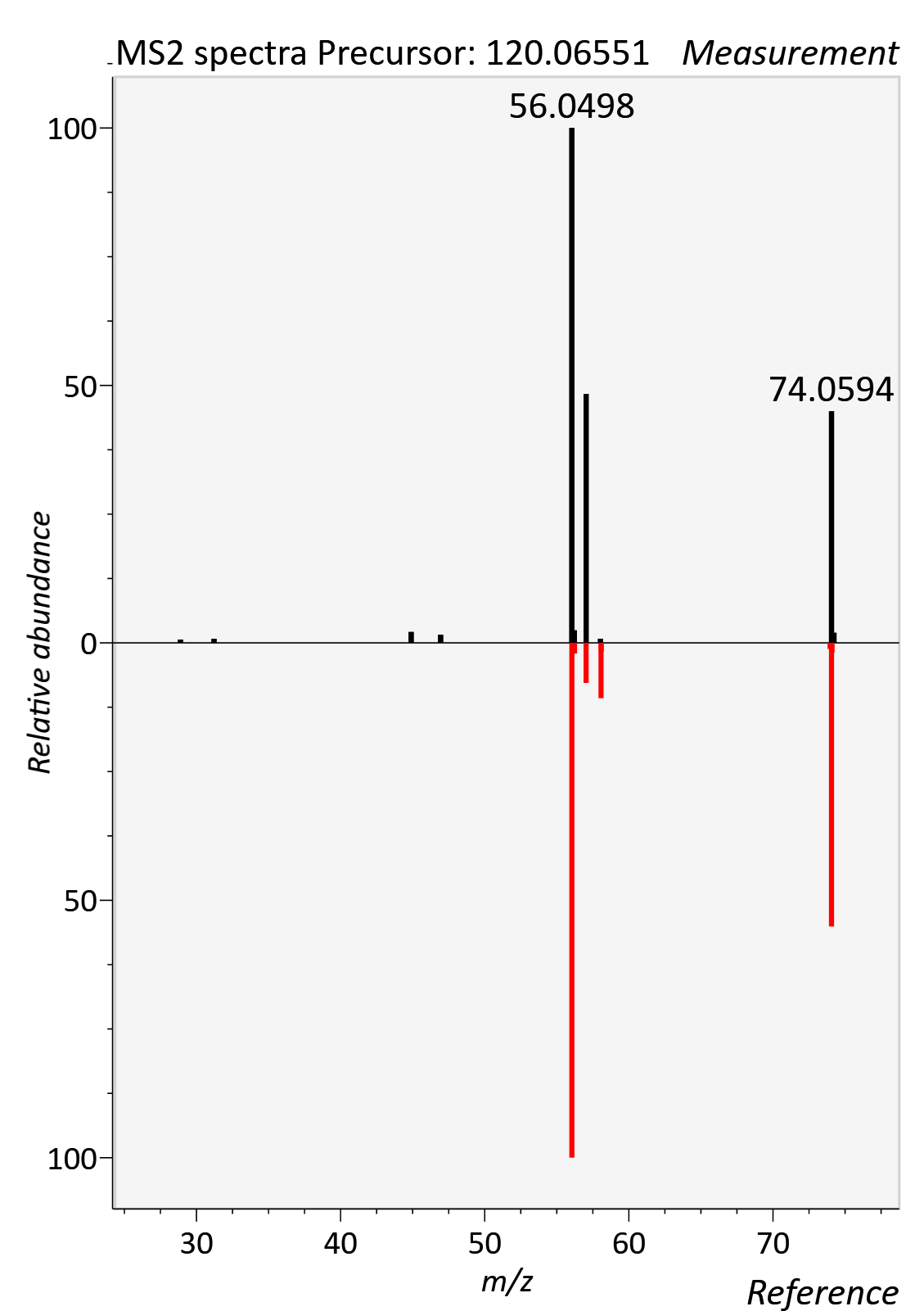

Trigonelline

RT std: 4.03min, RT experimental: 4.52min, RT Δ 0.49min

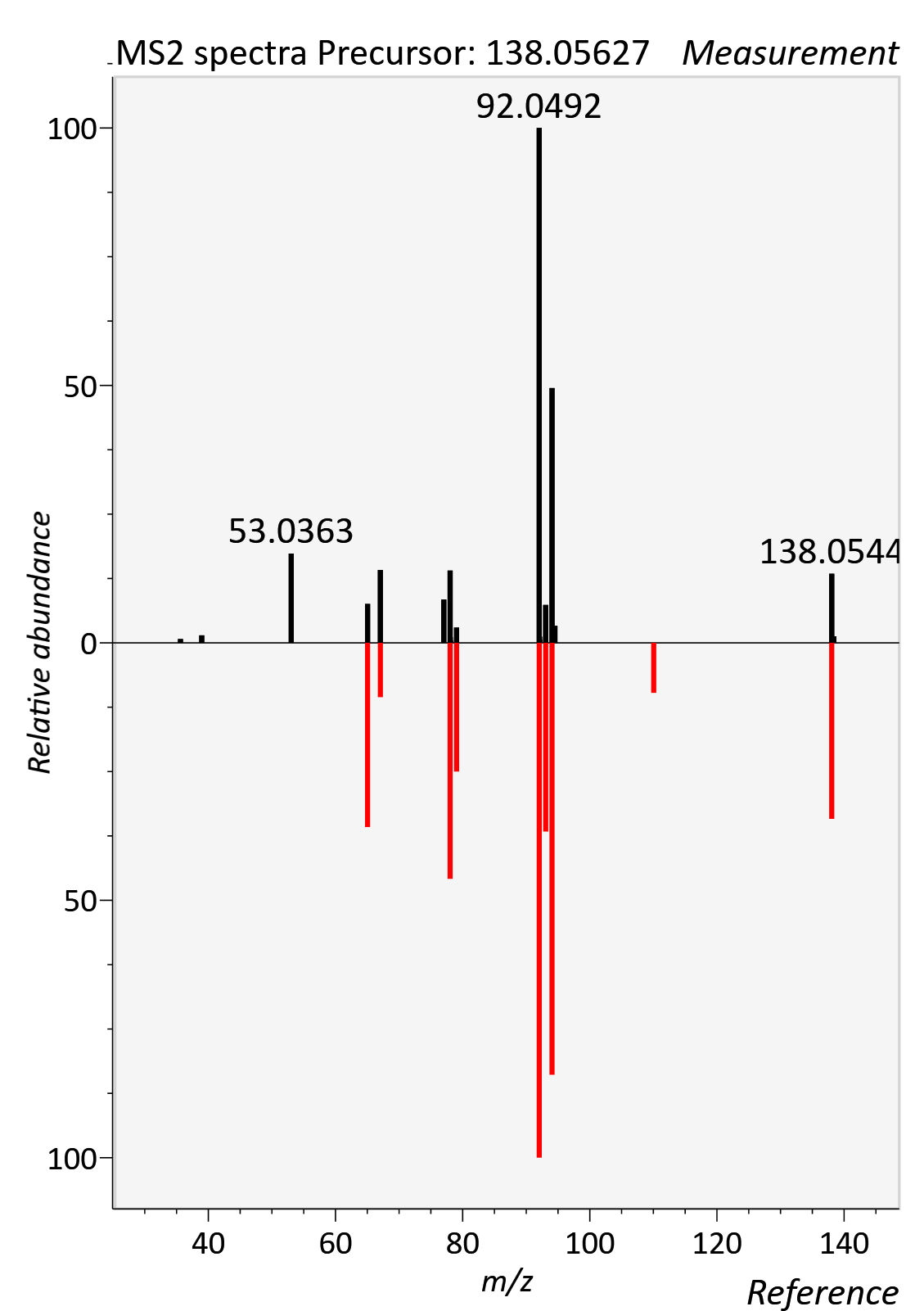

Tryptophan

RT std: 2.76min, RT experimental: 2.43min, RT Δ 0.33min

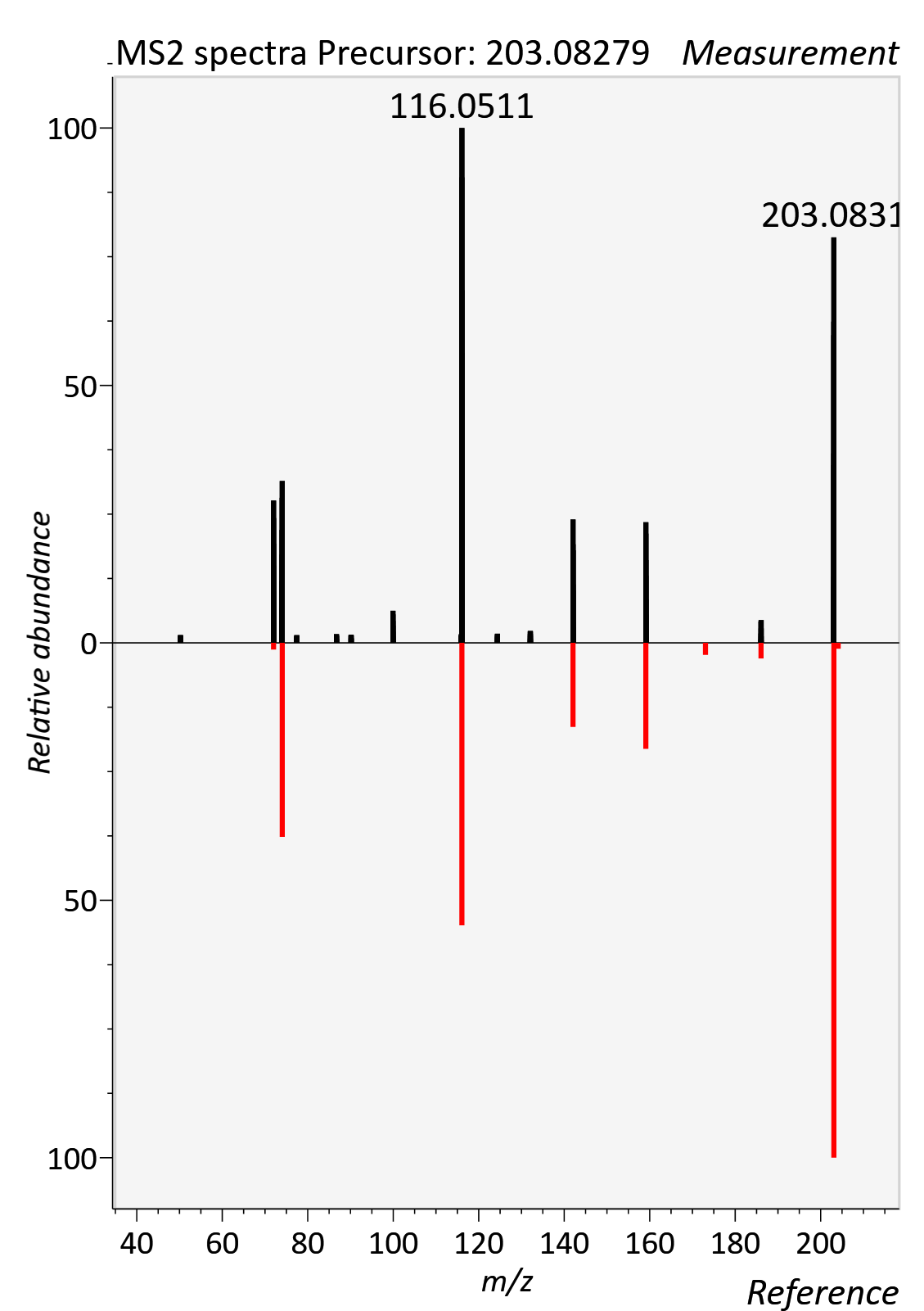

Tyrosine

RT std: 5.01min, RT experimental: 5.30min, RT Δ 0.29min

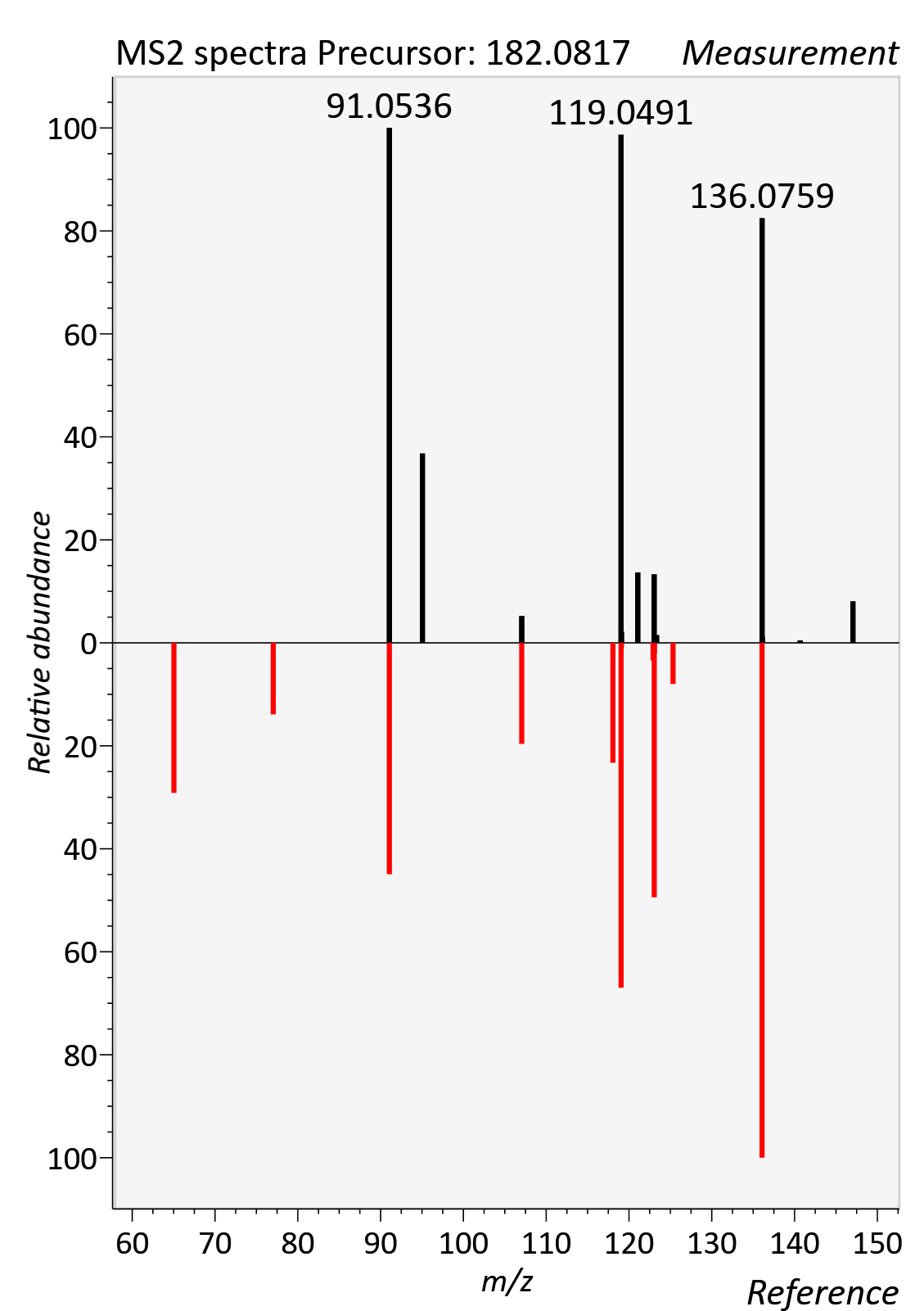
